# Predictive Biophysical Neural Network Modeling of a Compendium of *in vivo* Transcription Factor DNA Binding Profiles for *Escherichia coli*

**DOI:** 10.1101/2024.05.23.594371

**Authors:** Patrick Lally, Laura Gómez-Romero, Víctor H. Tierrafría, Patricia Aquino, Claire Rioualen, Xiaoman Zhang, Sunyoung Kim, Gabriele Baniulyte, Jonathan Plitnick, Carol Smith, Mohan Babu, Julio Collado-Vides, Joseph T. Wade, James E. Galagan

**Affiliations:** Department of Biomedical Engineering, Boston University, 44 Cummington Mall, Boston, MA 02215; Instituto Nacional de Medicina Genómica, Periférico Sur 4809, Arenal Tepepan, Ciudad de México 14610, México; Escuela de Medicina y Ciencias de la Salud, Tecnológico de Monterrey, Ciudad de México, México; Centro de Ciencias Genómicas, Universidad Nacional Autónoma de México, Avenida Universidad s/n, Cuernavaca 62210, Morelos, México; Department of Biochemistry, University of Regina, Regina, Saskatchewan, SK S4S 0A2, Canada; Wadsworth Center, New York State Department of Health, Albany, NY, USA; Centre for Genomic Regulation (CRG), The Barcelona Institute of Science and Technology, Dr. Aiguader 88, Barcelona 08003, Universitat Pompeu Fabra (UPF), Barcelona, Spain; Department of Biomedical Sciences, University at Albany, SUNY, Albany, NY, USA; Bioinformatics Program, Boston University, 24 Cummington Mall, Boston, MA 02215

## Abstract

The DNA binding of most *Escherichia coli* Transcription Factors (TFs) has not been comprehensively mapped, and few have models that can quantitatively predict binding affinity. We report the global mapping of *in vivo* DNA binding for 139 *E. coli* TFs using ChIP-Seq. We used these data to train BoltzNet, a novel neural network that predicts TF binding energy from DNA sequence. BoltzNet mirrors a quantitative biophysical model and provides directly interpretable predictions genome-wide at nucleotide resolution. We used BoltzNet to quantitatively design novel binding sites, which we validated with biophysical experiments on purified protein. We have generated models for 125 TFs that provide insight into global features of TF binding, including clustering of sites, the role of accessory bases, the relevance of weak sites, and the background affinity of the genome. Our paper provides new paradigms for studying TF-DNA binding and for the development of biophysically motivated neural networks.

## Introduction

*Escherichia coli* is the most widely used cell in biology and biotechnology, the best studied model prokaryote^1,2^, and the foundation for most efforts in synthetic biology. Yet despite its central importance, and a wealth of knowledge, surprising gaps remain^3^. Bacteria typically encode hundreds of transcription factors (TFs) whose binding to DNA modulates gene regulation. Decades of work in *E. coli* have led to a deep mechanistic understanding of TF function. Yet this high level of knowledge is limited to a handful of well-studied TFs. The binding affinities of most of the ∼300 TFs in *E. coli* have not been comprehensively mapped; indeed a recent publication estimates that only 30% of *E. coli* TF-DNA interactions have been identified^4^. Moreover, many TFs lack information about even a single sequence to which they bind. And very few have experimentally validated biophysical models that can be used to quantitatively understand TF binding behavior. Such an understanding is crucial to fully deciphering cellular function and predictively engineering synthetic biology circuits.

Chromatin-immunoprecipitation followed by sequencing (ChIP-Seq) enables the genome-wide characterization of TF binding under *in vivo* conditions. However, differences in protocols, conditions, and computational processing can modulate the degree of real or apparent binding, complicating the comparison and interpretation of these results. Applied to the smaller genomes of bacteria, ChIP-Seq also has the capability to detect binding over a wide range of apparent affinities, down to very weakly bound sequences^5-8^. Commonly, weakly bound sites are often discarded^9^ despite being highly reproducible^5,7^. And recently, weak affinity TF binding has been demonstrated to have unique regulatory function^10,11^. Typically, ChIP-Seq analyses focus on cataloging a discrete set of “true” high affinity sites. A complete model of TF binding, however, must predict affinity for any sequence.

In recent years, convolutional neural networks (CNNs) for analyzing ChIP-Seq, and related, data have been developed with promising results^12-14^. These models leverage the feature discovery capabilities of CNNs and the flexibility and optimization capabilities of advanced neural network frameworks. Developed for eukaryotes, existing models borrow the template of image processing CNNs, with many kernels and many convolutional layers using standard neural network activation functions. These deep architectures were motivated by the recognition that multiple factors, including chromatin context and co-factors, mediate the binding of TFs in eukaryotes. However, it is not clear the degree to which these complex factors are required to understand TF binding to DNA in bacteria^15^. Moreover, deep CNNs come at a cost of greater complexity, which can lead to overfitting and necessitates additional algorithms for interpretation. Significantly, the scores derived from such interpretations are not directly tied to any physical parameters that can be experimentally verified.

Thermodynamic models provide a predictive description of TF function tied to biophysical quantities that can be experimentally validated^16-19^. TF binding is characterized by the energy released, Δε, when a protein binds a DNA sequence. Under the assumption of thermal equilibrium, Δε can be used to predict the probability of binding, and corresponding dissociation (or association) constants via the Boltzmann distribution. Multiple studies have demonstrated that this framework can be used to accurately predict the sequence specificity of TF binding and downstream gene regulation^20^. This framework has been applied to a range of high-throughput TF binding assays (Spec-Seq, MITOMI, PBMs, SELEX). Most recently, a machine learning method, ProBound, was developed for predicting biophysical parameters from SELEX experiments that could be applied to ChIP-Seq data^21^. However, ProBound was designed for eukaryotes, and is based on a custom machine learning framework, making it more difficult to adapt to other uses.

In this study, we report the global mapping of DNA binding for 139 *E. coli* TFs under controlled *in vivo* conditions using a standardized ChIP-Seq and computational analysis protocol. Our data provides the most comprehensive view of *E. coli* TF binding to date, and a foundation to study the determinants of TF-DNA interactions in all bacteria. We have developed a novel neural network, “BoltzNet”, that accurately predicts ChIP coverage and TF binding affinity from DNA sequence. In contrast to previous neural networks, BoltzNet is based on an explicit quantitative biophysical model of TF-DNA binding and provides directly interpretable physical predictions genome-wide at nucleotide resolution. We use BoltzNet to quantitatively design novel binding sites, which we then experimentally validate using independent *in vivo*, and *in vitro* biophysical binding assays. Our results confirm that BoltzNet directly predicts a highly accurate model of relative binding energies for existing and novel binding sites. They also provide insight into several global features of TF binding behavior, including clustering of binding sites, the importance of poorly conserved accessory bases, the physiological relevance of weak binding sites, and the background affinity of the genome. We have generated high-confidence models for 125 TFs that can be used and extended by any researcher to quantitatively interpret TF binding behavior and to predictively engineer new binding sites.

## Results

### Large-scale mapping of TF binding sites in *E. coli*

We developed a standardized protocol for mapping *E. coli* TFs with ChIP-Seq (Figure S2). TFs were tagged for efficient IP and expressed in two ways: from their native genomic promoters and loci, or inducibly expressed from a plasmid. We used both systems to maximize the likelihood of adequate TF expression. We applied both methods to each TF and required at least two successful experiments with either method for a TF to be considered mapped. All ChIP-Seq was performed *in vivo* under a single controlled condition. In addition, for selected TFs, we also performed ChIP-Seq *in vitro* with controlled protein concentrations. All results were analyzed using a standardized pipeline that filters out common artifacts and identifies enriched regions (Figure S4). Enriched regions are defined as contiguous sequences longer than 150 bp with coverage that is statistically enriched compared to background and that show the expected signature of forward and reverse coverage for protein binding (Figure S5). Enriched regions may contain multiple binding sites^22^. Coverage for enriched regions is normalized to the genome mean.

We generated high-confidence global mapping data for 139 TFs (Figure 1A, Figure S10). Enriched regions included the vast majority of reported high-confidence binding sites in RegulonDB for these TFs as well as thousands of additional reproducible binding regions (Table S1). Novel regions were found for TFs that lack any reported sites, as well as for TFs that have been previously studied. Native and inducible expression were complementary. No single approach was successful for all TFs (Figure 1B) and both were required to recover known sites (Figure 1C), even for TFs with data from both approaches (Figure 1D), although the inducible system outperformed native expression. Our results show enrichment is highly reproducible across replicates of the same expression mode, and highly correlated between different expression modes (Figure 1E,F).

**Figure 1:**
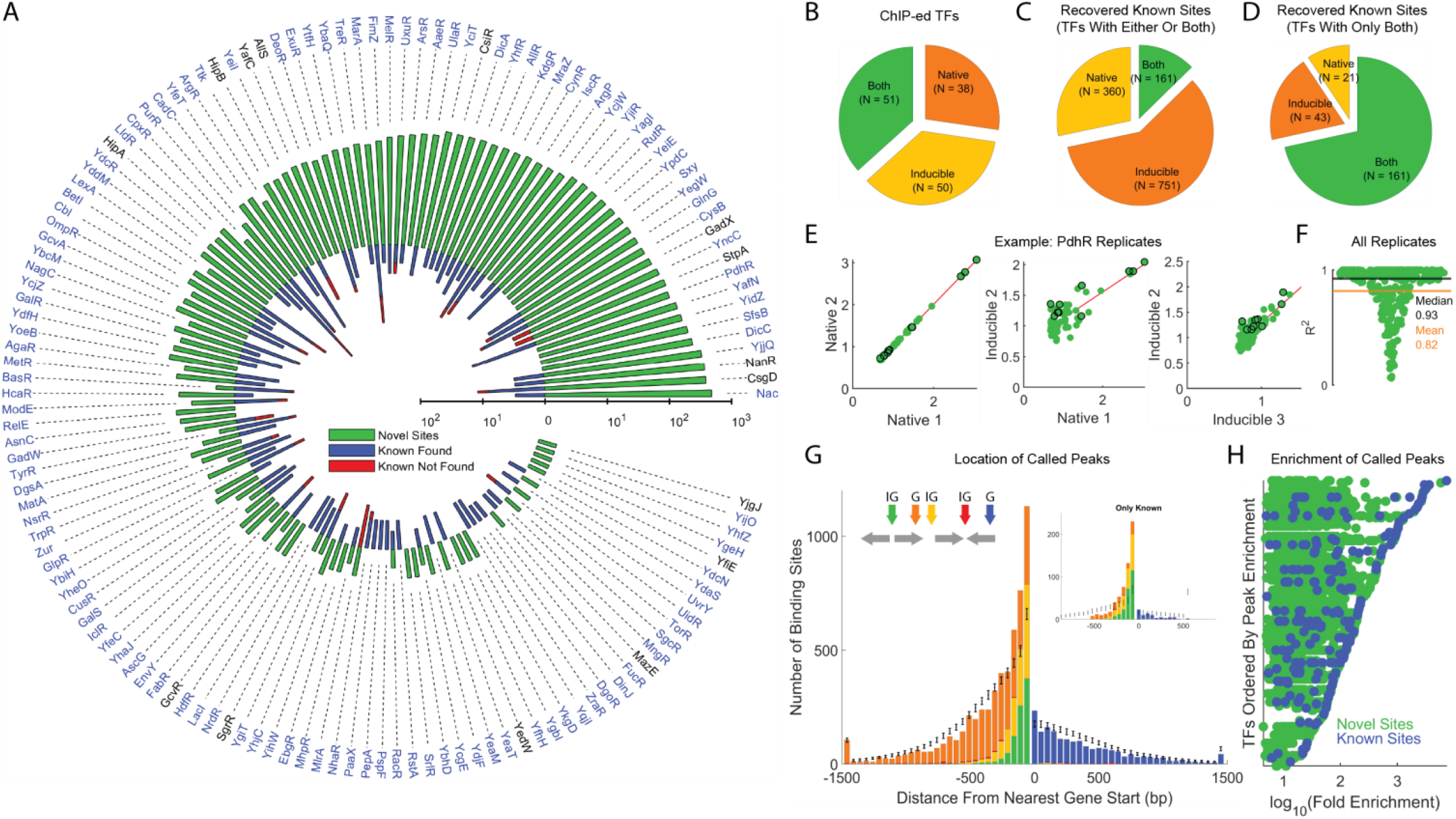
Large-scale mapping of *E. coli* TF binding regions. **A)** Summary of novel regions (green) and known regions (found:blue, not found:red) for 139 mapped TFs, ordered by total number of binding regions (TFs in blue have BoltzNet models). **B)** Mode of TF expression used to map TFs. Both experimental approaches required to map all TFs. Recovery of known sites by different expression modes for TFs mapped by **C)** either or both modes of expression, or **D)** both modes only. No single approach recovers all known sites, although TF induction performs better. **E)** Example for PdhR of comparisons of ChIP-Seq experiments within and between expression modes. **F)** Correlation of enrichment across all replicates for all TFs. **G)**. Location of binding regions relative to start position of nearest gene (G – genic, IG – intergenic). Binding regions between 150 bp upstream and 50 bp downstream are overrepressented relative to random expectation (grey symbols), regions > 1.5 kb up or down-stream of the nearest gene are grouped into a single bar on either edge. Data for regions with known sites shown in inset. **H)** Both novel and known regions are observed over 3 logs of enrichment. Each row is a TF and each dot is a called region (green new region, blue known region).

Binding regions were highly enriched within 150 bp upstream and 50 bp downstream of gene starts (Figure 1G). We also see extensive binding within genes, although intergenic binding is ∼2.5-fold overrepresented (28% of enriched regions vs ∼10.96% intergenic genome). Conversely, binding further from gene starts is not markedly different from expectation for all ChIP regions but is underrepresented in the set of high-confidence known sites^23^. We observe over three logs of enrichment across all binding regions, and within regions for individual TFs (Figure 1H, green). Known sites are skewed to regions of higher enrichment but span the range of absolute and relative enrichment (Figure 1H, blue). The number of binding regions varies dramatically across TFs (Figure 1A, F) and can be approximated by a power law (p(k)∼k^-1.9^)^24,25^. We observe several TFs that reproducibly bind throughout the genome. Conversely, 23 TFs have only a single binding region (Table S1) and of these 17 (74%) are regions just upstream of the TF operon that imply autoregulation. Of all 139 TFs reported, 95 (68%) have autobinding, consistent with autoregulation as a highly enriched regulatory motif^26^. Genome-wide, we observe that most regions are bound by at most one TF, while certain regions can be bound by multiple TFs (Figure S14). This large diversity of binding regions provided a unique opportunity to study and model the global binding preferences of bacterial TFs.

### BoltzNet architecture and training

To model sequence affinity and ChIP-Seq coverage as a function of sequence, we developed a convolutional neural network (CNN), BoltzNet, specifically designed to mirror a two-stage quantitative biophysical model of TF-binding and ChIP-seq. The first stage consists of a thermodynamic model of TF binding to a DNA site driven by the binding energy of the site relative to an unbound state, Δε^16^. Several studies have demonstrated that simple additive “energy matrix models” are often sufficient to describe TF-DNA binding energies^17-19,27,28^. Assuming TF binding is in thermal equilibrium, the probability that the site will be bound can be calculated via the Boltzmann distribution as:

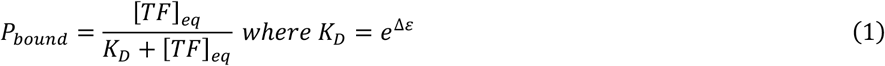

where K_D_ is the equilibrium disassociation constant of the binding reaction. Crucially, both K_D_ and the probability of binding depend on the exponentiation of Δε. Given a sequence with multiple sites, the probability of a protein being bound to the sequence is related to the sum of the exp(Δε) for all sites (see Methods). In the second stage, ChIP-Seq coverage is a function of the probability that a site is bound, which we have shown can be modeled as a signal convolution^22,29^.

BoltzNet is designed with two components to mirror these two stages (Figure 2A). The first component inputs 101 bp DNA sequences and models the affinity at every position in both orientations using a convolution layer consisting of a single 25 bp weight matrix (kernel) and bias term. The use of a single kernel encodes the hypothesis that a single linear energy matrix can predict binding energy. The output of the kernel feeds into an exponential activation function, so that the convolution layer models the affinity for each site. Subsequent pooling layers sum these values to generate an affinity score for the entire 101 bp sequence. The second component predicts coverage in multiple ChIP-Seq experiments as a function of the affinity score. We learn this mapping using the universal function approximation capabilities of a fully connected feedforward neural network^30^. BoltzNet is trained on a set of sequences consisting of all enriched regions for a TF (positive samples) as well as 5000 randomly selected genomic regions (negative samples) (see Methods for details).

**Figure 2:**
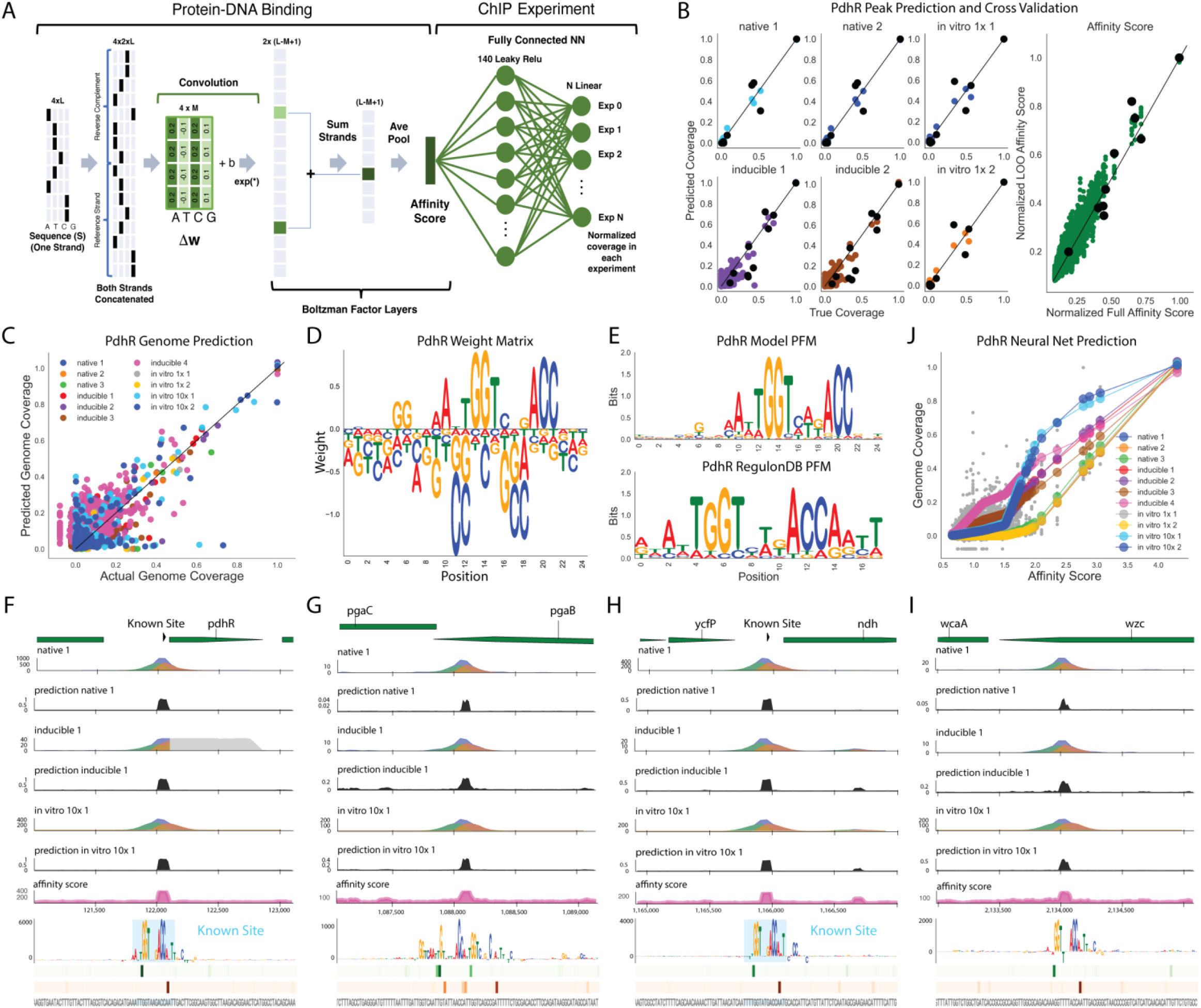
BoltzNet accurately predicts binding site locations, motifs, and coverages. Detailed example for PdhR. **A)** BoltzNet architecture. BoltzNet mirrors a thermodynamic model of TF-DNA Binding to predict ChIP-Seq coverage from sequence. A convolution component models the effective energy of TF binding to the sequence as an affinity score. The fully connected neural network component models normalized ChIP coverage in multiple experiments as a function of this score. **B)** Prediction of coverage and cross-validation on training set. Predictions on a subset of representative experiments (full set in Figure S12). Black circles show leave-one-out cross-validation predictions. All coverage is normalized between zero (baseline) and 1 (most enrichment). Native=native promoter replicate number, inducible= inducible promoter replicate number, in vitro #x = *in vitro* ChIP-Seq replicate numbers at two different protein concentration. **C)** Accuracy of genome-wide prediction of normalized coverage in all experiments. **D)** A single energy weight matrix is sufficient for accurate prediction. Positive values represent more energetically favorable bases. **E)** Predicted PFM matches known PFM from RegulonDB^1^. **F-I)** BoltzNet provides direct interpretation at all scales. Four representative regions are shown. Top track is gene annotation for region. Next tracks show true coverage (red, green, blue), genome-wide coverage predictions (black) for three representative experiments, and the predicted affinity score (pink). Coverage in units of fold enrichment over mean coverage. Bottom track shows single nucleotide resolution predictions around binding site. Sequence logo shows base contribution score. Cyan shading shows known binding sites. Heatmaps shows predicted affinity at each position in positive (green) and negative (red) orientation. Sequence shown in gray. **J)** Neural network predicts experiment specific coverage from a single sequence affinity score. Experiments with higher known or predicted protein concentrations are correctly predicted to have higher coverage for the same predicted affinity. Large symbols are predicted coverage in each experiment and small gray symbols are actual normalized coverage.

### BoltzNet accurately models sequence binding strength

Figure 2B-F provides an analysis of a BoltzNet accuracy for PdhR. PdhR is a GntR family TF with 10 validated known binding sites and a known HTH binding motif. We performed 11 ChIP-Seq experiments on PdhR including replicates of *in vivo* ChIP-Seq with both native and inducible promoters. Additionally, we performed *in vitro* ChIP-Seq at two different protein concentrations. Across all experiments we call 204 enriched regions spanning three logs of enrichment.

BoltzNet accurately predicts enrichment in all experiments (Figure 2B) and demonstrates high specificity when applied to all 101 bp sequences in the genome (Figure 2C). As an initial assessment of generalizability, we performed leave-one-out (LOO) cross-validation on the top 10 enriched regions. For each left-out sample both the output coverages and relative affinity scores were correctly predicted (Figure 2B and Figures S15 & S16). And when the most enriched region was left out, it was correctly predicted in all experiments, demonstrating the potential for BoltzNet to extrapolate.

### BoltzNet is interpretable and verifiable genome-wide at nucleotide resolution

The accuracy of BoltzNet is achieved from a single weight matrix that directly represents the relative contributions of each base at every position of a 25bp binding site (Figure 2D). The weight matrix differs from motif logos based on position frequency matrixes (PFMs), which are the most common means of representing binding sites. However, PFMs model the frequency of bases at each position in a collection of sites but not directly the contribution of each base to binding strength. Moreover, PFMs are not capable of predicting ChIP-Seq coverage (Figure S18). PFMs can be derived from weight matrixes by scanning matrixes over a set of sequences and counting the bases in each position scoring above a threshold^31^. Applying this approach, we confirm that BoltzNet identifies the known binding motif for PdhR (Figure 2E).

On a genome-wide scale, the model demonstrates spatial accuracy in predicting coverage in multiple experiments simultaneously, all derived from a single affinity score for each 101 bp sequence (Figure 2F-I). Within each sequence region, the output of the convolution provides a measure of the affinity at every position and orientation (heatmap). And at nucleotide resolution, we can measure the contribution of each base by summing the base weight in all overlapping sites multiplied by the affinity score of that site. This can be summarized in a sequence logo (Figure 2F-I). A comparison of nucleotide level predictions with the location of know binding sites (cyan shaded regions) demonstrates that BoltzNet has accuracy to exact binding locations.

### BoltzNet models expected behavior of different ChIP experiments

The neural network component must learn a mapping from a single affinity score to coverage in multiple experiments accounting for variations in IP, cell state, growth conditions, and especially protein concentration. As protein levels increase, stronger sites are bound first and saturate as weaker sites are bound. Figure 2J demonstrates that BoltzNet learns this behavior. For *in vitro* experiments, the model accurately predicts higher and more saturating coverage with 10x protein compared to 1x (Figure 2J). Similarly, comparing *in vivo* experiments show increasing coverage and saturation with the expected greater protein concentration from inducible promoters relative to native promoters. This verification of the neural network component provides a consistency check on the model as a whole.

### A compendium of TF binding models

Applied to *in vivo* ChIP-Seq data for 139 *E. coli* TFs, we have generated BoltzNet models for 125 that passed criteria for accuracy in predicting coverage and for specificity on the whole genome (Table S2). For 36 models that overlap with known motifs in RegulonDB, we see strong concordance with the model-predicted PFM (Figure 3A). Figure 3B-E illustrates the range of different TF behavior captured by BoltzNet models. TFs with a range of binding site profiles can be modeled. UlaR illustrates a model based on one strong site and multiple weak sites while Nac (as we have previously reported^5^) and GlnG have many called sites spanning the range of relative coverages. And TFs with only one called site can be modeled with the same high accuracy in identifying known sites (Figure S19). Binding sites of both low (GlnG) and high (Nac) AT-content are accurately modeled. TFs associated with σ^70^ (PdhR, AllR, Nac) and σ^54^ (GlnG) are equally well modeled.

**Figure 3.**
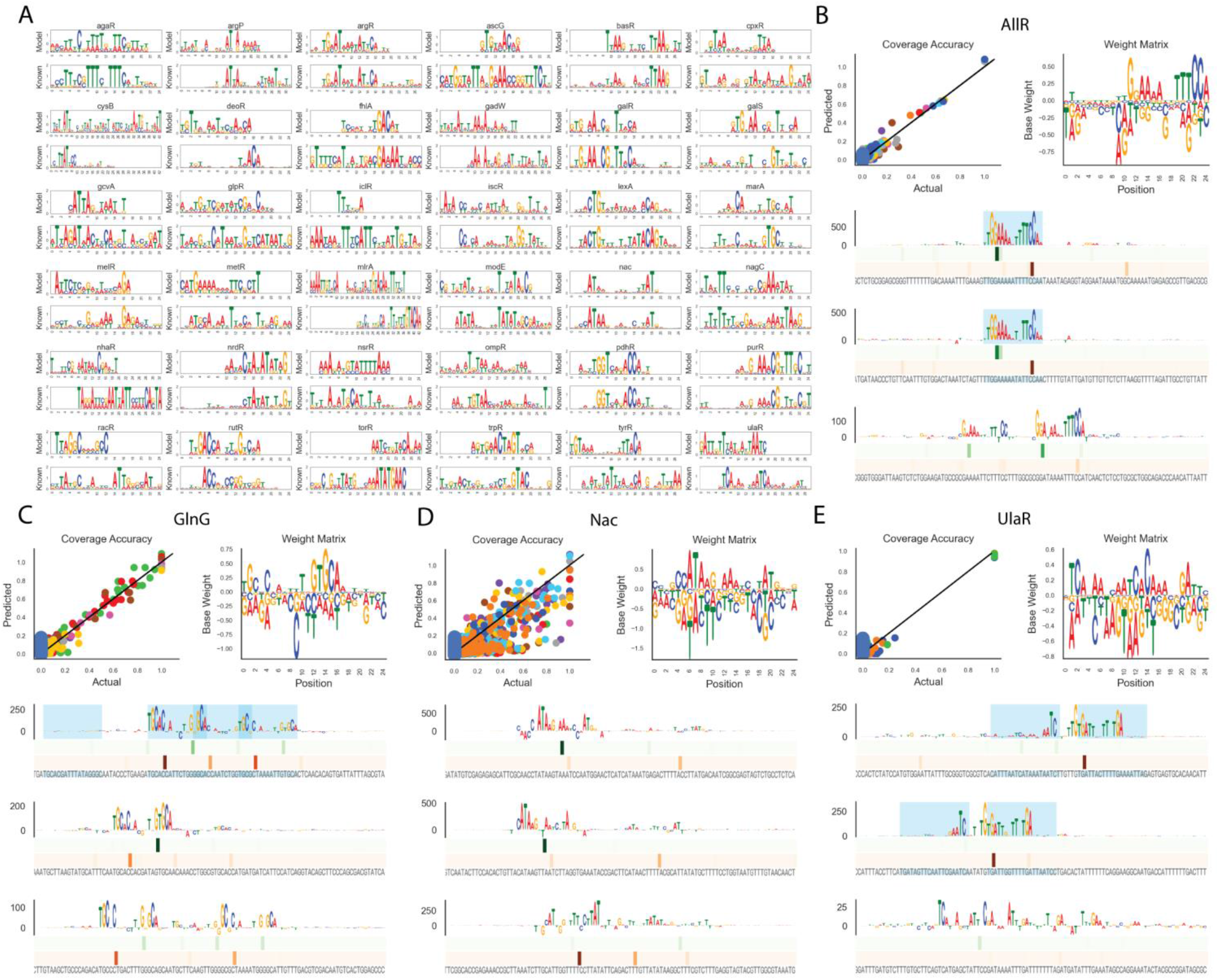
A compendium of predictive models. We have generated models for 125 TFs (Table S2). **A)** Alignments of motifs for all models with known RegulonDB motifs. **B-E)** Summary of models for AllR, GlnG, Nac, and UlaR. Top row of each panel displays a scatter plot of region coverage prediction accuracy (left) and the weight matrix (right). The bottom half of each panel shows nucleotide level resolution predictions as in Figure 2G for top 3 binding sequences for each TF ordered from strongest (top) to weakest (bottom).

### Role of clustered binding sites and accessory bases

An examination of nucleotide level predictions across models reveals two themes predicted to contribute to binding strength. First, many sequences contain multiple predicted non-overlapping binding sites (e.g. Nac, AllR, GlnG). Clustered sites (e.g. Figure 2G) can result in affinity scores equal to (e.g. Figure 2I) or stronger than any single site (e.g. Figure 3C, top sequence), suggesting that similar sequence occupancies can be achieved both ways. Moreover, BoltzNet sums the contributions of clustered sites, the accuracy of which suggests that cooperative interactions do not substantially impact occupancy^32^. Second, core bases are commonly conserved across a range of binding site strengths. For TFs that bind as a dimer this core is typically palindromic. Differences in binding strength appear to be determined by accessory bases outside the core, potentially by stabilizing or destabilizing contacts with the core motif. This is also apparent from weight matrices, where core bases have the strongest relative contributions, but bases outside the core can be equally important (e.g. for PdhR and GlnG the presence of cytosines between the two core palindromes has significant negative weight – Figure 2D,Figure 3C).

### Design and validation of novel binding sites by Library-ChIP confirms role of accessory bases

To validate the predictive ability of BoltzNet as well as the role of accessory bases, we used our models to design new binding sites to be tested experimentally. This was accomplished by taking a reference binding sequence and generating all variants outside the core (Figure 4A and Figure S20). For each TF, we selected a reference sequence containing only one strong binding site, for simplicity. BoltzNet models based on only *in vivo* ChIP-Seq data were used to score all variant sequences, and a set of sequences spanning a range of predicted binding affinities was selected for experimental validation.

**Figure 4:**
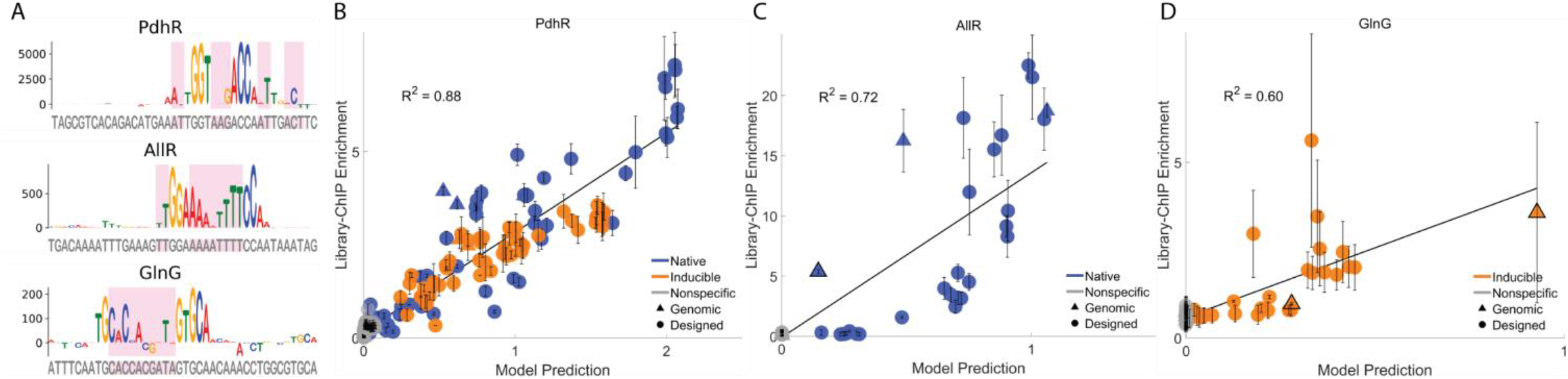
Novel site prediction and validation with Library-ChIP confirms role of accessory bases. **A)** Novel site designs for PdhR, AllR, and GlnG. For each TF, the top singleton binding site was selected as a reference (sequence logo) and all combination of accessory bases (pink shading) were generated and used as input to the corresponding BoltzNet model **B-D)** Scatter plots of predicted BoltzNet predictions vs Library-ChIP enrichment.

We first tested sequences for PdhR, AllR, and GlnG using an independent assay for *in vivo* binding, Library-ChIP^33^ (Figure S21). Library-ChIP enabled high-throughput testing of a large number of sites and tested the ability of BoltzNet to generalize to a different type of binding assay. The results of Figure 4B-D demonstrate that BoltzNet predictions were highly correlated with actual Library-ChIP enrichment.

### BoltzNet accurately predicts binding energy

The variability observed in Figure 4B-D, especially for GlnG, likely reflects the fact that ChIP is as much a function of the extrinsic experimental conditions spanning a population of cells as it is a function of the intrinsic binding energy of a single protein to a single DNA sequence. The affinity score of BoltzNet can be used to derive a measure of this intrinsic energy that is independent of ChIP coverage prediction. We thus sought to verify this using a biophysical assay of protein-DNA interactions, Biolayer Interferometry (BLI, Figure 5A).

**Figure 5:**
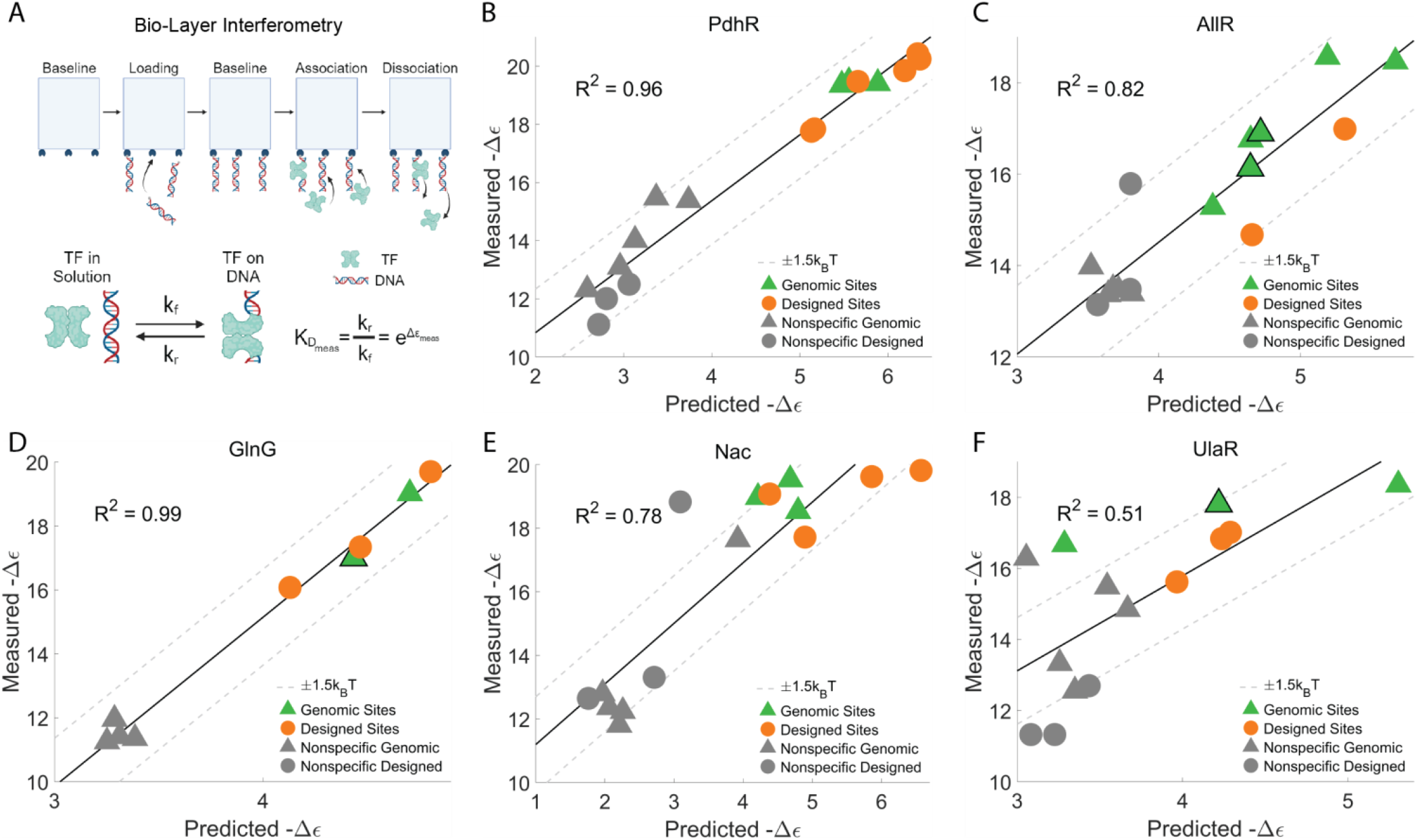
Validation of BoltzNet prediction of effective binding energy. Measurement of binding kinetics for purified proteins and 61 bp DNA sequences using Bio-Layer Interferometry (BL). **A)** BLI schematic. DNA sequences are attached to glass probes (loading-baseline). The probe is placed into a solution of TF and the kinetics of binding are measured (association). The probe is then placed into a solution without TF and the kinetics of unbinding are measured (disassociation). Based on a bimolecular model of binding, a measured effective binding energy is calculated. **B-F)** Scatter plots of measured effective energies (relative to solution) versus predicted effective binding energies for PdhR, AllR, GlnG, Nac, and UlaR. For each TF, sequences include sites from the genome (triangles) and novel sites generated using BoltzNet. All sites were tested against all TFs. Green and orange symbols are sites selected for the given TF. Grey symbols are sites selected for other TFs. Dotted lines represent bounds of +1.5kBt, equivalent to average thermal energy.

We performed BLI experiments on 5 TFs with both genomic sequences and novel sequences designed as in Figure 4A (and Figure S20 for Nac and UlaR). The kinetic measurements of BLI were used to estimate binding energies. Measured values were consistent with those for other DNA binding proteins^34^. Predicted values show a strong concordance with measured values (Figure 5), with R^2^ values between 0.51 and 0.99 spanning the strongest specific binding sites (green and orange) to non-specific sequences (gray) as well as genomic (triangle) and novel designed (circle) binding sites. Sequences that contain both single (no border) and multiple (black border) binding sites were accurately predicted, supporting the energy summation model used by BoltzNet. More generally, the results confirm the ability of BoltzNet to predict relative binding energies across a range of binding site strengths and configurations, and to extrapolate to sites stronger than any found in the genome.

### Large differences in enrichment reflect physiologically relevant differences in binding energy

Calibrating our model predictions with the results of Figure 5 allows us to relate ChIP-Seq coverage to quantitative binding energies. Figure 6 reveals that large changes in relative coverage between enriched regions result from differences in binding energy spanning ∼4 k_b_T. This is consistent with an estimated 5.8 k_b_T span of energies between the operators of LacI^18,34,35^. Weak binding sites are well within a physiological range of binding energy differences relative to strong sites. And our models predict that even small differences in sequence can differentiate weak from strong binding. Moreover, the accuracy of our models trained on weak binding regions suggests these sites contain information about the specificity of TF-DNA interactions. Many assays for DNA binding focus on cataloging the strongest binding sites. Our data support the view that weak sites are necessary for a complete picture of TF-DNA binding^10,36^.

**Figure 6:**
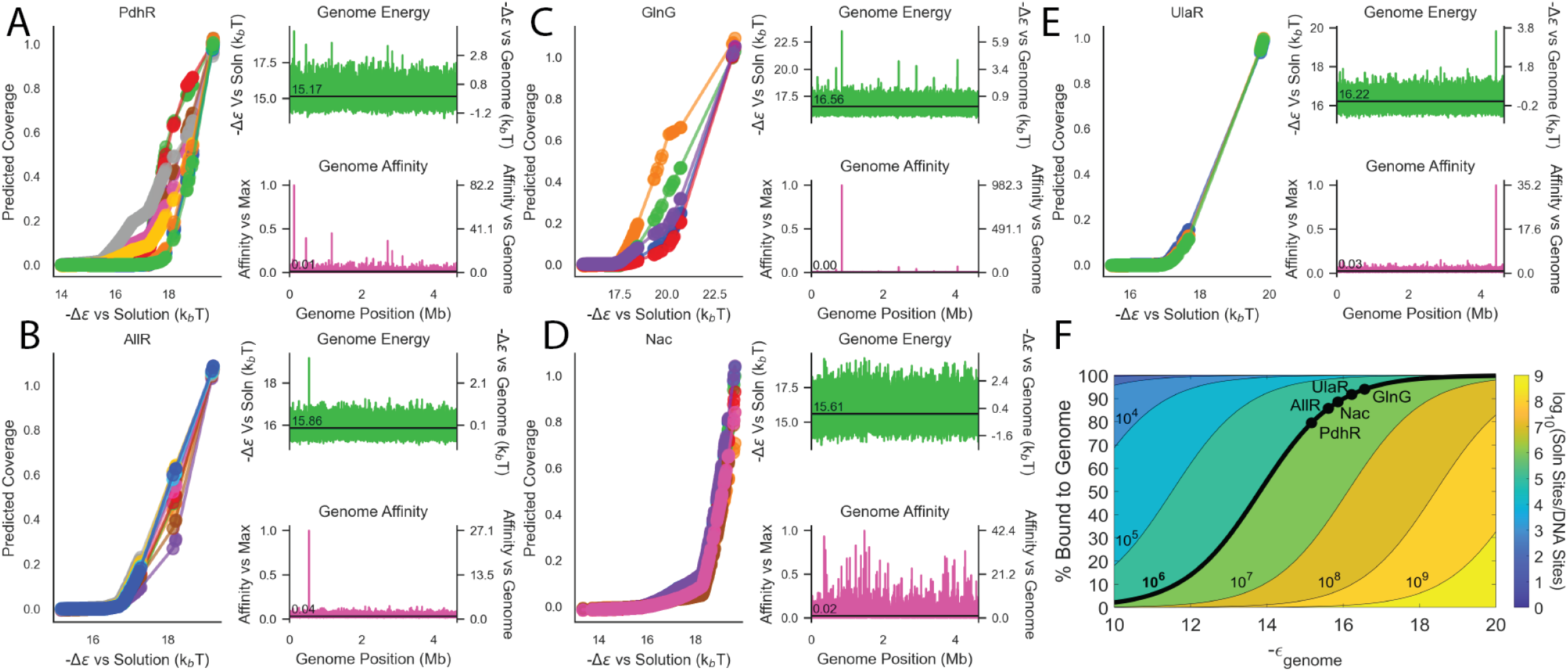
Implications of measured and predicted binding energies. **A-E)** Results for PdhR, AllR, GlnG, Nac, and UlaR. (Left panels) Predicted relative coverage vs predicted -Δε relative to solution for 101 bp sequences. Large relative differences in coverage arise from physiologically relevant differences in binding energy. (Top and bottom right panels) Binding energy versus solution (green left axis) and genome (green right axis) and affinity (exponentiated energy) relative to strongest region (pink left axis) and genome average (pink right axis). **F)** Probability of binding of a TF to genome versus solution as a function of energy of genome energy ε_genome_, (ε_soln_=0) and ratio of number of sites in solution to number of sites in genome (contours). Solid line shows solution sites/genome sites = 10^6^, points on the line display predicted probability that TFs in (A-E) are bound based on their average genome-wide energy. Results predict that most TF is bound non-specifically to genome relative to solution.

### Transcription factors primarily bound non-specifically to genome

The results of Figure 5 reveal that all 5 TFs have substantial binding energies to apparent non-specific sequences (grey symbols). Predictions of our calibrated models further suggest that all 101bp genomic sequences have substantial binding affinity (Figure 6A-E, green), with average genome values spanning ∼15-16.5 k_b_T. The cellular location of TFs has long been a question of interest^37,38^. An analysis based on our results predicts that the majority (>79%) of protein molecules for our studied TFs are bound “non-specifically” to genome rather than being free in solution. This is consistent with reports on specific TFs in a number of organisms^38-41^, though at odds with other reports^42^ possibly due to differences in background genome binding energy. It also has functional implications for TF binding and gene regulation^40,43-45^.

Our results also suggest the genome average as a more natural *in vivo* reference point for TF binding energies compared to solution. TF binding specificity can then be seen as the distribution of binding energies relative to this reference. Highly specific TFs (e.g. GlnG, UlaR, and AllR; Figure 6B,C,E) have lower variability in genome-wide binding energy and have one or a few sites with substantially higher affinity than this background. Less specific TFs (e.g. Nac, Figure 6D) have much higher genome-wide binding energy variability, with few sites substantially stronger but many sites moderately stronger. TFs like PdhR demonstrate an intermediate pattern. From this perspective, TF binding affinities *in vivo* exist on a spectrum with the genome. BoltzNet builds quantitative models of TF binding behavior that provide this genome-wide perspective.

## Discussion

We report the mapping of *in vivo* DNA binding for 139 *E. coli* TFs using ChIP-Seq. We also describe BoltzNet, a novel neural network that interprets ChIP-Seq data through the lens of a biophysical model of TF-DNA binding. Together, our results suggest a new paradigm for developing interpretable neural networks, and an alternative paradigm for describing TF-DNA binding. Most analyses of ChIP-Seq data catalog specific binding sequences, often by focusing on the most enriched binding regions. This effort is complicated by variability between experiments. Our data suggest that strong and weak binding exists on a spectrum of affinity to any sequence. Consequently, the notion of a discrete set of “canonical” TF binding sites is incomplete. A complete picture must predict binding affinity to any sequence, interpret the results of individual experiments, account for differences between experiments, and enable the quantitative design of novel binding sites.

We designed BoltzNet for these purposes. BoltzNet is trained on ChIP-Seq data and can predict coverage from any DNA sequence. However, the primary goal of BoltzNet is to learn an underlying thermodynamic model of TF-DNA binding affinity. BoltzNet is thus a bridge between high-throughput genomics and detailed biophysics. The model learns a single set of parameters that explain affinity across multiple genomic experiments. These parameters can then be used to design new sites with quantitative accuracy in measurements of purified protein and DNA. BoltzNet also learns a mapping from affinity to ChIP-Seq coverage that implicitly accounts for differences in experimental conditions, including TF concentration.

Crucially, we designed the architecture of BoltzNet for the physical process being modeled rather than the process of learning the model. This resulted in critical differences between BoltzNet and prior applications of CNNs to genomic data. One key difference was the use of an exponential activation function which is not typical in CNNs where it is common to re-scale and zero-center internal outputs to facilitate backpropagation. In the case of BoltzNet, the exponential activation was necessary to internally represent biophysically meaningful values, and empirically it was required to learn accurate models with only one weight matrix. By explicitly modeling a biophysical process, BoltzNet avoids issues of overparameterization and interpretability associated with black-box neural networks.

Our approach is more aligned with biophysical methods like ProBound but has significant differences. BoltzNet is designed for prokaryotic ChIP-Seq while ProBound is designed for eukaryotic SELEX. BoltzNet also only models TF-DNA binding in biophysical detail. Coverage, by contrast, is modeled as a functional mapping motivated by our prior work^22^. This significantly simplifies the BoltzNet model. BoltzNet then leverages the feature selection power of convolutions and the function approximation power of fully connected neural networks. By taking advantage of the flexibility and optimization engines of modern neural network frameworks, BoltzNet is also more easily trained, used, and adapted. BoltzNet thus acts as an important bridge between neural networks and thermodynamic models and provides a foundation for future biophysically inspired CNNs.

The application of BoltzNet to 125 *E. coli* TFs provides broad insight into prokaryotic TF-DNA interactions. Our results are consistent with previous reports that TF-DNA binding in prokaryotes is driven by localized interactions that can be modeled with a single energy matrix^15^. They also highlight the role of degenerate residues in sculpting overall affinity. Such residues may only be present in a few strong sites, and thus not apparent in PFMs. Our findings also reframe the discussion of weak and strong binding sites into real physical units. In doing so, they highlight the physiological relevance of weak sites and stress the importance of deep ChIP-Seq coverage to reveal weaker sites that are nonetheless important for understanding specificity. One important observation is that clusters of weaker sites can lead to occupancy equivalent to that of a single strong site, consistent with reports in other organisms^32^. Finally, our results speak to the long-standing question of the cellular localization of TFs^37,42^. For five TFs, we estimate >79% of protein is bound to the genome as compared to being free in solution. This conclusion depends on estimates of cell volume and only considers predictions in 101 bp sequences. It does not account for longer range interactions or sequence context. It is nonetheless consistent with previous reports and argues that “non-specific” genome binding is an important factor in gene regulation^37^.

Whether BoltzNet can be applied to other organisms is an important question for future study. Even within prokaryotes, more complex binding configurations including protein-protein interactions and DNA-looping are known to be important. An important advantage of BoltzNet is that it can be readily extended to encompass these additional interactions. Moreover, interactions can be systematically added to BoltzNet to test each hypothesis and determine the minimal model required for accuracy and generalization. Finally, while our results do not speak to the potential functional rules of the binding, the thermodynamic framework used can extend to gene regulation^16^, and we anticipate analogous extensions to BoltzNet to enable the study of gene expression.

## Methods

### ChIP-Seq Protocol

We developed a standardized ChIP-seq protocol based on previous methods (Figure S2)^7,46^. A more detailed protocol can be found in Supplementary Material (see ChIP-Seq). At a high level, we grew cells with different tagged TFs (see Strain Construction in Supplement) until log-phase in M9 minimal media supplemented with 0.4% glycerol as a carbon source. We crosslinked TFs to DNA and performed washes to prepare cells for sonication on a Covaris S2, which lysed cells and sheared DNA into fragments with a tight length distribution. We used ChIP-grade monoclonal antibodies in immunoprecipitation that target our engineered tags to specifically pull-down DNA bound to the TF of interest. This DNA was purified and prepared for NGS on Illumina’s NextSeq platform.

### ChIP-Seq Analysis Pipeline

We developed a standardized pipeline for identifying enriched regions from ChIP-Seq data based on previous methods^7,46,47^. This pipeline is described graphically in (Figure S4A). At a high level, we started by performing quality control on raw sequencing reads, before pruning Illumina adapter sequences from read ends, and aligning to the *E. coli* reference genome (Genbank Accession U00096.3). We calculated coverage from alignments as the number of reads aligning to each strand at each position, which was used to identify statistically enriched regions of coverage. We applied several filters to these regions, including: (1) filtering for the expected signature of forward and reverse coverage; (2) filtering for background enrichment in control experiments; and (3) filtering for ChIP-Seq specific artifacts that are not apparent in controls (Table S3). After filtering likely spurious regions, we performed ChIP-Seq-specific quality control to verify tagging of the correct TF. Results of all analyses were imported to a MySQL database.

After all samples of a given TF have been analyzed, we removed experiments with poor background coverage by filtering those with < 90% of bases covered with at least 1 read; as well as experiments with poor enrichment by removing experiments where the strongest region was < 10-fold enriched. For TFs with at least 2 experiments passing this filter, we merged the output regions from each experiment into a unified binding region set (Figure S4B). We found all overlapping regions and kept any that were found in at least two experiments, or one experiment and contains a RegulonDB known binding site.

### Mathematical Model of TF-DNA Binding and ChIP Coverage

We model a ChIP-seq experiment as a two-step process. The first step is TF binding to DNA, which we analyze through the lens of a previously described thermodynamic model of protein-DNA interaction^16-18,20^ resulting in the estimated probability that any given sequence will be bound by a TF at thermal equilibrium. The second step models ChIP coverage as a function of the probability of binding which we have previously shown can be treated as a signal convolution process^22,29^.

### Thermodynamic Model of Binding to a Single Site

To analyze TF-DNA binding, we seek to model the sequence-specific binding energy of a TF to DNA. In the simplest case, we consider a system consisting of one sequence of interest and N_NS_ non-specific sites. We then consider N_TF_ transcription factors, each of which can be either be bound to the sequence of interest with energy ε_seq_ or occupy a non-specific site with energy ε_NS_. By convention, more negative energies are more favorable. We further assume that occupancy of these states is in thermal equilibrium over the timescales of interest for our analysis, motivated by the fact that speed of binding reactions is very fast relative to the timescale of a typical ChIP experiment.

The probability that the sequence of interest will be bound by a TF can be calculated via the Boltzmann distribution^16^. This is accomplished by considering two states for the system: (1) an “unbound” state where all N_TF_ molecules occupy “non-specific” sites, and (2) a “bound” state where N_TF_ – 1 TF molecules occupy non-specific sites, and one TF molecule is bound to the sequence of interest. Each state is associated with a multiplicity, Z, which is the number of configurations by which that state can be realized. For example, for the unbound state, Z_unbound_ is the number of ways in which N_TF_ TFs can be arranged in N_NS_ non-specific sites. The energy of each configuration can be calculated by summing the energies of the N_TF_ TFs in that configuration. For example, for any unbound configuration:

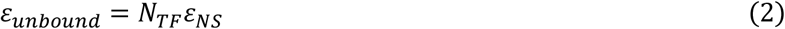

and for any bound configuration:

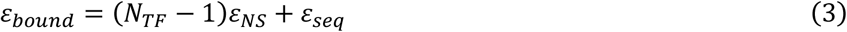

For each configuration, we then calculate the Boltzmann factor (also called statistical weight), defined as the exponential of the negative energy relative to k_b_T for that configuration where k_b_ is the Boltzmann constant and T is temperature (in K):

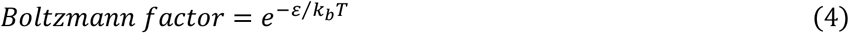

For the following, we assume that all energies are given in units of k_b_T so that this term will be omitted. The probability that the system will be in the bound state can then be calculated as the sum of the Boltzmann factors for all bound configurations divided by the sum of the Boltzmann factors for all configurations:

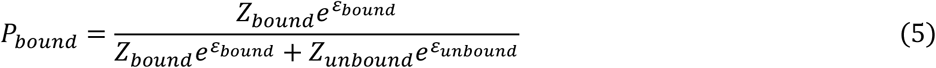

(where, as noted above, we assume energies are in units of k_b_T). The denominator of equation 4 is the partition function of the system. As shown previously^16,18^ (see also Thermodynamic Model of Binding to a Single Site in Supplement) for the simple system of one sequence and N_NS_ non-specific sites, equation 5 can be simplified to:

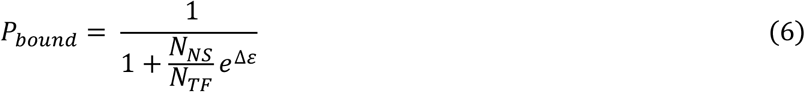

Where Δε is defined as the difference in energy between a single TF bound to the specific sequence and the TF occupying a non-specific site:

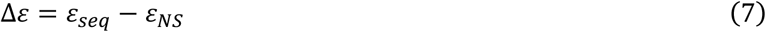

Δε can be interpreted as the energy released when a TF transitions from a non-specific site to the specific sequence, or conversely the energy that must be provided to dislodge a TF from the specific sequence to a non-specific site. From equation 6 more negative Δε increases P_bound_ and thus reflects a sequence with higher affinity to the TF relative to non-specific sites.

### Relationship to Hill Function

Equation 6 can also be related to a unimolecular model of TF binding to a DNA sequence:

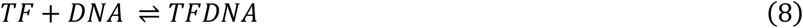

which can be used to derive the familiar Hill function form for TF binding at equilibrium (see Relationship to Hill Function in Supplement):

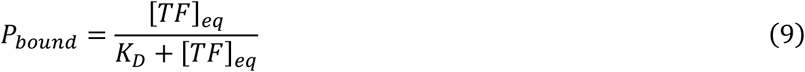

where [TF]_eq_ is the concentration of free TF at equilibrium, P_bound_ is the probability that the DNA is bound to TF, and K_D_ is the dissociation equilibrium constant, or the inverse of the association equilibrium constant, K_A_:

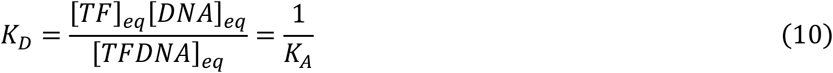

K_D_ (and K_A_) can in turn be related to the Gibbs free energy of the reaction (see Equilibrium Constant Relation to Free Energy in Supplement):

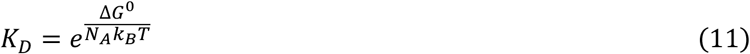

where ΔG^0^ is the Gibbs free energy associated with the reaction under standard conditions in which all reactants are present a 1M, and N_A_ is Avogadro’s number or the number of molecules in 1 mole.

Equations 6 and 9 can be related by letting 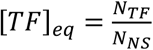 leading to:

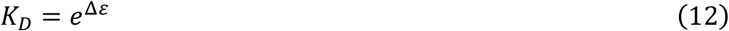

or equivalently:

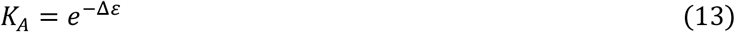

providing an interpretation of the equilibrium constants in terms of the change in energy (in units of k_b_T) when a single TF molecule binds DNA.

Moreover, relating equations 11 and 12, we can recognize that:

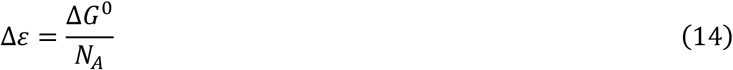

which indicates that the change in energy associated with TF binding per molecule (left) is the change in energy for 1 mole of molecules divided by the number of molecules per mole (right).

### Probability of TF binding to a sequence with multiple sites

A more realistic model assumes that each of N_region_ positions in a DNA sequence can be considered as an individual binding site i, each with their own energy ε_i_. As before, there are N_TF_ proteins available to bind any site within this region, and N_NS_ nonspecific sites that can be bound each with energy ε_NS_. We further make the simplifying assumption that only 1 TF can bind the sequence at any time. Determining the probability of binding anywhere in this sequence requires deriving the appropriate Boltzmann factors and multiplicities where the sequence is or is not bound, as in equation 4. As shown in Probability of TF Binding to a Sequence with Multiple Sites, the corresponding P_bound_ can be shown to be:

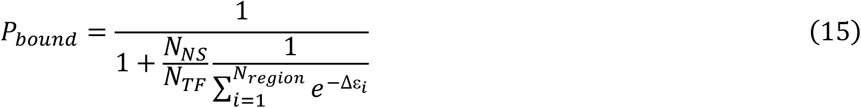

where Δε_i_ = ε_i_ - ε_NS_ as in equation 6.

If we then consider the entire sequence as a single potential binding site with an effective binding energy of Δε_eff_, we can relate equations 6 and 15 to derive:

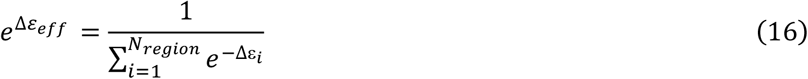

which implies that the effective energy of the sequence is related to the log of the *e*^−Δε^_*i*_ of each site in the sequence:

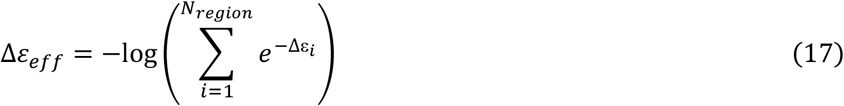

### Additive model of binding energy

In general, the energy of binding of proteins to DNA could depend on the underlying sequence through numerous complex non-linear interactions. However, several studies have demonstrated that simple additive “energy matrix models” models of are often sufficient to describe TF-DNA binding energies^17-19,27,28^. In such models, each base in a sequence contributes an additive term to the overall binding energy. Such models can be seen as first-order Taylor Series approximations to more complex models. Mathematically, we implement such models using one-hot strategy to represent a sequence of length L as an 4xL matrix S_ij_, where i represents the four bases (A,C,G,T) and S_ij_=1 if an only if the base at position j is i. The binding energy matrix M_ij_ in turn contains 4xL values where M_ij_ is the energy contributed by base i at position j. The total energy of the sequence is then the inner product of both matrices:

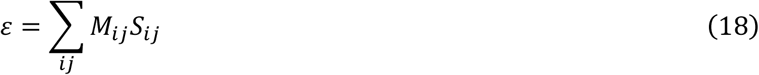

### Signal processing model of ChIP-Seq Coverage

We have previously shown that ChIP-Seq coverage can be approached from a signal processing perspective as a signal convolution^22,29^. In this context, an impulse signal represents a binding site and the process of ChIP-seq emits a corresponding impulse response. The magnitude of the impulse signal is assumed to be proportional to the probability that the site is bound. The impulse response spreads out the signal of the binding site due to the shearing and sequencing process of ChIP-Seq. An analysis of this process revealed that the impulse response can be modeled as an extreme value distribution, resulting in an impulse response that follows a Gumbel distribution ^22^. The sum of impulse responses from all TF binding sites in a region generates an observed ChIP-seq enrichment peak.

### BoltzNet Architecture and Implementation

BoltzNet is a neural network designed to predict ChIP coverage from sequence by directly estimating a model of TF-DNA binding energy. Following the mathematical model described in the previous section, BoltzNet consists of two components: one that models the processes of TF-DNA binding and one that models coverage in ChIP experiments (Figure 1A). BoltzNet is implemented using TensorFlow v2.7.0.

### TF-Binding component

The TF-DNA binding component takes DNA sequence as input and models the effective equilibrium association constant of TF binding. This component is a specifically designed convolution model. DNA sequences are input using one-hot encoding, as described above, to create a 4xL matrix where L is the length of the sequence. To model TF-DNA binding in either forward or reverse orientation, one-hot versions of the forward and reverse complemented strands of the input sequence are then concatenated together to generate a 4x(2L) matrix. This matrix is then input to a convolution layer.

The BoltzNet convolution layer follows a specific biophysically motivated design: it consists of a single convolution weight matrix (kernel), with a bias term, that feeds into an exponential activation function. We use a single convolution weight matrix of length 25 to model the energy of binding to each 25-mer in an input sequence relative to a constant reference energy:

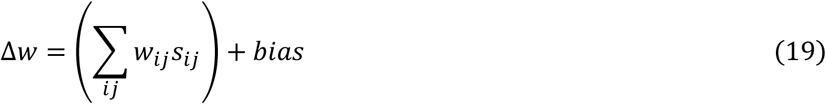

where s_ij_ is the one-hot encoded 25-mer. Because positive values of weights contribute positively to neural network outputs, the weight matrix models the negative of the relative binding energy.

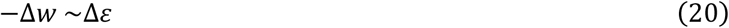

An exponential activation function is then used to generate the exponentiated energies required by the Boltzmann distribution, so that the output of the convolution layer models the exponentiated relative energy at each position, p, of a sequence.

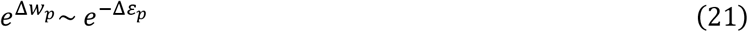

Exponential activations are not typical in CNNs where the values of internal layers are not intrinsically meaningful, and the common practice is to minimize and zero-center internal outputs to facilitate backpropagation. In the case of BoltzNet, it is key to ensuring that internal layers represent verifiable physical values.

The output of the convolution is then passed through two pooling layers. The first pooling layer, SumStrands, sums the contribution from each strand at each position (implemented using a tailored 2D Convolution layer). The second sums the value at each position of the SumStrands layer and divides this value by a constant, C_ave_, that depends on the length of the SumStrands layer (implemented using a 2D Global Average Pooling layer). We refer to the output of the final pooling layer as the Affinity Score of the sequence, and mirroring equation 16, we interpret this score as the exponent of an effective weight, Δw_eff_, for the sequence:

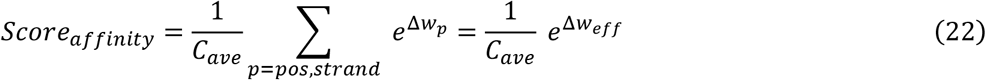

Then, we relate equation 22 with equation 17 using equation 20 to derive:

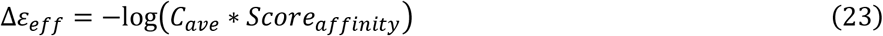

### ChIP coverage component

The second component takes the output of the final pooling layer as input and models the coverage in one or more ChIP experiments as a function of this value. This component follows the signal processing model of ChIP-Seq described above. In this context, the connection between Δε_*eff*_ for a sequence and expected coverage involves two transformations. First, from equation 6, Δε_*eff*_ implies a probability of binding. Second, binding to the sequence gives rise to impulse response of coverage that is proportional to P_bound_, and coverages from nearby sites combine additively (see above). This two-step process implies a function from 1/K_A_ to coverage that is monotonically increasing and smooth, and that differences in coverage between different experiments for the same site are expected to be due in large part to differences in TF protein concentration which alter P_bound_ (via equation 15), and differences in IP efficiency which alter the proportionality between coverage and P_bound_.

We learn this function for each experiment using a fully connected neural network. This approach takes advantage of the well-known universal function approximator nature of fully connected neural networks with at least one hidden layer with non-linear activations^30^. BoltzNet employs a neural network with one input layer with one node, one hidden layer with 140 nodes, and one output layer with one node per experiment. The input and hidden layer nodes use Leaky ReLU activation functions, while the output layer uses a linear activation function appropriate for regression. Both deep (multiple hidden layers) and wide (few hidden layers with many neurons) neural networks have the capability for universal function approximation, with deep neural networks learning certain classes of functions more efficiently^48-51^ at the expense of potential vanishing gradients^52^. In the case of BoltzNet, the expected function does not fall into the class of functions that are expected to require a deep architecture. And empirically, the use of a deep architecture did not outperform the architecture with one hidden layer.

### BoltzNet Training

BoltzNet is trained using a set of training sequences and associated coverage values. Training sequences consist of all the called enriched regions (see ChIP-Seq Analysis Pipeline) for a TF (positive samples) as well as 5,000 randomly selected genomic regions (negative samples). Sequences of 101 bp are used consisting of 50 bp on either side of the center position of a sequence. For positive samples, the center position is the location of maximum coverage for a called region and the coverage value is the maximum coverage across the region of the sequence in each experiment after masking known artifacts (see Filtering of Artifacts), normalized to 1 where 1 is the strongest region. For negative samples, the center positions are chosen randomly from all genomic positions that are at least 500 bp from any positive sample and the coverage value is set to 0 in all experiments.

To increase model generalization, we augment positive training samples using circular shift permutation^53^. Each positive sequence is augmented with two additional sequences – one sequence shifted 1 bp in the 5’ direction and one shifted 1 bp in the 3’ direction. Each positive sequence is thus represented three times in the training set – once with the original sequence and twice with permuted sequences. The coverage values for all three sequences are the same as the original sequence.

The training of neural network parameters through gradient descent via backpropagation is sensitive to the starting values of the parameters. Initial values may constrain models to regions of the parameter space that are less optimal. To address this, we train BoltzNet in two stages. First, we pre-train 10-15 models (implemented with multiple-process support) for a restricted number of epochs on all the positive data and number of negative samples equal to the number of called regions times 3. Second, the pre-trained model with the highest accuracy then underwent additional training on the entire positive and negative training set. Empirically, we observed that this process increased the probability that the binding sites in each sequence associated with higher affinity binding energies would be discovered. However, the limited number of negative samples used during pre-training typically meant that pre-trained models would have a greater number of false positives on the full genome sequence. Full training on the best pre-trained model with all negative sequences typically increased model specificity on the full genome by further learning parameters associated with non-specific binding energies.

All training was performed using the Adam optimizer^54^, with an exponential decay learning rate schedule, a mean-squared error loss function, a batch size of 256, and early stopping with patience of 100 epochs. Pre-training was performed over a maximum of 1,000 epochs. Full training was performed over a maximum of 4,500 epochs.

Regularization was essential for accurate model learning and prediction. L1 kernel, activity, and bias regularization were applied to both the convolutional layer and all neural network layers. Conversely, since our goal was to train deterministic models whose predictions and parameters could be experimentally verified, we did not incorporate Batch Normalization layers which would re-scale and re-center parameters^55^. Nor did we incorporate Dropout layers^56^ which can lead to non-deterministic results during training. Empirically, neither was required for accurate training of our models.

We applied BoltzNet to *in vivo* ChIP-Seq data for all 139 TFs and generated 125 models that passed two accuracy criteria: R^2^ > 0.6 on predicting coverage on the training set and specificity of more than 60% when applied to the entire genome. Of the models for the 14 TFs that do not pass, 12 failed to model the training set with R^2^>0.6 (allS, csgD, csiR, gadX, hcaR, hipB, maze, nanR, stpA, yafC, yfiE, yjgJ) while the remaining two (aaeR, rutR) failed to achieve >60% specificity when applied to the entire genome (specificity of 48% and 40% respectively).

### Calculating PFMs from Weight Matrices

To calculate Position Frequency Matrices from convolution weight matrices, we used a previously described method^31^. The method counts nucleotide frequencies in sequence sites that score above a threshold when multiplied by the weight matrix. For a given weight matrix, we scan all positive training sequences and identify all locations that score above some fraction of the maximum score for that matrix (typically 0.8 maximum). The frequencies of bases in each position of all such locations is used to construct the corresponding PFM. All sequence logos were generated using Logomaker^57^.

### Calculation of Base Contributions

As described above, the weight matrix provides a direct prediction of the relative contribution of every base at any binding location and orientation, the exponentiation of which provides a measure of the affinity at each location and orientation. The affinity of the whole sequence is determined by the affinity of all locations and orientations. Any given base thus contributes to the overall affinity of the sequence through all overlapping binding locations in both orientations. To calculate a measure of this contribution, for every base we calculate the weight of the base in each overlapping binding location multiplied by the model affinity (exponentiated kernel score) of that location and sum these values over overlapping locations. These values are then used to scale the base in plots of base contributions.

### Alignment of PFMs

To align PFMs we transformed frequency matrices into information matrices and compared every possible motif alignment (including reverse-complementing one motif, see Motif Comparison). We obtained PFMs from RegulonDB v12.0^23^ based on sites for 36 TFs with strong classical binding evidence for comparison to PFMs from BoltzNet, allowing us to demonstrate that weight matrices can recover existing PFMs, while providing a more quantitative explanation of TF sequence preference.

### Prediction of Novel Binding Sites

For each TF, we selected a 101 bp reference sequence. This sequence was chosen as the strongest enriched region with a single binding site. We then manually selected 10 bases to vary in each sequence, focusing on accessory bases outside the bases that were the most conserved. We generated all combinations of all of the 10 bases while keeping the remaining bases the same. This resulted in a range of predicted affinities. We selected sequences from this range, including where possible predicted sites stronger than the strongest singleton genomic site. These selected sequences were then trimmed to 61 bases for ordering oligos. The trimmed sequences were rerun through BoltzNet (with Ns at trimmed positions) to generate final predictions for all selected sequences. Oligos with the selected sequences were then ordered from Integrated DNA Technologies (IDT), and used for validation with either Library-ChIP and/or BioLayer Interferometry.

### Library-ChIP

Library-ChIP is performed according to previously published methods^7,46,47^. We used the same tagged-TF strains that were used in ChIP-seq, carrying an additional single-copy plasmid containing one of ∼100 different designed binding sites (see Library-ChIP Library Construction). Cultures were treated identically to those in ChIP-Seq experiments up through sonication. After pelleting the sonicated lysate, we saved ∼10% of the DNA-containing supernatant as a pre-IP control. The remaining 90% of supernatant went through immunoprecipitation as a normal ChIP-Seq experiment would. Both the pre- and post-IP samples underwent protein digestion, and DNA was purified. DNA was converted into NGS libraries using a custom protocol, then pooled for sequencing in the same manner as our ChIP-Seq samples (see Library-ChIP Library Prep and Pooling). After retrieving the sequencing reads, we counted the abundance of each designed sequence and normalized by sequencing depth, then took the ratio of post-IP/pre-IP counts as the Library-ChIP enrichment (see Analysis).

### Protein Purification

To purify proteins for *in vitro* experiments, we grew 500mL cultures of each TF-6xHis construct (see Inducible Tagging for *in vitro* Experiments). After cultures reached log-phase, we induced TF expression by adding rhamnose to a final volume of 0.2% and allowed induction to continue overnight. We purified TFs with Qiagen’s Ni-NTA Spin Kit (#31314) and dialyzed against a storage buffer (50mM Tris-HCl, 0.1mM EDTA, 1mM DTT, 200mM KCl, 10mM MgCl2, 50% glycerol^58^; GlnG was stored in 10mM Tris-HCl pH 8, 50mM NaCl, 1mM DTT, 1mM EDTA, 50% glycerol^59^) using desalting columns (Cytiva cat. #17085101). TF expression and purity were assessed with SDS-PAGE and concentrations were determined with the Bradford Assay using BSA as a standard. 10μM stocks were made in respective storage buffers and proteins were stored at -80°C for downstream experiments.

### BioLayer Interferometry

BLI was carried out using a previously published protocol that we optimized for our application^60,61^. We loaded biotinylated DNA on an Octet Red 384 using Streptavidin Tips (Sartorius cat. #18-5020). All BLI was performed at 30°C with shaking at 1000rpm in binding buffer (20mM Tris-HCl, 0.1mM EDTA, 10mM MgCl2, 1mM DTT, 120mM KCl, 5% glycerol, and added 0.05% Tween-20 as a blocking agent^62^). A 1:1 ratio of 10nM biotin and biotinylated-dsDNA in binding buffer was used for loading and different TF concentrations were used as analyte. For specific DNAs, 50-100nM proved to be a sufficient upper-limit of TF concentration and a 2-fold serial dilution was used from that upper limit for a total of 6 TF concentrations and a 0nM TF reference. For non-specific DNAs, more protein was needed for an observable signal and we used TF at 100, 250, 500, 750nM, and 1uM in addition to the 0nM TF reference. We performed an initial 10-minute pre-incubation where the tips are soaked in buffer A, followed by a 60 second baseline reading. We loaded DNAs for 2 minutes and took another baseline reading for 60 seconds to align traces before TF binding. TF was allowed to bind to probe-bound DNAs with a 10-minute association step before probes were returned to a well of buffer to monitor unbinding in a 15-minute dissociation phase.

This assay followed the design for fitting with the 1:1 binding model, which corresponds to a unimolecular TF-DNA binding reaction. We implemented this model in MATLAB to estimate dissociation constants for each TF-DNA pair and calculated binding energies from equation 12 (see Analyzing BLI).

## Acknowledgements

We acknowledge the Boston University’s Microarray and Sequencing Resource Core Facility where the majority of NGS was performed. We particularly would like to acknowledge Yuriy Alekseyev and Ashley Leclerc for their sequencing expertise. We acknowledge the Center for Macromolecular Interactions at Harvard’s Medical School where we performed BLI experiments and received assistance from Kelly Arnett in troubleshooting and optimizing our protocol. We acknowledge the RegulonDB team, particularly Socorro Gama-Castro who contributed to curation addressing issues from our experiments, and Heladia Salgado for her contributions to data management and representation in RegulonDB. We acknowledge the Wadsworth Center Applied Genomic Technologies Core Facility where some NGS was performed. We acknowledge the Wadsworth Center Media and Glassware Facility for providing supplies to perform strain construction and some experiments. We acknowledge Anne Stringer who provided technical support in several areas related to experimental methodologies. We acknowledge BioRender which assisted in the generation of figures 5A, S1, S3, S4, S9, S10, S21.

## Authors Contributions

Conceptualization – JEG, JCV, JTW

Data Curation – PL, LGR, VHT, PA, CR, JEG

Formal Analysis – PL, LGR, JEG

Funding Acquisition – JCV, JTW, JEG

Investigation – PL, LGR, VHT, PA, XZ, JEG

Methodology – PL, PA, JTW, JEG

Project Administration – JCV, JTW, JEG

Resources – PL, PA, SK, GB, JP, CS, MB, JTW, JEG

Software – PL, LGR, JEG

Supervision – JCV, JTW, JEG

Validation – PL, LGR, JEG

Visualization – PL, LGR, JEG

Writing Draft – PL, LGR, JEG

Reviewing & Editing – PL, LGR, VHT, JCV, JTW, JEG

## Data and Materials Availability

Raw data will be made available in NCBI’s Gene Expression Omnibus (GEO).

Processed data will be uploaded to RegulonDB.

Models will be made available through RegulonDB.

