## Supplementary Material for "Predictive Biophysical Neural Network Modeling of a Compendium of *in vivo* Transcription Factor DNA Binding Profiles for *Escherichia coli*"

### Supplementary Methods

#### *Strain Construction*

We used a total of 4 tagging methods for TFs presented here, 3 of these were used for *in vivo* ChIP-Seq and Library-PCR, these included two methods for tagging TFs at their genomic locus for native expression levels, and one method for tagging TFs on an inducible plasmid to help achieve adequate TF expression. Additionally, we used one tagging method to create strains that allowed protein purification for *in vitro* ChIP-Seq and BLI.

**Native Tagging for *in vivo* ChIP-Seq.**
TFs tagged at their native locus were generated with either P1 transduction [1] or the FRUIT method [2]. Across these methods, we used 3 different tags (Figure S1). We tagged 133 TFs with a SPA tag and 64 TFs with a 3xFLAG tag using P1 transduction by initially introduced into *E. coli* strain DY330 using recombineering as described previously [3] and transferred to MG1655 by P1 transduction before removing the KanR marker with FLP recombinase [4]. Additionally, 134 TFs with a V5-tag were made with FRUIT. This yielded 314 TFs with at least one of these tags.

**Inducible Tagging for *in vivo* ChIP-Seq.**
We generated 311 pBAD24-3xFLAG (Figure S2) inducible strains for *in vivo* ChIP-Seq by performing PCR on genomic DNA using primers that targeted the TF of interest (Table S4). The forward primer included an overhang with homology 5' to the pBAD insert site as well as a start codon followed by the first 15-18 bases of the TF sequence after the start codon. The reverse primer included an overhang with homology 3' of the pBAD insert site followed by the last 15-18 bases of the TF, omitting the stop codon. Homologous overhangs matched a KpnI insertion site on pBAD24. Plasmid was linearized with KpnI from NEB (#R0142) and purified with Qiagen's PCR Purification Kit (Qiagen #28104). PCR fragments and linearized vector were ligated with NEB HiFi Assembly Mix (NEB #E2621) and ligation was transformed by heat shock back into *E. coli K-12 MG1655*. Colonies were restreaked and sent for Sanger sequencing for clone validation. This generated 311 TFs tagged with this method.

**Inducible Tagging for *in vitro* Experiments.**
We generated tagged strains for AllR, GlnG, Nac, PdhR, and UlaR using Lucigen's Expresso Rhamnose Cloning System (#49012-2) to create inducible 6xHis-tagged TF strains that can produce TFs easily purified for *in vitro* assays. Briefly, we followed kit instructions to clone each TF into the pRham vector included with the kit, where the 6xHis tag was added on the C-terminal end. Clones were selected using kanamycin and verified with Sanger sequencing, valid strains were preserved in 50% glycerol stocks.

#### *ChIP-Seq*

We have established a ChIP-Seq method that we have now applied to a range of bacteria (Figure S3) [5-9]. The method involves the following steps: (1) growing cells and inducing TFs if necessary; (2) crosslinking, DNA shearing, and DNA extraction; (3) immunoprecipitation; (4) purification of immunoprecipitated DNA; and (5) creation of NGS libraries for sequencing from purified DNA. These steps are described in more detail below.

**Standardized Culture Conditions and (Optional) TF Induction.**
For *E. coli* experiments reported in this paper, we used a standard culture condition for all experiments to prevent potential media-dependent differences in binding. We used M9 minimal media supplemented with 0.4% glycerol as a carbon source and 100μg/mL ampicillin for inducible strains. Cultures were inoculated from glycerol stocks into 5mL media and grown at 30°C with shaking at 250rpm for a day then subcultured into 50mL media in vented flasks until OD ~0.5. For inducible ChIP-Seq, we added arabinose to a final concentration of 0.2% and continued growth for 1hr to allow TF induction before crosslinking.

**Crosslinking, DNA Shearing, and Extraction.**
Cells were crosslinked to fix all TFs bound to DNA by adding formaldehyde to a final concentration of 1% and shaking at 100rpm, RT for 30min. Crosslinking was ceased by adding glycine to a final concentration of 250mM and shaking for 15min at 100rpm, RT. Crosslinked cultures were prepared for searing by pelleting in a 4°C centrifuge at 4000rpm for 10min and washed twice in 50mL PBS. A mixture of Buffer 1 (20mM HEPES Potassium Salt pH 7.9, 50mM KCl, 0.5mM DTT, 10% glycerol) + Protease Inhibitor (PI) (Sigma Aldrich #04693116001) was used to prepare cells for shearing via sonication while protecting TFs from degradation. This was made by dissolving 1 PI tablet in 1.5mL DI water and adding 8.5mL Buffer 1. After the second wash, cells were resuspended in 600μL Buffer 1+PI mixture, transferred to Covaris-compatible tubes (Covaris Cat. #520056), and left on ice until sonication.

We used a Covaris S2 with an optimized sonication protocol to shear DNA. This allowed us to obtain DNA fragments with a tight length distribution centered around 250bp, which proved critical in our analysis of the data. Cells were sheared for 12min with amplitude 20%, intensity=5, and 200 cycles/burst. After sonication, lysate was transferred to 1.5mL centrifuge tubes and spun at 13000g, 4°C for 10 min. DNA was extracted by transferring supernatant to fresh 1.5mL tubes. Salt adjustment was performed by adding Tris-HCl pH8, NaCl, and NP-40 to final concentrations of 10mM, 150mM, and 0.1%, respectively.

**Immunoprecipitation.**
We used ChIP-grade antibodies for each epitope tag, ensuring specific pulldown of our TF of interest and its bound DNAs, while minimizing nonspecific pulldown. For TFs tagged with SPA or 3xFLAG, we used Anti-FLAG antibody (Sigma-Aldrich #F1804), and for V5-tagged TFs, we used Anti-V5 antibody (Sigma-Aldrich #V8012). 5μL of antibody was added to the salt-adjusted supernatant and samples were rocked at 4°C overnight for primary antibody binding.

We used Protein G Agarose Beads (Thermo Scientific #20399) to pull down antibody-bound TFs. These interact specifically with the antibodies added above and allowed us to wash away the rest of the lysate, keeping only the TF of interest and its bound DNAs. We washed 50µL of beads by adding 1mL IPP150 (10mM Tris-HCl pH 8, 150mM NaCl, 0.1% NP-40) buffer. We pelleted the beads by spinning at 2000g at RT for 2min and discarding buffer. Lysate with primary antibody was added to the bead pellet and rocked at 4°C for 30min, followed by another 90min of rocking at RT for primary & secondary antibody binding. We pelleted the beads at 2000g for 2 min, discarded the lysate, and started washes. These were carried out by adding 1mL of buffer, rocking tubes for 2min, spinning for 2min at 2000g, and discarding the supernatant; 5 washes were done with IPP150 as buffer, followed by 2 washes with 1x TE.

**DNA Purification.**

To purify immunoprecipitated DNA, we eluted TF-DNA complexes from beads by adding 150µL EBB (50mM Tris-HCl pH 8, 10mM EDTA, 1% SDS) to the bead pellet and incubated at 65°C for 15 minutes. Tubes were spun at 2000g for 5 min, and the elution was transferred to a fresh 1.5mL tube. We performed a second elution using 100µL TE + 1% SDS, incubated at 65°C for 5 min, and spun at 2000g for 5 min. The two elutions were pooled for a total volume of 250µL. We added 12.5µL Proteinase K (Sigma-Aldrich #P4850) to the pooled elution and incubated at 37°C for 1hr to digest proteins, releasing DNA from TF-complexes, followed by incubation overnight at 65°C to deactivate the proteinase. ChIP DNA was purified using Qiagen's PCR Purification Kit (Qiagen #28104) using standard procedures, with a final DNA elution volume of 35µL EB.

**Library Preparation.**

We converted purified ChIP DNA to NGS libraries for sequencing with New England Biolabs' NEBNext Ultra II Library Prep Kit for Illumina (E7645). Library prep consists of the following steps, where we modify reaction volumes as described below: (1) end preparation; (2) adapter ligation; (3) adapter cleanup; (4) library PCR; (5) size-selection; and (6) library quality control and pooling for NGS. These steps are described in more detail below.

<u>End Preparation.</u> The first stage blunted fragment ends, phosphorylated the 5' nucleotides, and added a dA tail on the 3' nucleotides to prepare the ends of DNA for adapter ligation. This was achieved by mixing 33.3µL ChIP DNA with 2µL End Prep Enzyme Mix and 4.7µL End Prep Reaction Buffer and incubating on a thermocycler at 20°C for 30min, and 65°C for 30min.

<u>Adapter Ligation.</u> We ligated adapters with sequences for targeting in PCR by diluting NEBNext Adaptor for Illumina 1:25 in 10mM Tris-HCl pH 8 + 10mM NaCl (as suggested for low-input samples) and adding 1.67µL of diluted adapter to the end prep reaction along with 20.67µL of premixed ligation master mix and ligation enhancer. This reaction was incubated at 20°C for 15min with the thermocycler lid off. Then we added 2µL USER enzyme and incubated for another 15min at 37°C with the thermocycler lid on and set > 47°C to complete adapter ligation.

<u>Cleanup of Adapter Ligation.</u> Samples were cleaned up to remove residual adapters that could prevent efficient library PCR using AMPure XP beads (Fisher Scientific #NC9933872). Briefly, we added 60µL room temperature beads to the adapter ligation reaction (~0.9X bead ratio) and incubated for 5min allowing adapter-ligated DNA to bind the beads, while shorter fragments remained free. Samples are moved to a magnetic rack to separate the DNA-bound beads. The supernatant was discarded, and beads were washed twice with 200µL 80% ethanol. After the second wash, tubes were removed from the magnetic rack, residual ethanol was allowed to dry with the tube tops open (without over-drying beads which can reduce

DNA recovery), and beads were resuspended in 11µL 0.1X TE. We allowed this to incubate for 2min which separated adapter-ligated DNA from the beads before returning the tubes to the magnetic rack. After beads were separated from the eluted DNA, we transferred 9µL of the DNA-containing supernatant to PCR fresh tubes.

Library PCR. We performed PCR on the cleaned-up adapter-ligated fragments to amplify DNA while adding Illumina NGS adapters and unique barcodes that were used to identify each sample on an NGS flowcell. This was done by adding 15µL NEBNext Ultra II Q5 Master Mix to the 9µL DNA fragments. For samples sequenced with NextSeq500, we used NEBNext Multiplex Oligos for Illumina (E7335, E7500, E7710, and E7730) for multiplexing. We added 3µL of one of the 48 i7 primers to each sample, and 3µL of the universal i5 primer to all samples. For samples sequenced with NextSeq2000, we used unique dual indexes (E6440, E6442, E6444, E6446, E6448). These primer sets have i5 and i7 primers premixed, so we added 6µL of a primer pair to each sample. Reactions were mixed and thermocycled with an initial denaturation step of 30sec at 98°C, followed by 14 cycles of 10sec denaturation at 98°C then 75sec annealing/extension at 65°C. After the last cycle, a final elongation was performed at 65°C for 5min, followed by cooling to 4°C until samples were retrieved.

Size-Selection. After library-PCR, we performed size-selection to retrieve adapter-linked DNA fragments with a tight size distribution centered around 350bp. This was done by increasing the PCR volume to 100µL with water and adding 65µL room temp AMPure XP beads for a 0.65X bead ratio. We separated beads and unbound DNAs by incubation on a magnetic rack and transferred supernatant (which contains the library fragments of interest) to fresh PCR tubes. Left-side selection was performed by adding 110µL of beads to the supernatant for an approximate bead ratio of 1.8X. Beads were washed according to the same procedure for cleanup after adapter-ligation, namely magnetically separating the beads and washing in 80% ethanol twice. Library DNA was eluted in 33µL 0.1X TE.

Library QC, Pooling, and Sequencing. Libraries were analyzed with on-chip electrophoresis to verify size distributions fit the expected profile and to obtain library molarities for pooling before NGS. We used Agilent's Bioanalyzer 2100 with High-Sensitivity DNA chips (Agilent #5067-4626). We typically diluted libraries 1:10 in water to ensure concentrations were within Bioanalyzer's measurable range. To create pools for NGS, each library was normalized to 4nM concentration, and equal volumes of each library were added to the pool. In cases where library concentration was < 4nM, we added an equimolar volume of library to the pool. Libraries were sequenced on either a NextSeq500 or NextSeq2000 with 75bp single-end reads. In a few cases, 75bp paired-end sequencing was performed.

#### *In Vitro ChIP-Seq*

For five TFs, we performed an *in vitro* variant of ChIP-Seq. Our *in vitro* ChIP-Seq protocol differs from *in vivo* ChIP-Seq by utilizing pre-sheared gDNA and using purified epitope-tagged TFs to bind gDNA *in vitro*. The overall workflow is shown graphically in Figure S4, and gDNA shearing and binding reactions are described in more detail below, along with a brief discussion of the relationship between our *in vitro* ChIP-Seq protocol and DAP-Seq.

##### Generation of Sheared gDNA.

We generated sheared gDNA for *in vitro* ChIP-Seq by extracting gDNA from WT *E. coli* and shearing to the 250 bp size targeted for *in vivo* ChIP-Seq. We started by growing WT *E. coli* in 2mL LB from glycerol stocks at 37°C with shaking at 250rpm in vented tubes. Genomic DNA was extracted using the GenElute Bacterial

Genomic DNA Kit from Millipore-Sigma (#NA2110). gDNA was sheared by increasing gDNA volume to 500µL with 1x TE buffer, and sonicating on a Covaris S2 for 12min with amplitude 20%, intensity=5, and 200 cycles/burst. Fragments with the correct size were retrieved by mixing the sample from Covaris with 750µL of AMPure XP beads in a 1.5mL tube (1.5x bead ratio) and incubating for 10 minutes. Tubes were transferred to a magnetic rack for 5 minutes, isolating beads bound to target gDNA fragments. Beads were washed twice with 200µL 80% ethanol, and air dried for up to 5 minutes, ensuring not to over-dry the beads. gDNA fragments were eluted by resuspending beads in 150µL 0.1x TE buffer, incubating for 5 minutes off the magnetic rack, followed by 5-minute incubation on the rack. After beads separated, supernatant containing sheared gDNA was transferred to a new tube and ran on Bioanalyzer to confirm gDNA concentration and fragment size.

**_In Vitro_ Binding of Purified Proteins.**
We combined gDNA and different TF concentrations to assess the relationship between TF concentration and binding. We created a Protease Inhibitor-buffered IPP150 and by dissolving 1 tablet of Protease Inhibitor (Sigma Aldrich #04693116001) in 10mL IPP150 (10mM Tris-HCl pH 8, 150mM NaCl, 0.1% NP-40). We combined 500ng gDNA and either 50ng or 500ng purified 6xHis-tagged TF in the PI-buffered IPP150, with a final reaction volume of 500µL. This was mixed and allowed to incubate for 1hr at RT for protein binding. After an hour, we added 5µL anti-6xHis antibody (Invitrogen cat #MA1-21315) and allowed tubes to rock at 4°C overnight. All subsequent steps from immunoprecipitation to analysis were carried out identically to *in vivo* ChIP-Seq.

**Comparison to DAP-Seq.**
During the completion of this work, a similar *in vitro* binding assay called DAP-Seq was reported [10, 11]. Our in vitro ChIP-Seq protocol differs from DAP-Seq in two primary ways. First, we purify protein as described above while DAP-Seq uses an in vitro expression and purification protocol. Moreover, a variant of DAP-Seq called biotin-DAP-Seq introduces biotinylated lysines during translation for use in IP. Second, DAP-seq begins by performing library preparation on pre-fragmented gDNA prior to IP, whereas our *in vitro* ChIP-Seq protocol performs library preparation after IP.

#### *ChIP-Seq Analysis Pipeline*

We developed a comprehensive pipeline for analyzing our ChIP-Seq data (Figure S5A). Applied to *E. coli*, the high-level steps the pipeline performs are as follows: (1) quality control on raw reads; (2) trimming adapters and low-quality bases from reads; (3) aligning the reads to the *E. coli* reference genome; (4) generating read coverage matrices; (5) identifying statistically enriched regions of coverage; (6) filtering likely artefactually enriched regions; (7) verifying the correct TF as a ChIP-Seq-specific QC. Each of these steps are applied uniformly to all samples and are described in more detail below.

**Read Quality Control.**
All samples began with an initial quality control check to ensure raw read quality with FastQC [12], which provided several metrics assessing the reads and underlying base calls should any troubleshooting be needed. One metric is the distribution of base quality scores, which are a measure of the probability that a base call is incorrect:

$$Quality\ score = -10 \log p \qquad (1)$$

where p is the probability of an incorrect base call. From equation 1, we can see that higher quality scores indicate a lower probability of an incorrect call. From FastQC, our median quality score over the first 5 bases was typically around 32, with a sharp increase to 36 or higher for the remainder of the read. For reference, a quality score of 30 indicates a 0.001 chance that the base call was incorrect, and thus scores of 30 or higher are typically considered high-quality bases.

**Read Trimming.**

Next, we removed Illumina adapters and low-quality base calls from the ends of our reads. We used Cutadapt [13] to clip adapters and base calls with a quality score below 30. However, as discussed above, all base calls typically scored above this threshold, and thus the primary function of this step was generating reads containing only *E. coli* sequences inside of the adapters.

**Alignment of Reads.**

Reads were aligned to the *E. coli* reference genome. Alignment was done with Bowtie2 [14] with default settings, and bam files were returned. These were converted to sam files with samtools [15] for downstream applications. We also converted the alignments to tdf format, which is compatible with most genome viewers, using IGVTools [16]. We also collected statistics on the alignments using Picard's CollectAlignmentSummaryMetrics and CollectWgsMetrics [17]. These included the number & fraction of reads that were aligned, the number of bases that were/were not covered by reads, how many bases were covered by 1, 5, 10, etc. reads, and more. Results were imported to our MySQL database.

**Calculating Coverage from Alignments.**

Next, we calculated the number of reads covering each base. We used our previously published peak calling tool, SPAT [18], to parse aligned reads into an Nx4 matrix (N = genome size), with columns for the genomic position, number of reads covering the position aligned to the forward strand, reverse strand, or either strand.

**Identifying Statistically Enriched Regions.**

We used the coverage matrices to identify enriched regions that may contain one or more TF binding sites. An enriched region is a contiguous sequence of DNA, at least 150bp long, that has statistically enriched coverage at each position compared to a background model. SPAT discovered enriched regions by fitting total genome-wide coverage to a lognormal distribution and identifying regions that with coverage in the top 0.1 percentile of this distribution for the whole sequence. We also required the maximum coverage in the region to be at least 4.75-fold enriched compared to the genomic average. This threshold was determined empirically by ROC analysis, where we used a set of RegulonDB [19] known binding sites as a set of positive regions and randomly selected genomic coordinates as negative regions. A custom python script flagged regions that did not pass the enrichment threshold to be omitted from downstream analyses.

**Filtering for Expected Peak Shift.**

ChIP-Seq produces a strand-specific signature of enrichment that can be used to identify true binding peaks [8]. When ChIP-DNA fragments are sequenced, reads are generated starting at the edges of fragments and moving inward, with the TF binding site(s) occurring somewhere in between. Reads will thus align to either side of the binding site in a strand-specific manner, where reads aligning to the forward strand will be shifted upstream of the actual binding site and reads aligning to the reverse strand will be shifted downstream. The expected distance between the forward and reverse strand profiles can be determined by the combination of DNA fragment size and NGS read length and measured by finding the distance that maximizes the cross-correlation between stranded enrichment. For our configuration (350 bp fragments after library preparation and 75 bp reads) the expected shift is 45 bases, thus enriched regions with shift < 45 were flagged and omitted from downstream analyses.

##### Filtering of Artifacts.

The pipeline filters out two broad categories of IP artifacts: (1) regions enriched in IP controls, and (2) regions enriched across experiments for large numbers of different TFs.

Filtering For Enrichment in Controls. For each promoter/tag configuration we performed multiple control experiments. For native tagging, we performed experiments with WT *E. coli* both with and without anti-tag antibody. For inducible tagging, we performed experiments with an empty vector with and without anti-tag antibody. For any given ChIP-Seq experiment, we filter out regions enriched in at least one control corresponding to that experiment type unless the region in the experiment was sufficiently enriched over the region in all control experiments. For native expression samples using the anti-FLAG antibody, all peaks <1.5-fold enriched over controls were filtered, for native expression samples using anti-V5 antibody, peaks <2.2-fold enriched were filtered, and for inducible experiments, peaks <2-fold enriched were filtered. Thresholds were determined through an ROC analysis relative to confirmed binding sites in RegulonDB.

Filtering for Regions Enriched in Multiple TFs. By performing ChIP-Seq on many TFs, we were able to identify regions that were enriched across many or all TFs (Table S3). In some cases, these regions were not enriched in controls and were therefore not detected by that filter (Figure S7). Of note, several of these regions are frequently reported in published ChIP-Seq experiments. An examination of these artifacts suggested that they can broadly be categorized as SPA tag-specific and tag-independent.

SPA tag specific artifacts consisted of 12 regions enriched in nearly all experiments using SPA-tags. An examination of these regions revealed that the 12 regions included all previously reported binding sites for the TF NagC, and all twelve regions were discovered in our ChIP-Seq of NagC. This suggested that these artifacts were somehow associated with binding and pull-down of NagC. We confirmed this in two ways. First, were performed experiments with SPA tags in strains deleted for NagC and the 12 regions did not appear enriched. Second, it is known that binding by NagC is prevented by Glc-Nac [20]. Experiments performed using SPA tags in WT *E. coli* with the addition of N-Acetyl-D-glucosamine (Glc-Nac) also removed these 12 regions. Based on these results, if a SPA experiment had enriched regions overlapping at least 5/12 of the canonical regions, all overlapping regions were flagged for filtering. This threshold was determined empirically by comparing the proportion of known sites flagged or not, and the proportion random sites flagged or not at different thresholds.

Tag-independent artifacts were classified as strong or weak depending on the number of different TFs for which the region was enriched. Strong artifacts were enriched for at least 50 TFs and for at least half of these experiments, the artifact was at least half as strong as the most enriched region. These regions were removed by flagging any enriched region that overlapped an identified artifact. Weak artifacts were enriched in more than 25 TFs but less than 50. Enriched regions overlapping these artifacts were filtered if the region was less than 7x enriched relative to the average across all other TFs.

##### Filtering Watermarks

Inducible ChIP-Seq uses a high-copy plasmid, causing genes on the plasmid to be present in high number compared to the single-genomic copy of other genes. As a result, plasmid genes will have background coverage 100s-1000s of fold-enriched compared to background, a signature we call the "watermark" (Figure S8). In addition, because AraC is present in the plasmid, this TF also displays a watermark.

We filter enriched regions that overlap these expected watermarks. After removing regions overlapping those genes, we applied a custom detection script that aimed to recover enriched regions that could represent autobinding, while avoiding the artefactual enrichment of the plasmid. We used the same approach to detect enriched regions, but only look in a 500 bp window around the edges of the tagged-TF that would have artefactual inducible enrichment. Positive matches are added back to the set of passing regions.

**Verifying the Correct Transcription Factor.**

We developed two custom QC scripts to determine that the correct TF was tagged in a ChIP-Seq experiment. One script is used on every experiment where we search the NGS reads for sequences matching our epitope tags to determine which genes bore a tag. For inducible ChIP-Seq, we performed the additional QC step that aimed to identify genes that were present on the inducible plasmid. In both cases, we require these stages to demonstrate that only one TF was unambiguously tagged.

TF Verification Via Epitope Tag. We searched reads in every sample for sequences from the beginning and end of our epitope tags to verify that the correct TF was tagged (Figure S9). Because we encoded the tag alongside the TF, some reads span the junctions around the epitope and read into gDNA. We searched reads for the first or last 25bp of the tag sequence, and for reads that matched we retrieved the side of the read that would have come from the genome (taking the 5' side of a forward-aligned read that found the first 25bp of the tag, or a reverse-aligned read that found the last 25bp; or taking the 3' side of a forward-aligned read with the last 25bp of the tag, or a reverse-aligned read finding the first 25bp). We required the retrieved read fragment to be at least 8bp to find its genomic locus, aligned fragments back to the reference genome with Bowtie2. We identified which genes (if any) the aligned reads overlapped using Bedtools2 intersect [21]. Successful experiments are marked by clear enrichment of the TF of interest, where nearly all identified reads align to the correct TF (or potentially the downstream gene for reads found from the end of the tag).

TF Verification Via Watermark Detection. We developed a custom script that aimed to identify if the correct TF displays a watermark (see above) (Figure S10). Potentially watermarked regions were those with at least 9x average genomic fold-enrichment for a consecutive 200bp. We filtered these hits, considering only those regions that covered at least 80% of a gene, and at most 80% of the enrichment was genic. Because AraC was present on the pBAD plasmid to enable inducibility, it should always be watermarked. For this case, we permitted up to 1/3 of the enriched region to be intergenic, as the araBAD promoter upstream of AraC was also present on the plasmid and thus part of its watermark. Finally, a passing inducible experiment was one where at most one TF and AraC are watermarked.

#### *Mathematical Model of TF-DNA Binding and ChIP Coverage*

##### Thermodynamic Model of Binding to a Single Site

To analyze TF-DNA binding, we seek to model the sequence specific binding energy of a TF to DNA. In the simplest case, we consider a system consisting of one sequence of interest and $N_{NS}$ non-specific sites. We then consider $N_{TF}$ transcription factors each of which can be either be bound to the sequence of interest with energy $\varepsilon_{seq}$ or occupy a non-specific site with energy $\varepsilon_{NS}$. By convention, more negative energies are more favorable. The nature of the non-specific sites depends on context, as we describe in more detail below. We further assume that occupancy of these states is in thermal equilibrium over the timescales of interest for our analysis, motivated by the fact that speed of binding reactions is very fast relative to the timescale of a typical ChIP experiment.

Given the assumption of thermal equilibrium, the probability that the sequence of interest will be bound by a TF can be calculated via the Boltzmann distribution [22]. This is accomplished by considering two states for the system: (1) an "unbound" state where all $N_{TF}$ molecules occupy "non-specific" sites, and (2) a "bound" state where $N_{TF} - 1$ TF molecules occupy non-specific sites, and one TF molecule is bound to the sequence of interest. Each state is associated with a multiplicity, Z, which is the number of configurations by which that state can be realized. For example, for the unbound state, $Z_{unbound}$ is the number of ways in which $N_{TF}$ TFs can be arranged in $N_{NS}$ non-specific sites.

$$Z_{unbound} = Z(N_{NS}, N_{TF}) = \frac{N_{NS}!}{N_{TF}!\,(N_{NS} - N_{TF})!} \quad (2)$$

Similarly, $Z_{bound}$=Z($N_{NS}$, $N_{TF}$-1), and describes the number of ways in which $N_{TF} - 1$ TFs can be arranged in $N_{NS}$ non-specific sites. For $N_{NS} >> N_{TF}$, this can be approximated as:

$$Z(N_{NS}, N_{TF}) \sim \frac{N_{NS}^{N_{TF}}}{N_{TF}!} \quad (3)$$

The energy of each configuration can be calculated by summing the energies of the $N_{TF}$ TFs in that configuration. For example, for any unbound configuration:

$$\varepsilon_{unbound} = N_{TF}\varepsilon_{NS} \quad (4)$$

and for any bound configuration:

$$\varepsilon_{bound} = (N_{TF} - 1)\varepsilon_{NS} + \varepsilon_{seq} \quad (5)$$

For each configuration, we then calculate the Boltzmann factor (also called statistical weight), defined as the exponential of the negative energy relative to $k_bT$ for that configuration where $k_b$ is the Boltzmann constant and T is temperature:

$$Boltzmann\ factor = e^{-\varepsilon/k_bT} \quad (6)$$

For the following, we assume that all energies are given in units of $k_bT$ so that this term will be omitted. The probability that the system will be in the bound state can then be calculated as the sum of the Boltzmann factors for all bound configurations divided by the sum of the Boltzmann factors for all configurations divided by the sum of the Boltzmann factors for all configurations:

$$P_{bound} = \frac{Z_{bound}e^{\varepsilon_{bound}}}{Z_{bound}e^{\varepsilon_{bound}} + Z_{unbound}e^{\varepsilon_{unbound}}} \quad (7)$$

(where, as noted above, we assume energies are in units of $k_bT$). The denominator of equation 6 is the partition function of the system. As shown previously [22, 23], for the simple system of one sequence and $N_{NS}$ non-specific sites, equation 7 can be simplified to:

$$P_{bound} = \frac{1}{1 + \frac{N_{NS}}{N_{TF}}e^{\Delta\varepsilon}} \quad (8)$$

Where $\Delta\varepsilon$ is defined as the difference in energy between a single TF bound to the specific sequence and the TF occupying a non-specific site:

$$\Delta\varepsilon = \varepsilon_{seq} - \varepsilon_{NS} \tag{9}$$

$\Delta\varepsilon$ can be interpreted as the energy released when a TF transitions from a non-specific site to the specific sequence, or conversely the energy that must be provided to dislodge a TF from the specific sequence to a non-specific site. From equation 8, more negative $\Delta\varepsilon$ increases $P_{bound}$ and thus reflects a sequence with higher affinity to the TF relative to non-specific sites.

**Relationship to Hill Function**

Equation 8 can also be related a unimolecular model of TF binding to a DNA sequence:

$$TF + DNA \underset{k_r}{\overset{k_f}{\rightleftharpoons}} TFDNA \tag{10}$$

Here, $k_f$ is the rate of complex association (TF + DNA binding to form TF-DNA), and $k_r$is the rate of complex dissociation (TF-DNA unbinding into TF + DNA). We can model dynamics of TF-DNA concentration using the law of mass action:

$$\frac{d[TFDNA]}{dt} = k_f[TF][DNA] - k_r[TFDNA] \tag{11}$$

At thermal equilibrium, the association & dissociation reactions are balanced, and thus equation 11 is 0, allowing us to rewrite it as:

$$k_f[TF]_{eq}[DNA]_{eq} = k_r[TFDNA]_{eq} \tag{12}$$

Separating the rate-constants and equilibrium concentrations to different sides allows us to recover the association/dissociation constants, $K_A$ & $K_D$:

$$\frac{[TF]_{eq}[DNA]_{eq}}{[TFDNA]_{eq}} = \frac{k_r}{k_f} = K_D = \frac{1}{K_A} \tag{13}$$

To determine the probability of a site being bound at equilibrium, we simply divide the amount of TF-DNA complex by all DNA:

$$P_{bound} = \frac{[TFDNA]_{eq}}{[DNA]_{eq} + [TFDNA]_{eq}} \tag{14}$$

If we multiply top and bottom by $1/[TFDNA]_{eq}$, we get:

$$P_{bound} = \frac{1}{\frac{[DNA]_{eq}}{[TFDNA]_{eq}} + 1} \tag{15}$$

Multiplying top and bottom by $[TF]_{eq}$ gives:

$$P_{bound} = \frac{[TF]_{eq}}{\frac{[TF]_{eq}[DNA]_{eq}}{[TFDNA]_{eq}} + [TF]_{eq}} \quad (16)$$

Where we can recognize the dissociation constant in the denominator, and we recover the familiar Hill function form for TF binding at equilibrium:

$$P_{bound} = \frac{[TF]_{eq}}{K_D + [TF]_{eq}} \quad (17)$$

#### Equilibrium Constant Relation to Free Energy

$K_D$ (and $K_A$) can in turn be related to the Gibbs free energy of the reaction. From thermodynamics, we know that the energy change of the reaction of Equation 10 can be determined by:

$$\Delta G = \Delta G^0 + N_A k_b T \ln\left(\frac{[TFDNA]}{[TF][DNA]}\right) \quad (18)$$

Where $\Delta G^0$ is the Gibbs free energy of associated with the reaction under standard conditions in which all reactants are present a 1M, $k_b$ is the Boltzmann constant, and $N_A$ is Avogadro's number or the number of molecules in 1 mole. When $\Delta G^0 = 0$, the system is at equilibrium, and thus:

$$\Delta G^0 = -N_A k_b T \ln\left(\frac{[TFDNA]_{eq}}{[TF]_{eq}[DNA]_{eq}}\right) \quad (19)$$

Where we can recognize the quantity inside the log as $K_A$, and rearrange to solve for it:

$$K_A = e^{\frac{-\Delta G^0}{N_A k_B T}} \quad (20)$$

or equivalently:

$$K_D = e^{\frac{\Delta G^0}{N_A k_B T}} \quad (21)$$

Equations 8 & 17 can be related by letting $[TF]_{eq} = \frac{N_{TF}}{N_{NS}}$ leading to:

$$K_D = e^{\Delta \varepsilon} \quad (22)$$

or equivalently:

$$K_A = e^{-\Delta \varepsilon} \quad (23)$$

Providing an interpretation of the equilibrium constants in terms of the change in energy (in units of $k_bT$) when a single TF molecule binds DNA.

Moreover, relating equations 21 and 22, we can recognize that:

$$\Delta\varepsilon = \frac{\Delta G^0}{N_A} \tag{24}$$

which indicates that the change in energy associated with TF binding per molecule (left) is the change in energy for 1 mole of molecules divided by the number of molecules per mole (right).

**Probability of TF Binding to a Sequence with Multiple Sites**

A more realistic model of TF-DNA binding assumes that each of $N_{region}$ positions in a DNA sequence (for example, a 101mer) can be considered as an individual binding site i, each with their own energy $\varepsilon_i$. As before, there are $N_{TF}$ proteins available to bind any site within this region, and $N_{NS}$ nonspecific sites that can be bound each with energy $\varepsilon_{NS}$. We further make the simplifying assumption that only 1 TF can bind the sequence at any time but can do so at any of the $N_{region}$ positions. Determining the probability of binding anywhere in this sequence requires deriving the appropriate Boltzmann factors and multiplicities for configurations where the sequence is bound by a TF or not. For configurations where the sequence is not bound, the state weight is:

$$\frac{N_{NS}!}{N_{TF}!\,(N_{NS}-N_{TF})!}e^{-N_{TF}*\varepsilon_{NS}} \tag{25}$$

For bound configurations, we sum the Boltzmann factors of a TF binding to each position in the region:

$$\frac{N_{NS}!}{(N_{TF}-1)!\,(N_{NS}-(N_{TF}-1))!}\sum_{i=1}^{N_{region}} e^{-\varepsilon_i-(N_{TF}-1)*\varepsilon_{NS}} \tag{26}$$

Then, the probability of a TF binding the sequence is:

$$P_{bound} = \frac{\frac{N_{NS}!}{(N_{TF}-1)!\,(N_{NS}-(N_{TF}-1))!}\sum_{i=1}^{N_{region}} e^{-\varepsilon_i-(N_{TF}-1)*\varepsilon_{NS}}}{\frac{N_{NS}!}{(N_{TF}-1)!\,(N_{NS}-(N_{TF}-1))!}\sum_{i=1}^{N_{region}} e^{-\varepsilon_i-(N_{TF}-1)*\varepsilon_{NS}} + \frac{N_{NS}!}{N_{TF}!\,(N_{NS}-N_{TF})!}e^{-N_{TF}\varepsilon_{NS}}} \tag{27}$$

Using the assumption that $N_{NS} >> N_{TF}$, we can simplify:

$$P_{bound} = \frac{\frac{N_{NS}^{(N_{TF}-1)}}{(N_{TF}-1)!}\sum_{i=1}^{N_{region}} e^{-\varepsilon_i-(N_{TF}-1)*\varepsilon_{NS}}}{\frac{N_{NS}^{(N_{TF}-1)}}{(N_{TF}-1)!}\sum_{i=1}^{N_{region}} e^{-\varepsilon_i-(N_{TF}-1)*\varepsilon_{NS}} + \frac{N_{NS}^{N_{TF}}}{N_{TF}!}\,e^{-N_{TF}\varepsilon_{NS}}} \tag{28}$$

We then multiply both the numerator and denominator by $(N_{TF}\text{-}1)!/N_{NS}^{N\_TF-1}$:

$$P_{bound} = \frac{\sum_{i=1}^{N_{region}} e^{-\varepsilon_i-(N_{TF}-1)*\varepsilon_{NS}}}{\sum_{i=1}^{N_{region}} e^{-\varepsilon_i-(N_{TF}-1)*\varepsilon_{NS}} + \frac{N_{NS}}{N_{TF}}\,e^{-N_{TF}\varepsilon_{NS}}} \tag{29}$$

and divide the sum $\sum_{i=1}^{N_{region}} e^{-\varepsilon_i-(N_{TF}-1)*\varepsilon_{NS}}$ from the top and bottom:

$$P_{bound} = \frac{1}{1 + \frac{N_{NS}}{N_{TF}} \frac{e^{-N_{TF}\varepsilon_{NS}}}{\sum_{i=1}^{N_{region}} e^{-\varepsilon_i - (N_{TF}-1)*\varepsilon_{NS}}}} \tag{30}$$

which is equivalent to:

$$P_{bound} = \frac{1}{1 + \frac{N_{NS}}{N_{TF}} \frac{1}{\sum_{i=1}^{N_{region}} e^{N_{TF}\varepsilon_{NS}} e^{-\varepsilon_i - (N_{TF}-1)*\varepsilon_{NS}}}} \tag{31}$$

which we can simplify as:

$$P_{bound} = \frac{1}{1 + \frac{N_{NS}}{N_{TF}} \frac{1}{\sum_{i=1}^{N_{region}} e^{-\varepsilon_i + \varepsilon_{NS}}}} \tag{32}$$

or

$$P_{bound} = \frac{1}{1 + \frac{N_{NS}}{N_{TF}} \frac{1}{\sum_{i=1}^{N_{region}} e^{\Delta\varepsilon_i}}} \tag{33}$$

where $\Delta\varepsilon_i = \varepsilon_i - \varepsilon_{NS}$ as in equation 9. From this, we can see that the probability of a single TF binding to a sequence with multiple binding sites is a function of the number of protein molecules ($N_{TF}$), and the sum of energy differences at each site (difference between binding the site vs a nonspecific site).

If we then equate these equations to equation 8 above and consider a single sequence with $\Delta\varepsilon_{eff}$, then we have:

$$e^{\Delta\varepsilon_{eff}} = \frac{1}{\sum_{i=1}^{N_{region}} e^{\Delta\varepsilon_i}} \tag{34}$$

Which implies an effective energy of binding to the whole sequence of length $N_{region}$ is related to the sum of energetic contributions to each individual site in the region:

$$\Delta\varepsilon_{eff} = -\log\left(\sum_{i=1}^{N_{region}} e^{-\Delta\varepsilon_i}\right) \tag{35}$$

**Probability a TF is Genome-Bound vs In Solution**

To estimate the probability of a TF being bound to genomic DNA relative to solution, we define two states: (1) a single TF is bound to one of $N_{genome}$ sites with energy $\varepsilon_{genome}$; and (2) a single TF is in one of $N_{solution}$ sites with energy $\varepsilon_{solution}$. By convention, binding energies are taken with solution as the reference, thus $\varepsilon_{solution}$ is 0. We seek to define the probability a TF is bound vs in solution, and start by defining the appropriate Boltzmann factors, for the case where TF is in solution:

$$\frac{N_{solution}!}{N_{solution}!\,(N_{solution}-1)!} \tag{36}$$

And the case where TF is bound to the genome:

$$\frac{N_{genome}!}{N_{genome}!\,\left(N_{genome}-1\right)!}e^{-\varepsilon_{genome}} \tag{37}$$

The probability that of being genome-bound is the ratio of its Boltzmann factor to the sum of all Boltzmann factors:

$$p_{bound} = \frac{\frac{N_{genome}!}{N_{genome}!\,\left(N_{genome}-1\right)!}e^{-\varepsilon_{genome}}}{\frac{N_{genome}!}{N_{genome}!\,\left(N_{genome}-1\right)!}e^{-\varepsilon_{genome}} + \frac{N_{solution}!}{N_{solution}!\,(N_{solution}-1)!}} \tag{38}$$

With the assumption that $N_{solution} >> N_{genome}$ we can simplify:

$$p_{bound} = \frac{N_{genome}e^{-\varepsilon_{genome}}}{N_{genome}e^{-\varepsilon_{genome}} + N_{solution}} \tag{39}$$

Divide the top and bottom by $N_{genome}e^{-\varepsilon genome}$:

$$p_{bound} = \frac{1}{1 + \frac{N_{solution}}{N_{genome}}e^{\varepsilon_{genome}}} \tag{40}$$

We see that the probability a single TF is bound to the genome is a function of the genome binding energy relative to solution, and the ratio of solution sites to genomic sites. For our analyses, we calculate the value for a range of ratios of solution sites to genome sites. To select one value of this ratio for discussion, we used a previously published measurement suggesting that genomic DNA accounts for roughly 1% of the volume of an *E. coli* cell [24]. Given that binding to DNA occurs on a roughly one-dimensional manifold while the solution space is a three dimensional volume, we take the cube of this percent and invert it to arrive at a rough estimate of $10^6$ solution sites for every genomic base.

#### *Motif Comparison*

To align PFMs, we maximized the pairwise information content from comparing all possible PFM alignments (including the forward- and reverse-complement of one motif). To compare PFMs from BoltzNet and RegulonDB v12.0 [19], we converted PFMs into information matrices using transform_matrix in logomaker [25]. Information at each base was normalized to 1, and only bases with a total information of at least 0.6 in at least one of the matrices were compares. Alignments were scored by taking the inner product of the information vectors at each aligned position and summing those inner products across aligned positions. In cases where alignments included overhangs, edges were padded with 0.0001. The alignment maximizing the sum of information inner products was returned as the best alignment.

#### *Library-ChIP*

We employed Library-ChIP following a previously published method [26] (Figure S21), which allowed us to perform ChIP on designed binding sites in high-throughput. The protocol consists of the following steps: (1) library construction and transformation; (2) growing cells and preparing for IP; (3) immunoprecipitation of post-IP sample; (4) purification of pre- and post-IP DNA; (5) library prep and pooling; and (6) Library-ChIP analysis. These steps are detailed below.

##### Library-ChIP Library Construction

We cloned designed binding sites into a single copy pAMD plasmid (Figure S22) to create a plasmid library where each plasmid had a single designed site. These could be transformed into our tagged TF strains to assay binding via Library-ChIP.

We ordered an oligo-pool from IDT that contained over 100 novel binding sites (Table S5) designed with varied specificities toward PdhR, AllR, and GlnG, along with 3 genomic reference sites for each TF (2 in the case of GlnG). Each site was designed as a 61-mer, and 20 bp were added to each end with homology to the pAMD insertion site, creating 101-mer oligos. The oligo-pool was received as ssDNA and a universal primer pair we designed was used to create dsDNA with PCR (Table S4 – oPool dsDNA oligos). We mixed 1ng oPool DNA with 12.5μL KAPA HiFi HotStart ReadyMix (Roche Cat. #07958935001) and 0.75μL of each primer at 10μM, then filled the reaction to 25μL with sterile water. We performed PCR with a 3min initial denaturation at 95°C, followed by 5 PCR cycles with a 98°C denaturation for 20 seconds, 15 second annealing at 62°C (based on primer $T_m$), and a 15 second extension at 72°C. After the final PCR cycle, we did a 1-minute extension at 72°C before cooling to 4°C. Reactions were cleaned up with Qiagen's PCR Purification Kit (Qiagen #28104).

To facilitate cloning, pAMD was linearized with BamHI and HindIII restriction enzymes from NEB (Cat #'s R0136 & R0104, respectively) and then ligated with the dsDNA pool using NEBuilder HiFi DNA Assembly Kit (#E2621). Ligation was transformed into *E. coli* EPI300 received from the Wade Lab via electroporation and selected with carbenicillin. 12 colonies were selected for Sanger sequencing to confirm that different sequences from the pool had been inserted (Table S4 – oPool Test Primers). To ensure that each designed sequence is present in the pool, we aimed to get ~10X the number of colonies as there were binding sites in the pool, in this case, we targeted ~1200 colonies. These were scraped from plates and grown in 50mL LB at 37°C with shaking at 250rpm and carbenicillin selection. pAMD replication was induced in the EPI300 cells with arabinose and plasmid was extracted with Qiagen's Miniprep Kit (#27104). The resulting plasmid pool was electroporated into a natively V5-tagged AllR strain, an inducible GlnG strain, and native SPA-tagged and inducible PdhR strains for Library-ChIP.

##### Growing Cells and Preparing for Immunoprecipitation

Library-ChIP started identically to our ChIP-Seq protocol until the addition of appropriate primary antibody. Briefly, we grew cells from glycerol stocks and subcultured into 50mL fresh media, culturing until log-phase at which point arabinose was added for induction if necessary (note: arabinose does not induce pAMD replication in our tagged-TF strains – thus pAMD remains single-copy under TF induction). We crosslinked cells and stopped crosslinking with glycine, then pelleted and washed cells twice in PBS. After the second wash, we resuspended cells in Buffer 1+PI, and sheared via sonication with Covaris S2. At this point, we pelleted the lysate and reserved ~10% of the DNA-containing supernatant as a reference to quantify the abundance of designed sequences before IP.

**Immunoprecipitation**

The remaining 90% of lysate continued with immunoprecipitation exactly as previously described. Briefly, the lysate was salt-adjusted to 10mM Tris-HCl pH8, 150mM NaCl, and 0.1% NP-40, 5μL of appropriate primary antibody was added, and samples were rocked at 4°C overnight. The following day, we performed secondary antibody binding with Protein G Agarose beads, utilizing the same 7-wash setup used in ChIP-Seq. Following the 7$^{th}$ wash, DNA was eluted once with EBB and once with TE + 1% SDS, and the two elutions were pooled.

**Purification of Pre- and Post-IP DNA**

Proteins were digested from pre- and post-IP DNA samples by adding Proteinase K at a 1:20 ratio, incubating for 1hr at 37°C, followed by proteinase deactivation at 65°C overnight. DNAs were purified from this reaction using Qiagen's PCR Purification Kit.

**Library-ChIP Library Prep and Pooling**

To prepare Library-ChIP DNA for NGS, we employed a 2-step PCR process following Illumina's 16S Metagenomic Library Preparation (Illumina Ref #15044223). The first PCR used primers targeted the flanking regions that were common across all binding sites in the pool (Table S4 – oPool Universal PCR Primers). The primers also contained overhangs that could be targeted by Illumina multiplexing primers. Briefly, we added 2.5μL of sample DNA (normalized to 5ng/μL if the concentration was higher), along with 5μL each of 1μM Forward & Reverse primer, and 12.5μL KAPA HiFi HotStart ReadyMix. PCR used 25 cycles, and the reaction was cleaned up with Qiagen's PCR Purification Kit (#28104).

We performed a second PCR to barcode fragments into NGS-compatible libraries using primers from Illumina's Nextera XT Index Kit v2 (Illumina Cat. #FC-131-2001). This time, we used 25μL KAPA HiFi HotStart ReadyMix, along with 5μL of cleaned up DNA from the first PCR, 10μL sterile water, and 5μl each of Nextera Kit Index 1 & Index 2 primers, ensuring each sample received a unique pair of primers. This PCR utilized 8 cycles and libraries were cleaned up with AMPure XP beads using single-side selection to remove any adapter-dimers.

We used the Bioanalyzer 2100 to confirm fragment length distributions and obtain library molarites as we did for ChIP-Seq. However, because Library-ChIP libraries are much less complex than ChIP-Seq (Library-ChIP libraries are a collection of designed binding sites with high similarity while ChIP-Seq surveys a genome on the scale of megabases), care should be taken to ensure that Library-ChIP libraries do not dominate the overall composition of a pool, as this can lead to sequencing artifacts. Some applications may require spiking in exogenous DNA, though this was not needed in our case. Libraries were sequenced on a NextSeq2000 with 150bp paired-end reads to ensure that reads reached from the adapters across the designed sequence.

**Analysis**

We quantified the abundance of designed sequences in pre- and post-IP samples to determine the enrichment of each designed site in Library-ChIP. First, we created a database of designed 61mers for each of the 3 TFs. We developed a custom script which searched Library-ChIP reads for a 20 bp anchor sequence present at the end of the insertion site, and common among all sequences in the pool. Upon a successful match, we took the next 37 bases after the anchor, which was sufficient to cover all variant nucleotides in our pool. We recorded the sequence of each 37-mer, and their abundance in the raw reads of pre- and post-IP samples. Then, we searched for the 37-mers in each of the designed variants and marked any identified matches. In some cases, a sequence could be present in more than one 37-mer, in which case those counts were merged. We filtered out any sequences present in fewer than 0.1% of the raw reads, and then normalized the abundance of each sequence within the sample by the average number of sequence counts. Finally, the ratio of normalized post-IP:pre-IP counts is presented as the Library-ChIP enrichment.

#### *Biolayer Interferometry*

We used an optimized version of our previously published BLI protocol [27, 28]. This assay allowed us to derive binding energies for our TF-DNA pairs by fitting to the 1:1 binding model used in the Octet Analysis software, which is a particular application of the unimolecular TF-DNA binding reaction in equation 10. The steps involved in BLI are as follows: (1) annealing dsDNA; (2) preparing and plating buffered TF and DNA solutions; (3) running the BLI assay; and (4) analyzing the data. These are described in more detail below.

##### Generation of Double-Stranded Biotinylated DNA

We ordered biotinylated-forward and corresponding reverse oligo to measure TF-DNA binding kinetics in BLI (Table S6). Oligos were ordered from IDT and resuspended to 100μM in 0.1X TE. We annealed oligos by combining the forward and reverse strands to 10μM in annealing buffer (10mM Tris-HCl, 50mM NaCl) in 1.5mL tubes and incubated at 95°C for 5min. Tubes were removed and allowed to cool to room temperature for annealing.

##### Preparing Solutions and Assay Plate

We used a common binding buffer (20mM Tris-HCl, 0.1mM EDTA, 10mM $MgCl_2$, 1mM DTT, 120mM KCl, 5% glycerol, and 0.05% Tween-20) for all assays. PdhR was previously reported to be functional in this buffer [29], and the other TFs tested empirically worked well in it. We used the biotinylated-dsDNA as the load molecule, becoming immobilized on the streptavidinized biosensor probe. Load solutions were made by mixing biotinylated-dsDNA at a 1:1 ratio with biotin for a final concentration of 10nM in binding buffer based on suggestions in [30] that spiking in biotin can mitigate potential overloading of BLI probes that confounds analysis.

We used different TF concentrations to measure affinity to DNAs that were or were not designed to have affinity toward the TF. For specific TF-DNA pairs, we determined the maximum TF concentration empirically by finding the concentration where the maximum and ½ the maximum TF concentration equilibrate near the same value. This was found to be 50nM for AllR, GlnG, Nac, UlaR, and 100nM for PdhR. Subsequent experiments assayed a 2-fold serial dilution of TF concentrations below the maximum. For nonspecific TF-DNA pairs, more TF was needed to observe a signal, and we uniformly used 100nM, 250nM, 500nM, 750nM, and 1μM concentrations in those cases.

We prepared a soak plate for probe preincubation by adding 200μL binding buffer to each well of a Greiner 96W Plate, PP, Flat Bottom, Chimney Style, Black (#655209) that would contain an assay probe. The assay plate was prepared by adding 80μL of solutions to a Greiner 384W Plate, PP, Flat Bottom, Black (#781209). Each probe required 3 wells in the assay plate: (1) buffer for baselining and dissociation; (2) loading well with biotin and biotinylated-dsDNA; (3) analyte well with or without TF. For each TF-DNA pair, we tested 6 TF concentrations using 8 probes. One probe was used as a “reference sensor”, which sees buffer wells for the entire assay and allows subtraction of any background signal. Another probe was used as a “reference sample”, which has DNA loaded onto it but sees no TF and allows subtraction of the effect of DNA loading. The remaining 6 probes assayed the response of different TF concentrations to the same concentration of DNA.

#### Running BLI

Each BLI assay measured the binding of a TF-DNA pair over 5 steps: (1) initial baseline; (2) DNA loading; (3) second baseline; (4) TF association; and (5) TF dissociation. All steps were performed on an Octet Red 384 at 30°C with shaking at 1000rpm using SA-tips (Sartorius #18-5020). Before a group of assays began, probes were allowed to soak in binding buffer for a 10-minute pre-incubation. Assays began with a 60 second baseline step, where probes read from buffer wells. Next, probes were moved into wells containing biotinylated DNAs, which were loaded onto the probe's surface for 2 minutes. A second baseline was taken in buffer for 60 seconds. Probes were then moved into TF-containing wells for 10 minutes, allowing TF to associate with DNA. This step was optimized such that enough time was given for the 2 highest TF concentrations to reach equilibrium. Finally, probes were moved back to buffer wells with no TF for 15 minutes, and TF-DNA complex dissociation was recorded. This step was also optimized such that enough time was given to observe dissociation of all TF concentrations.

#### Analyzing BLI

Our BLI analysis followed that of the 1:1 Binding Model used in the Octet Analysis software, which we implemented in MATLAB to determine binding energies for our designed sites. This model describes the BLI assay as a unimolecular binding reaction. When solved analytically using the assay parameters as initial conditions for association and dissociation, this model gives rise to one equation per phase that allows simultaneous fitting of association and dissociation, which we use to derive binding energies.

In association, we have:

$$y_A(t) = \frac{R_{max}}{(\frac{K_D}{[TF]_0} + 1)}(1 - e^{-(k_f[TF]_0 + k_r)t}) \quad (41)$$

Where $R_{max}$ is the maximum response in BLI across all traces for a given TF, $k_f$ is the rate of TF and DNA binding, $k_r$ is the rate of TF-DNA unbinding, $K_D$ is the dissociation constant (equation 13), and $[TF]_0$ is the initial concentration of TF for a given trace.

In dissociation, we have:

$$y_D(t) = Y_A e^{-k_r(t - t_A)} \quad (42)$$

Where $t_A$ is the time at the end of association, and $Y_A$ is the signal at the end of association, given by:

$$Y_A(t) = \frac{R_{max}}{(\frac{K_D}{[TF]_0} + 1)}(1 - e^{-(k_f[TF]_0 + k_r)t_A}) \quad (43)$$

Taken together, we have a pair of equations that completely describes the association and dissociation phases of our BLI assay.

To fit our data to these equations, we started by processing raw BLI traces. Reference sensor and sample traces were subtracted from all traces where TF was present, which removed background noise and the effect of DNA loading from the signal. The beginning of the association phase was zeroed to the average value from the last second of the pre-association baseline. The beginning of the dissociation phase was aligned to the average value from the last second of the association phase. Processed traces were fit to equations 41 and 42. Resulting kinetic constants were converted to equilibrium constants through equation 13, and to binding energies through equations 22 and 23 (Table S7).

### Supplementary Figures

3xFLAG atgGCCTACAGC..CAACGAAAGAATggcggtggcGACTACAAAGAC...TGAGCTCCCAAC

SPA atgGCCTACAGC..CAACGAAAGAATtccatggaaAAGAGAAGATGG...GATATTCATATG

V5 atgGCCTACAGC..CAACGAAAGAATggcggtggcgacGGTAAACCTATT...GATTCAACATAG

TF Sequence Minus Stop Codon | Linker | Tag Sequence

**Figure S1: Examples of TF Tags.** Tagged TF strains were generated by cloning tag sequences onto the C terminal of a TF's sequence, omitting its stop codon (green). Tags contained a short linker sequence (black) between the TF and tag sequence (orange).

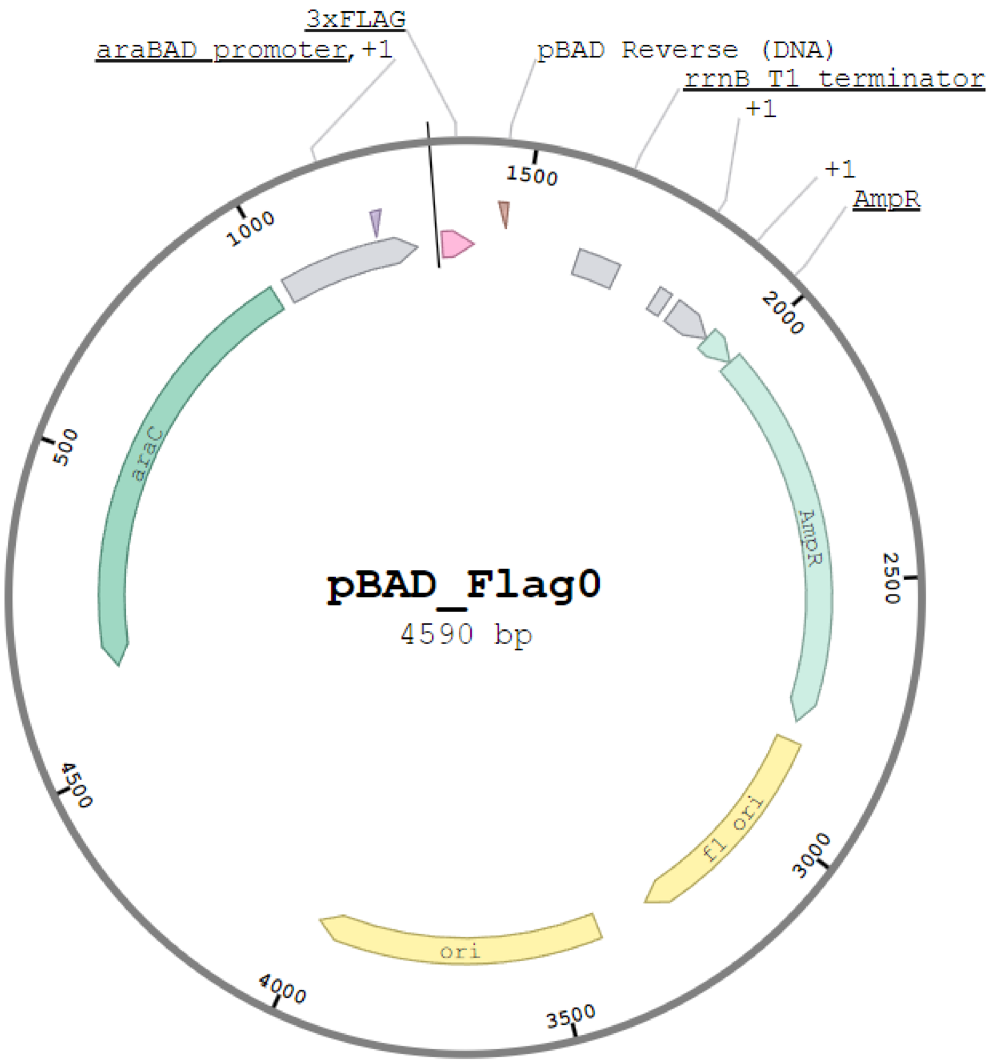

**Figure S2: pBAD24 Plasmid Map.** Our inducible ChIP-Seq experiments are facilited by cloning different TFs into the pBAD24 plasmid between the araBAD promoter and a 3xFLAG epitope tag. Successful clones can be selected for resistance to ampicillin due to the plasmid's AmpR gene. Tagged-TF expression can then be induced using arabinose which causes AraC to drive transcription from the araBAD promoter.

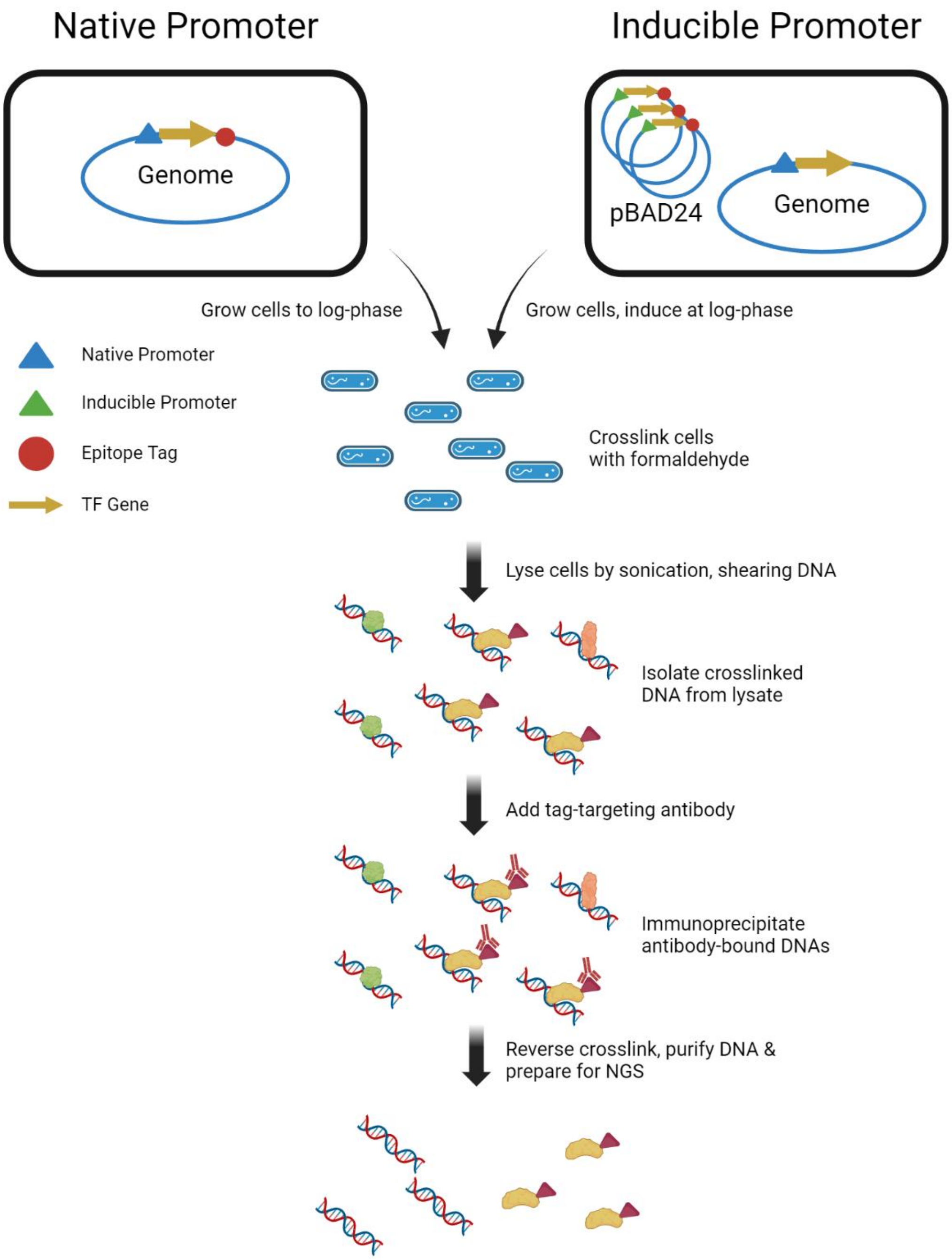

**Figure S3: ChIP-Seq Experiment Schematic.** We started by growing cells to logarithmic growth, at which point we may have added treatments (e.g. arabinose) as necessary. We crosslinked cells to fix TFs to any DNA sequences they were bound to, and lysed the cells via sonication, shearing the DNA at the same time. We isolated DNA fragments and added a tag-specific antibody to allow immunoprecipitation of fragments bound by the TF of interest. These fragments are purified and turned into NGS libraries for Illumina sequencing.

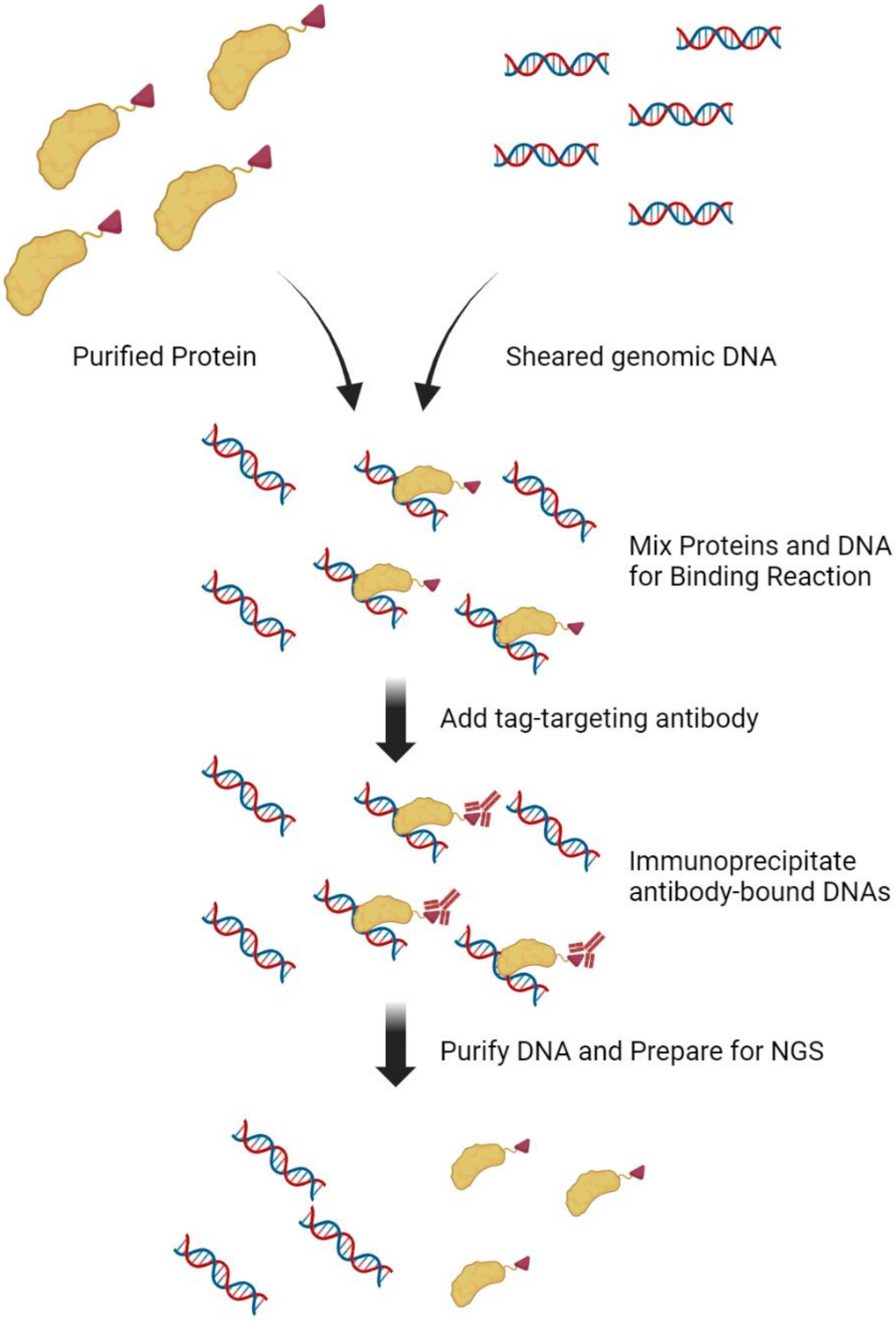

**Figure S4:** ***In Vitro* ChIP-Seq Experiment Schematic.** We started by combining a purified tagged-TF with pre-sheared genomic DNA. Next, we added a tag-specific antibody to immunoprecipitate DNA fragments bound by the TF. These fragments are purified and turned into NGS libraries for Illumina sequencing.

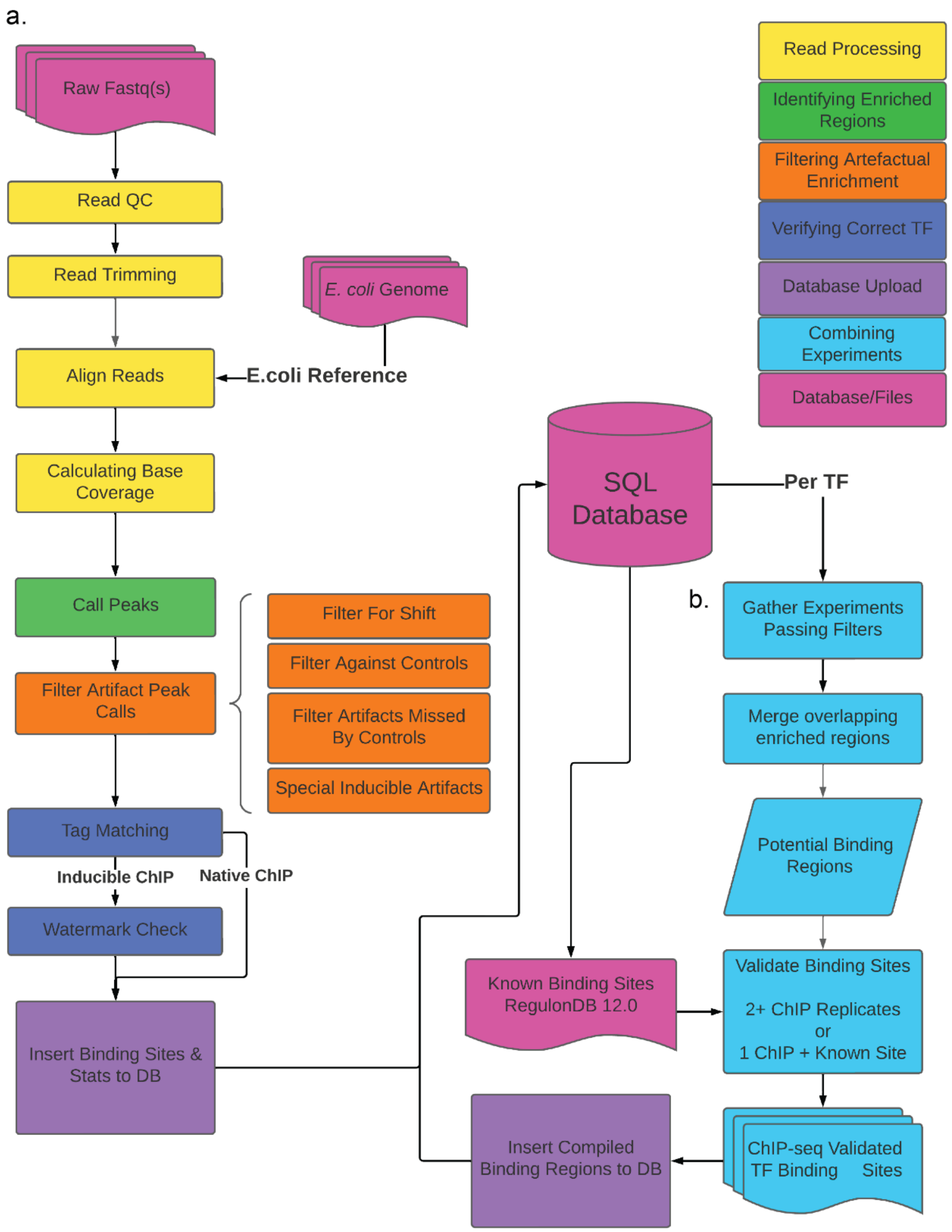

**Figure S5: ChIP-Seq Analysis Workflow.** For each experiment we uniformly processed our NGS reads to call high-confidence enriched regions (a). We began by assessing read quality and trimming low-quality bases and Illumina adapters from reads. We aligned reads to the *E. coli* genome and calculated the number of reads aligning to each strand at each base. We identified enriched regions and filtered likely artefactual enrichment. Finally, we performed custom QC to verify that the correct TF was tagged. All analysis results are imported to our MySQL Database. After all TF ChIP samples have been analyzed, we combine enriched regions across a TF's replicates to identify high-confidence binding regions (b).

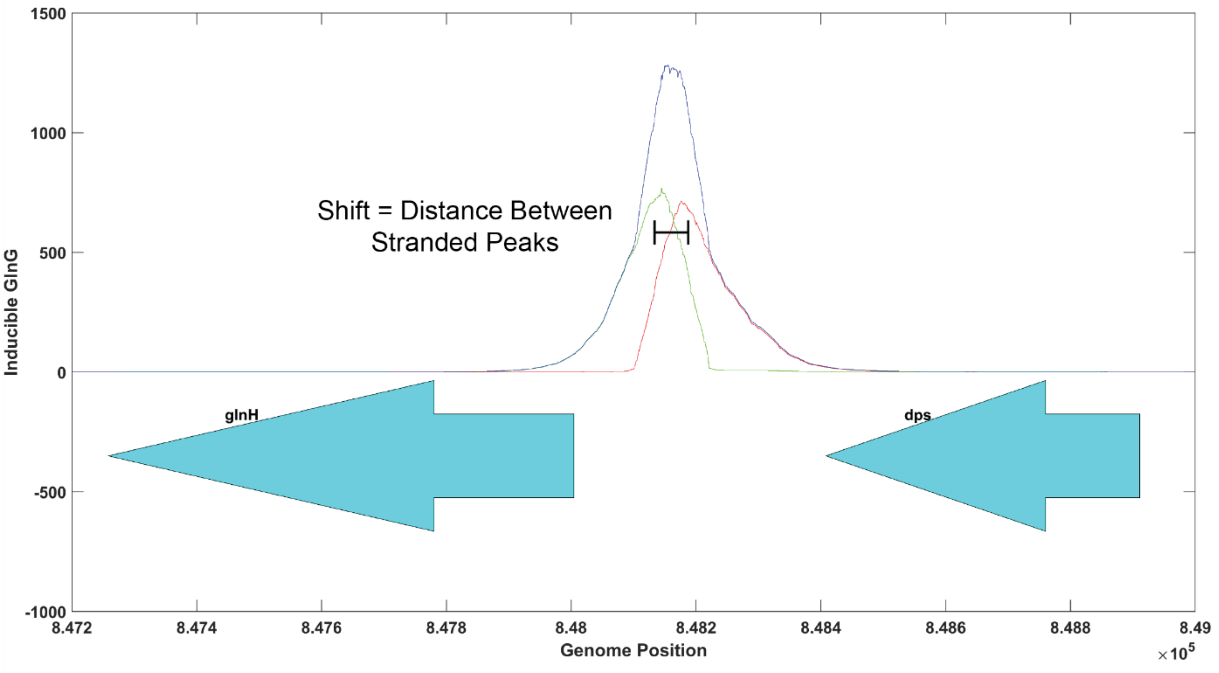

**Figure S6: Peak Shift Example.** Enriched regions that are a result of TFs binding point sources will appear with a characteristic shift between forward (green) and reverse (red) strand coverage as a consequence of sequencing bound fragments from the outside-in.

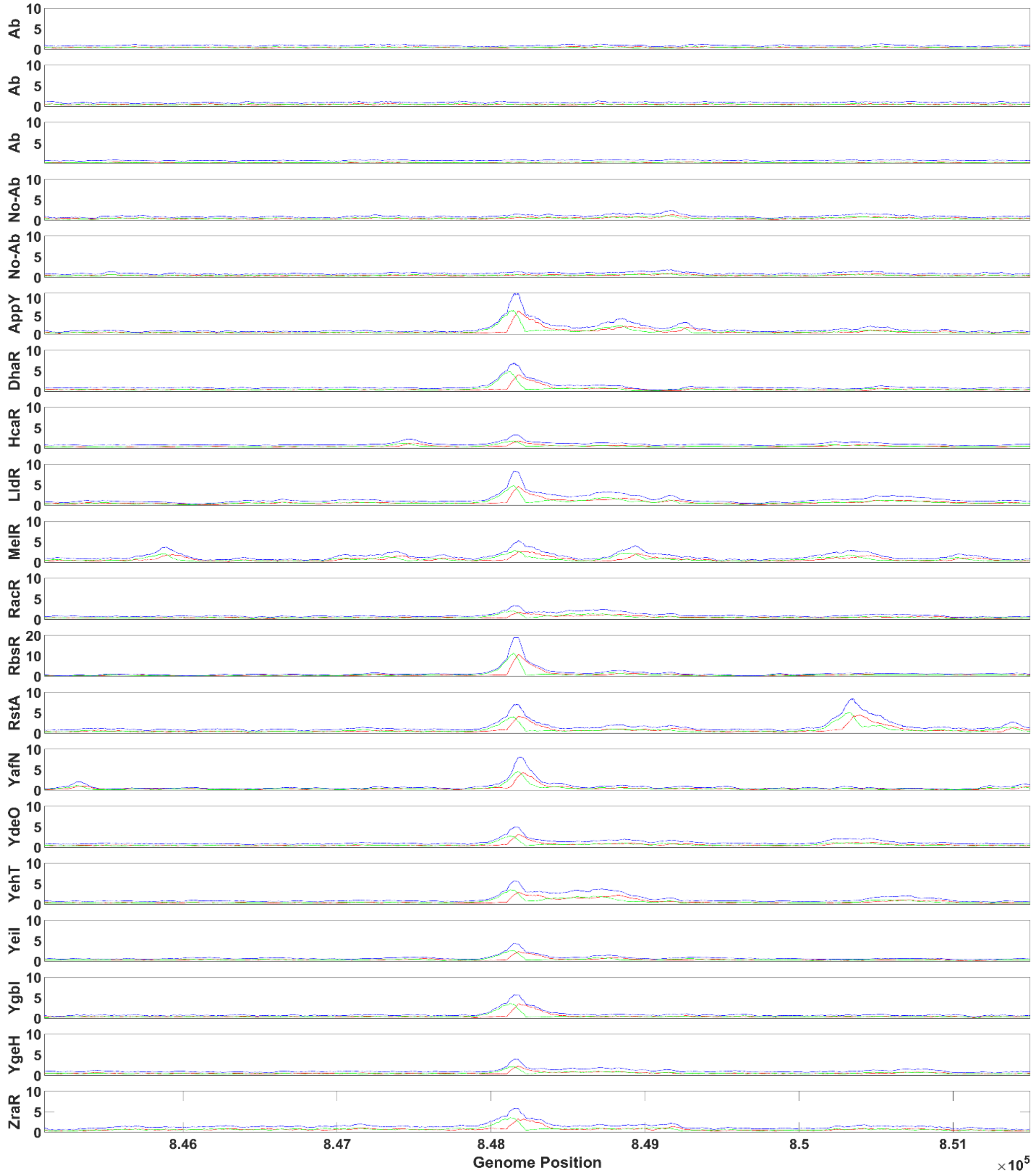

**Figure S7: ChIP-Seq Artifacts are not Always Apparent in Controls.** We have identified several genomic regions that appear to be artifacts on account of their ubiquity across TF ChIP-Seq datasets yet are not revealed in control experiments. The first 5 rows show different control experiments we performed, where this artifact region is not found. The following 15 tracks show a sample of different TFs where this region nearly always appears enriched and is therefore artefactual.

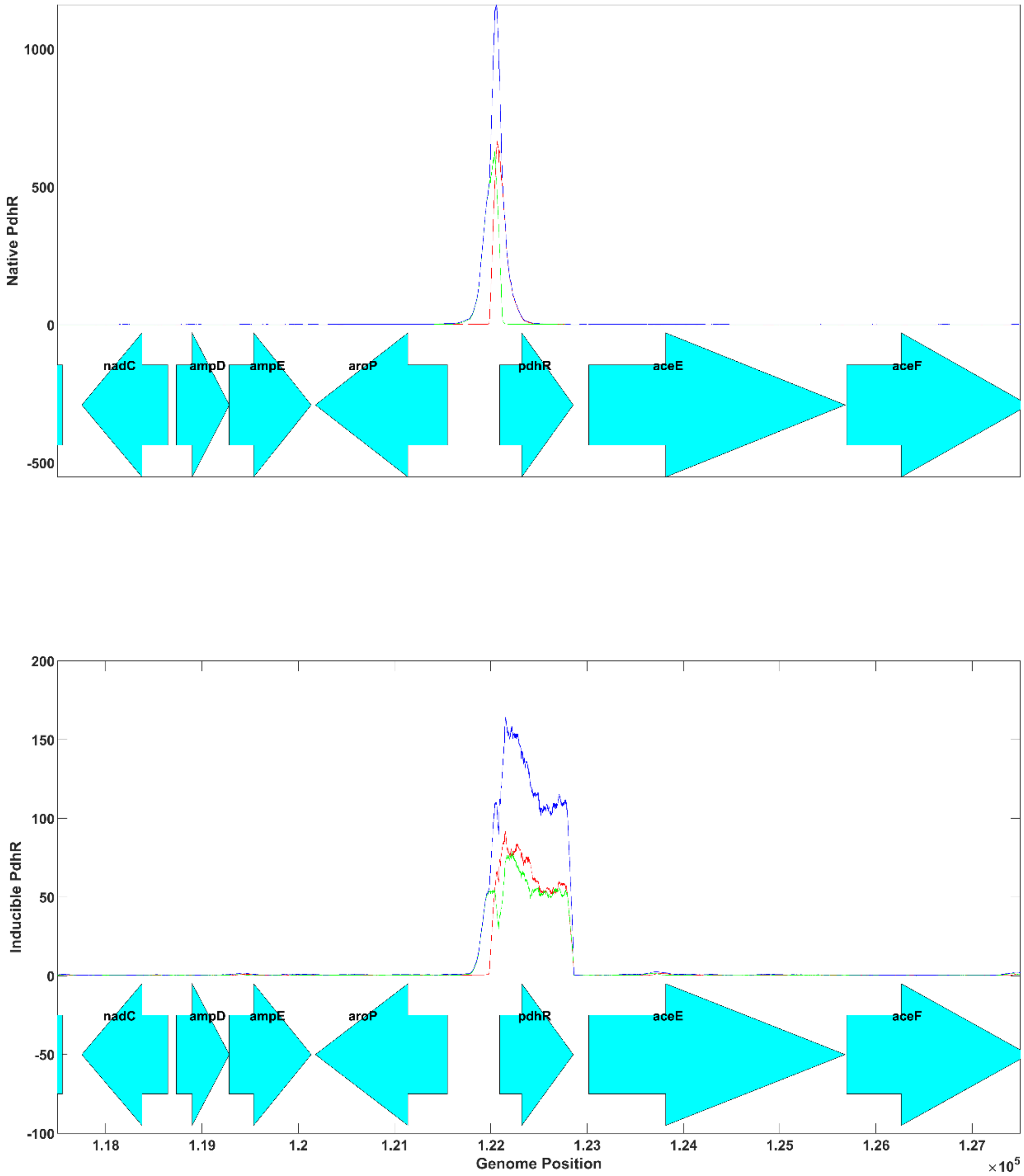

**Figure S8: High-Copy Plasmids Create Watermarks.** Coverage plot over PdhR's genomic locus showing a natively tagged experiment (top), and inducible experiment (bottom). Because of the high copy pBAD24 plasmid, we see significant enrichment across the entire PdhR sequence that is not present in the native experiment and can tarnish the appearance of the autobinding region seen clearly in the native experiment.

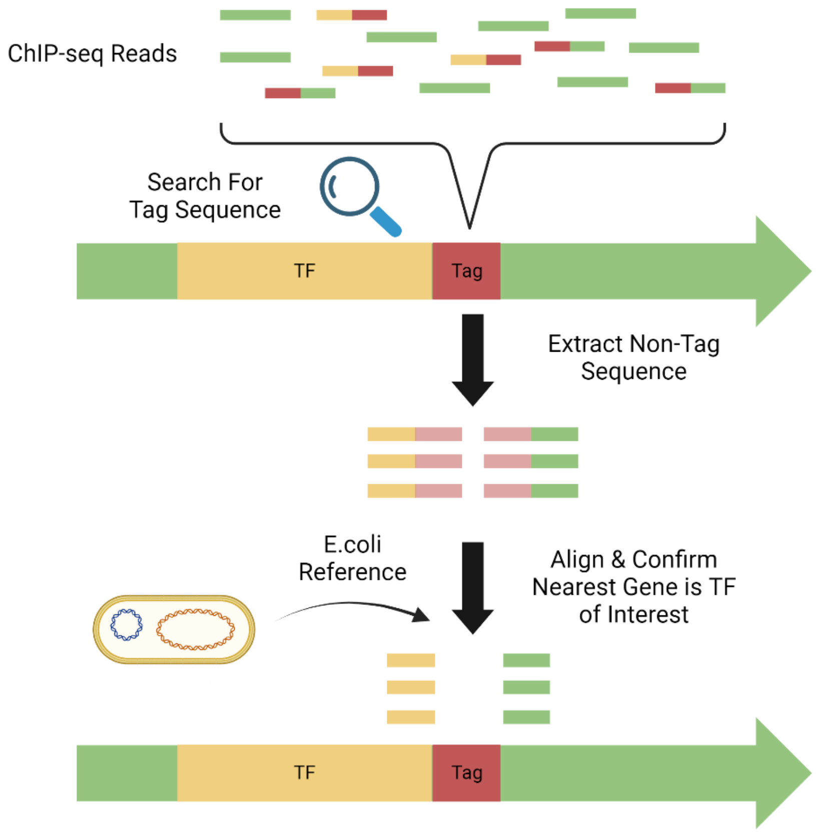

**Figure S9: Tag Matching Schematic.** We search raw ChIP-Seq reads for sequences matching the epitope tag cloned onto the TF's C terminus. The fragment of the read that does not match the epitope tag is aligned to the *E. coli* genome and any genes that the fragment overlaps are identified. For successful ChIP-Seq, we expect to see nearly all matched reads aligning at the end of the TF of interest.

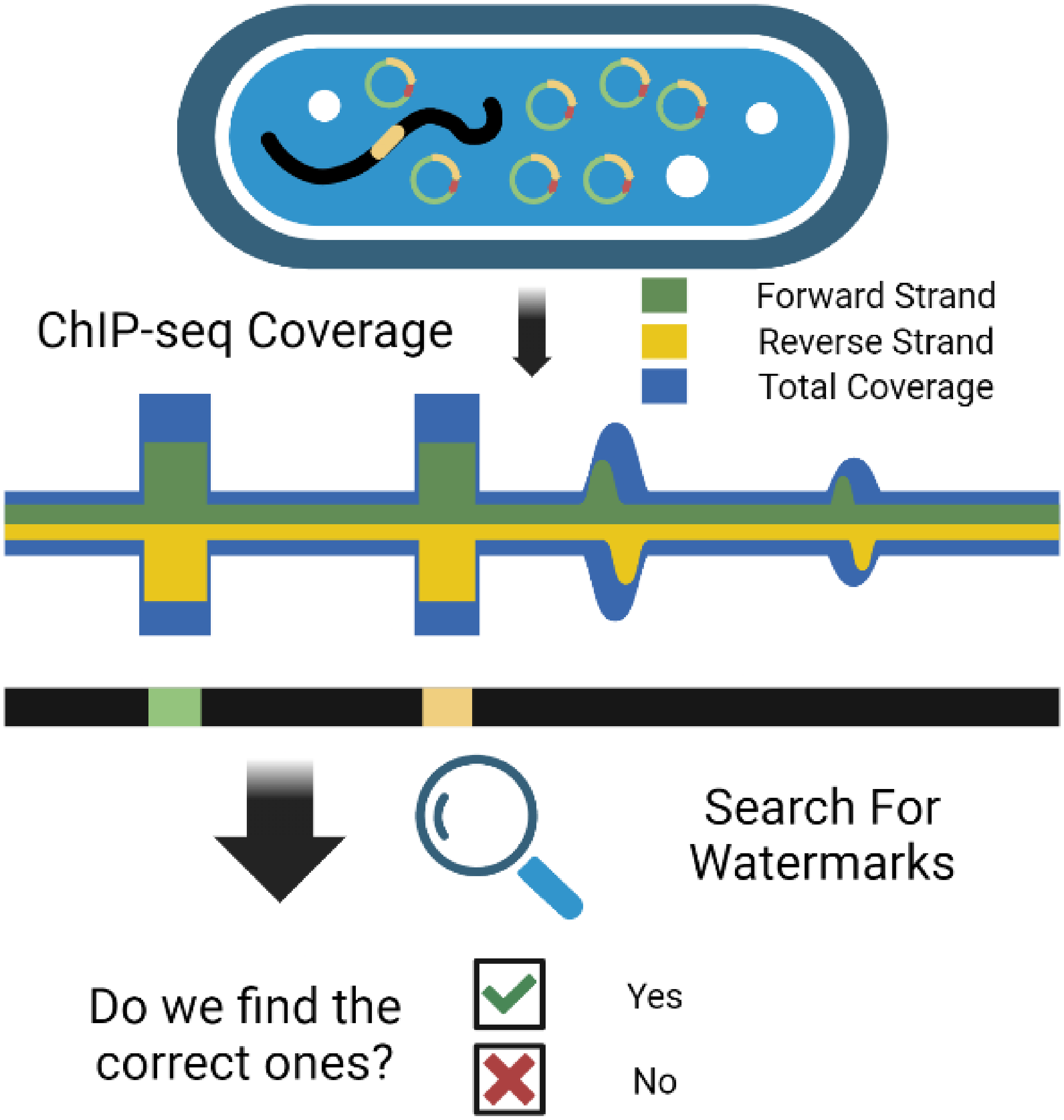

**Figure S10: Watermark Detection Schematic.** We perform QC on inducible ChIP-Seq experiments by searching for signatures of "watermarks". These appear visually as smears of coverage over the coordinates of genes on the inducible plasmid and are unlike ChIP-Seq peaks in two ways: 1) watermarks lack the peak shift characteristic of a real binding sites, and 2) coverage is enriched consistently across the length of the gene, often with much higher enrichment than ChIP-Seq peaks. Because our plasmids contain AraC (for induction) and the TF of interest, we consider a successful ChIP-Seq experiment one where at most these two watermarks are detected.

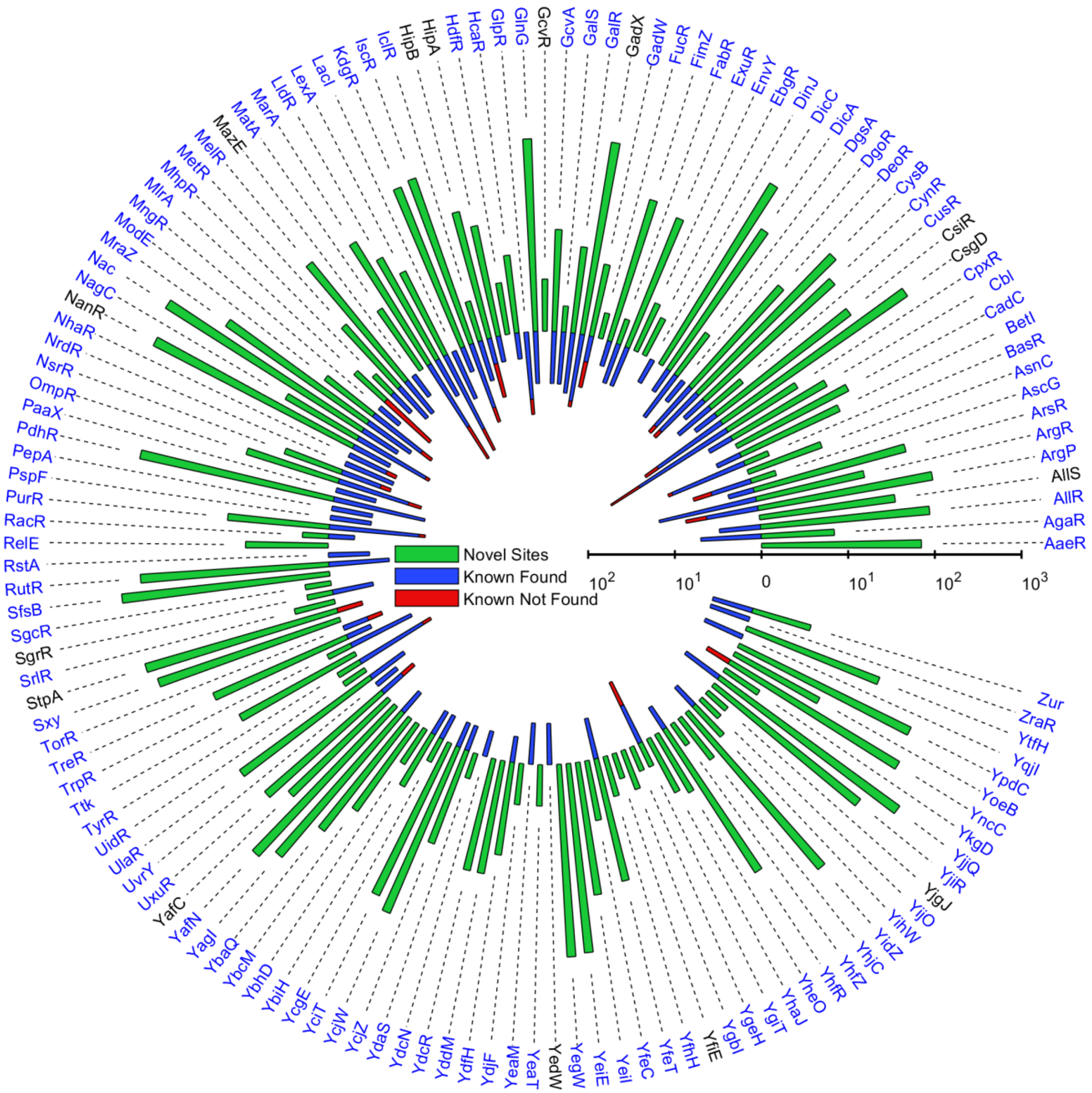

**Figure S11: Summary of TF Binding Sites Ordered by TF Symbol.** Plot displays the number of novel (green) sites and known sites (found: blue, not found: red) for 139 mapped TFs. Displays the same data as main Figure 1a, but the bars are sorted alphabetically by TF name.

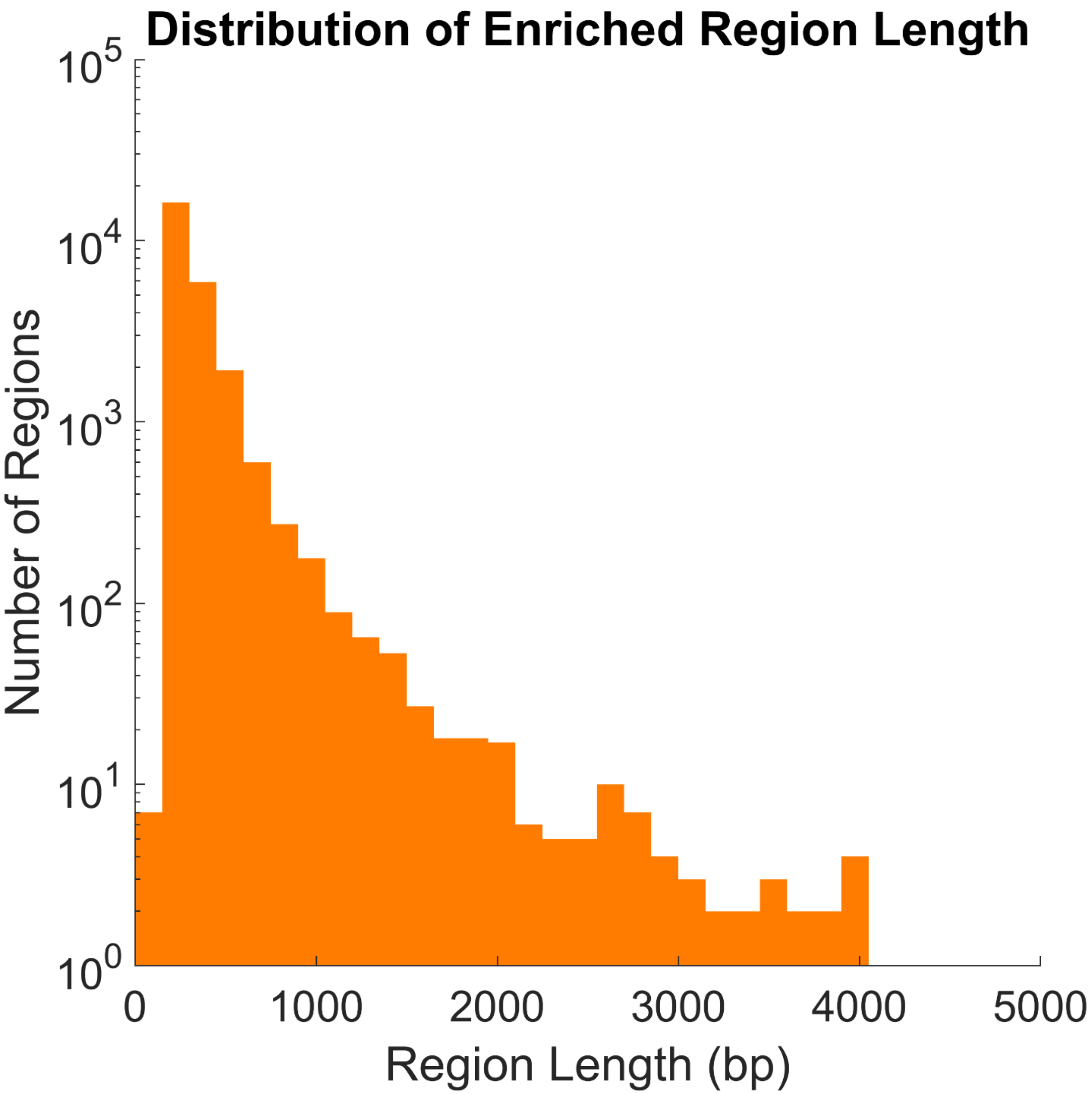

**Figure S12: Distribution of Enriched Region Length**. Histogram of enriched region lengths, with frequencies shown on the y-axis in log-scale. With the exception of a few special cases near the TF of interest in inducible ChIP-Seq, we required enriched regions to be at least 150 bp in length. The vast majority of regions were 500 bp or shorter (~90%).

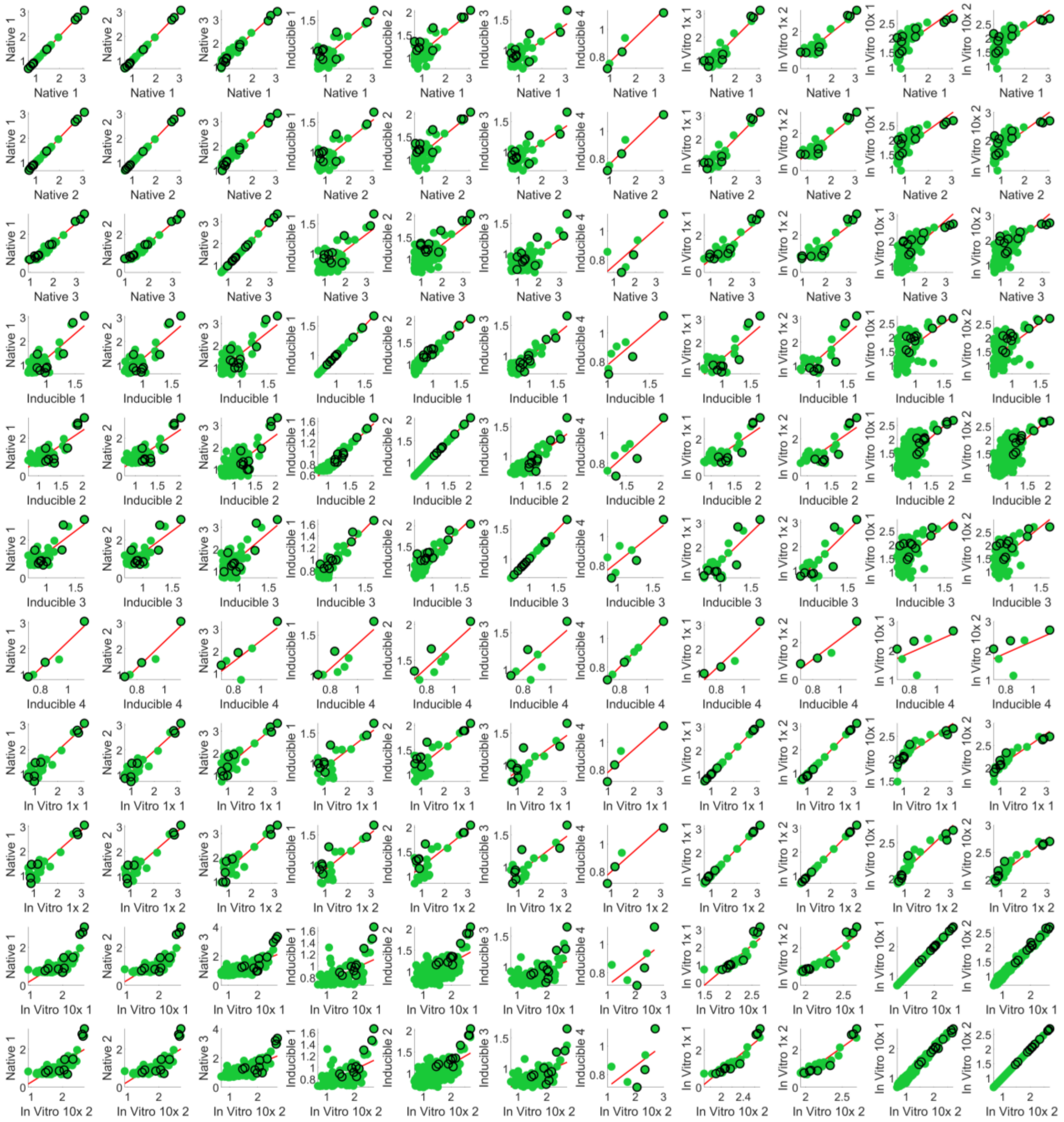

**Figure S13: Enrichment Scatterplots of All PdhR Replicate Comparisons.** Across 11 experiments performed *in vivo* (native or inducible) and *in vitro*, we see close agreement even across expression modes (median $R^2$ = 0.65). Between technical replicates of the same mode, we see exceptional agreement (median native $R^2$ = 0.98, median inducible $R^2$ = 0.76, median *in vitro* $R^2$ = 0.88). We also see the effect of saturation as a function of TF levels, where binding sites found in low TF-level (e.g. native) experiments have enrichment closer to the strongest binding site, and weaker sites not found with low-TF are revealed. This can be seen clearly with *in vitro* ChIP-Seq, where experiments with 10x TF find all of the sites from the 1x experiments with a tight range of enrichment, as well as a large number of sites that were lowly-enriched (and not identified) in the 1x experiments.

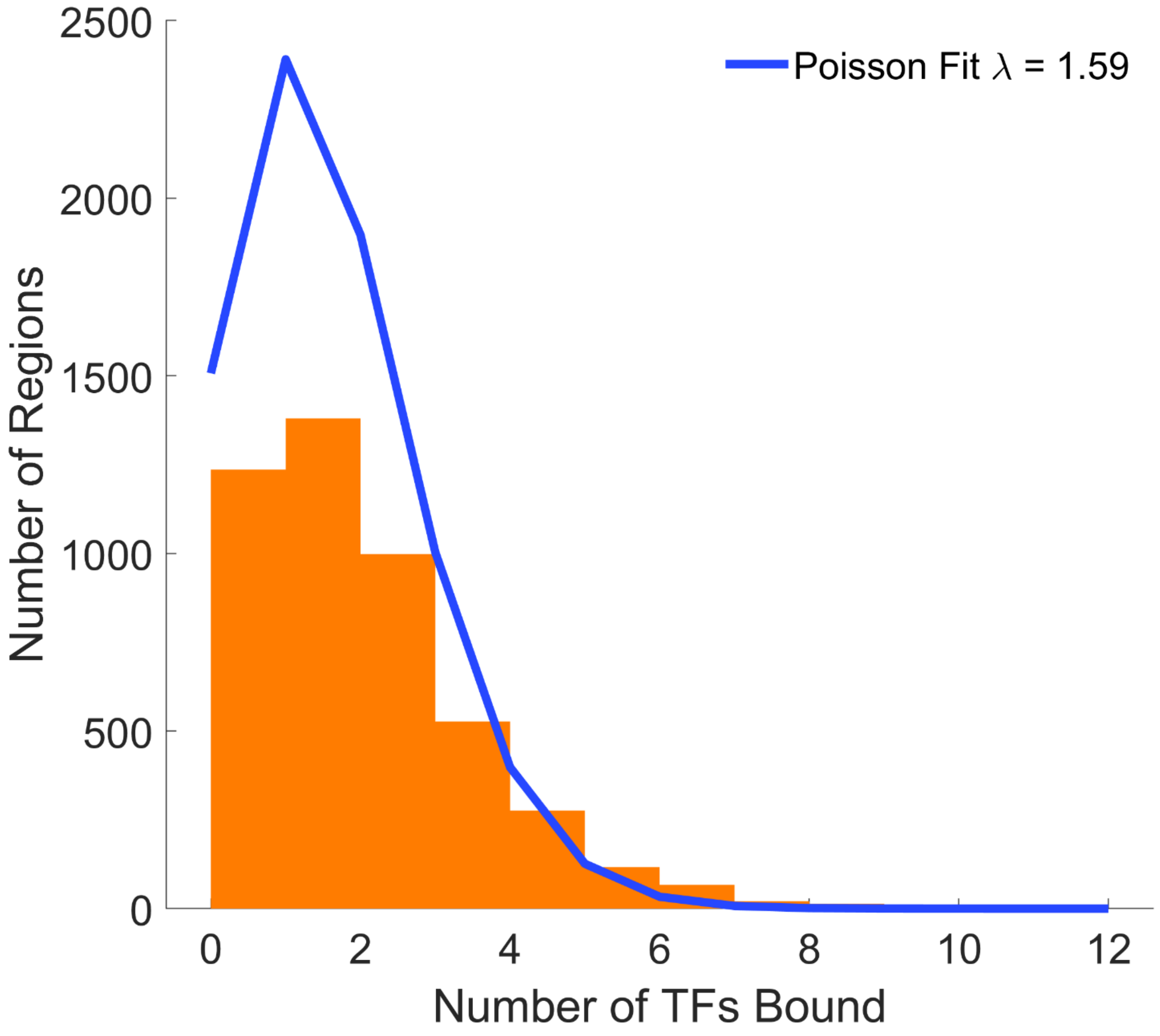

**Figure S14: Distribution of Number of TFs Binding All Non-Overlapping 1kbp Regions.** Most regions only have at most 2 TFs binding within them while a handful of regions have several TFs binding (as many as 11).

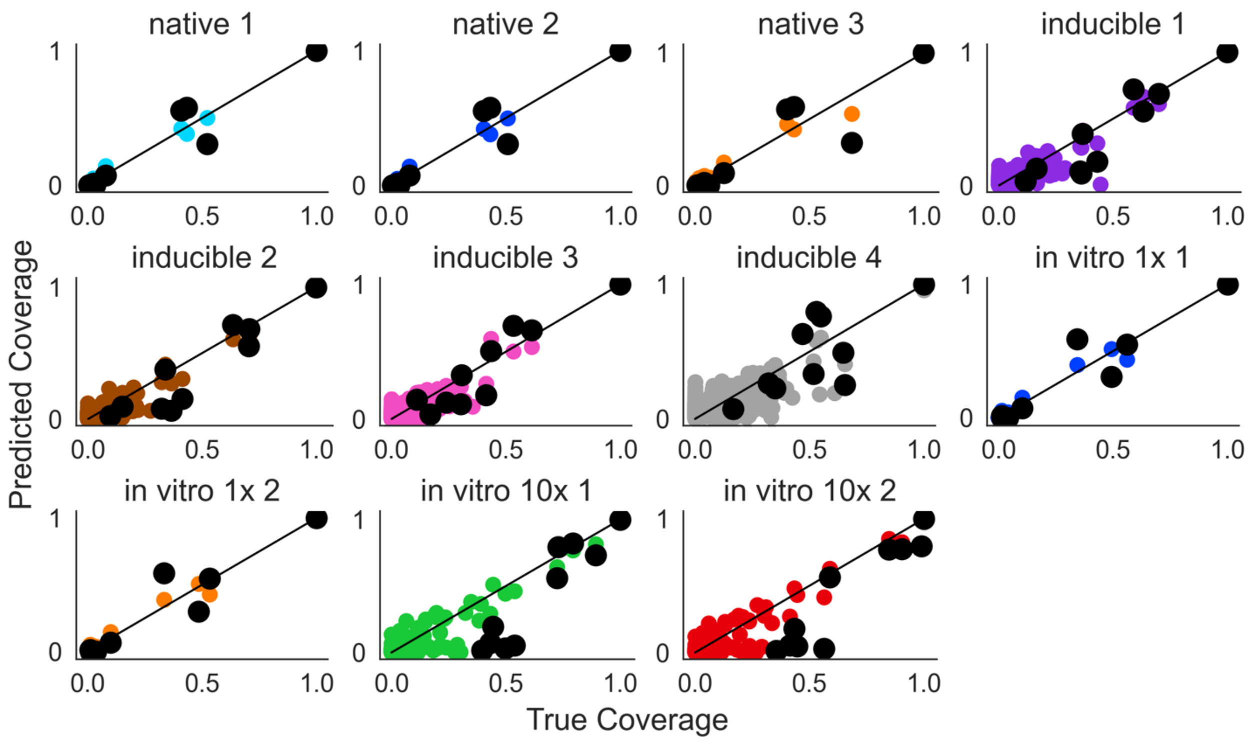

**Figure S15: Prediction of Normalized Coverage and Cross-Validation on Training Set for All Experiments for PdhR**. Predictions on a subset of representative experiments. Black circles show leave-one-out cross-validation predictions. All coverage is normalized between zero (baseline) and 1 (strongest called peak). Native=native promoter replicate number, inducible= inducible promoter replicate number, *in vitro* #x = *in vitro* ChIP-Seq replicate numbers at two different protein concentration

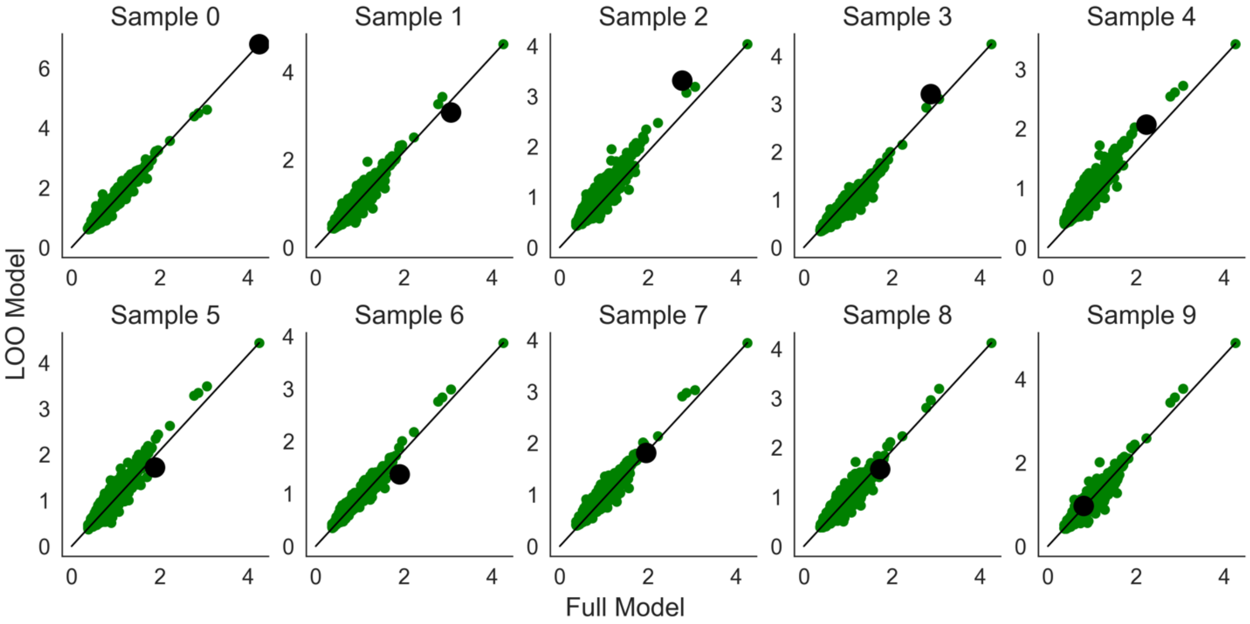

**Figure S16: Leave-One-Out Cross-Validation of Affinity Scores for PdhR.** For each of 10 samples starting with the strongest binding region, the sample was left out and an LOO model trained. For each LOO model, we plot the affinity score predictions on the full dataset against the predictions of the model built on the full dataset. The dark circle is the LOO sample.

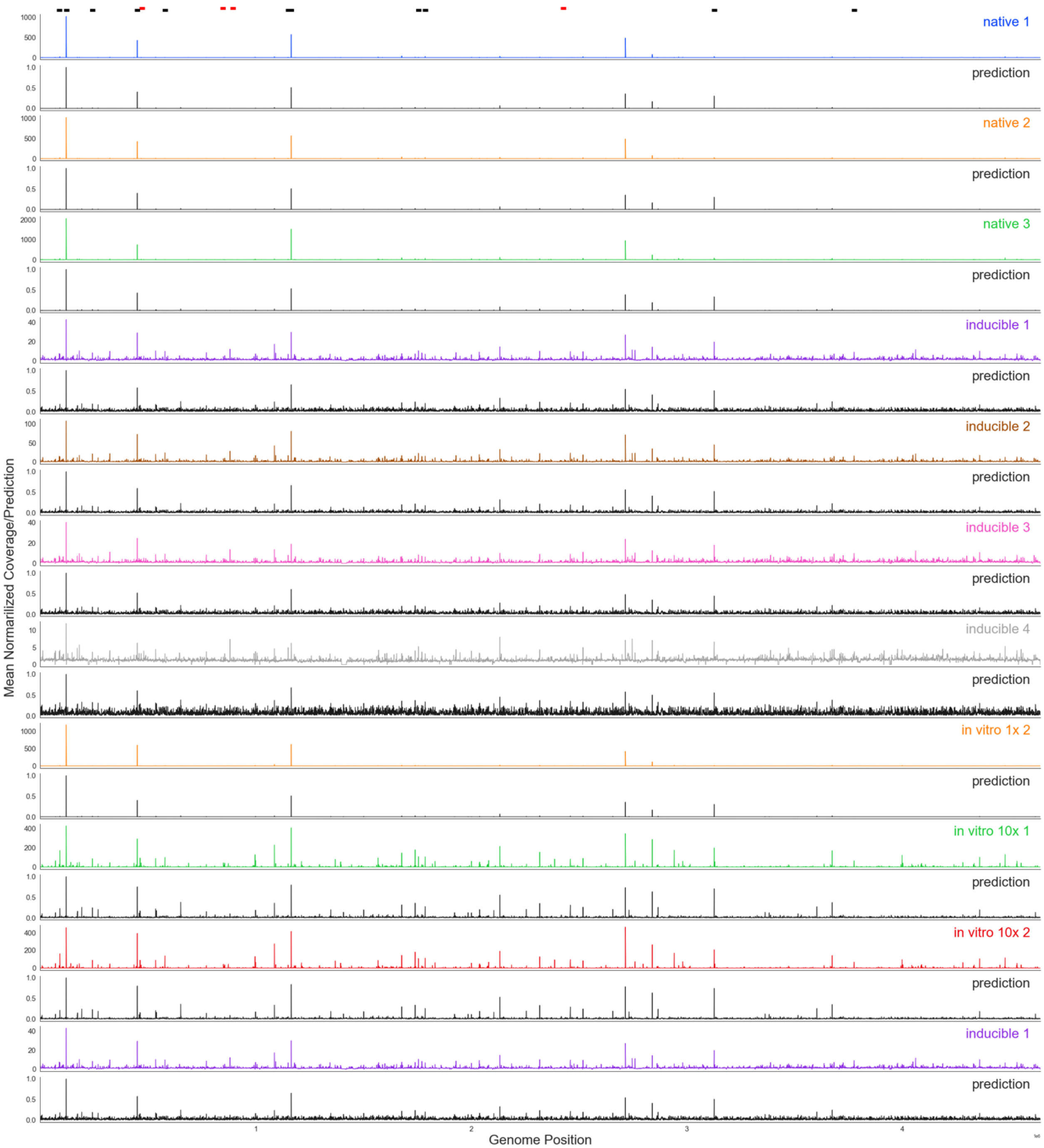

**Figure S17: Genome Wide Plot of Coverage and Coverage Predictions for All PdhR Experiments.** Each pair of tracks show actual coverage (blue) and model predicted coverage (black) for one experiment. Actual coverage in units of fold enrichment over mean coverage. Predicted coverage from zero to one. Positions of known sites (black rectangles) and Figure 2 details regions (red rectangles) shown above top track.

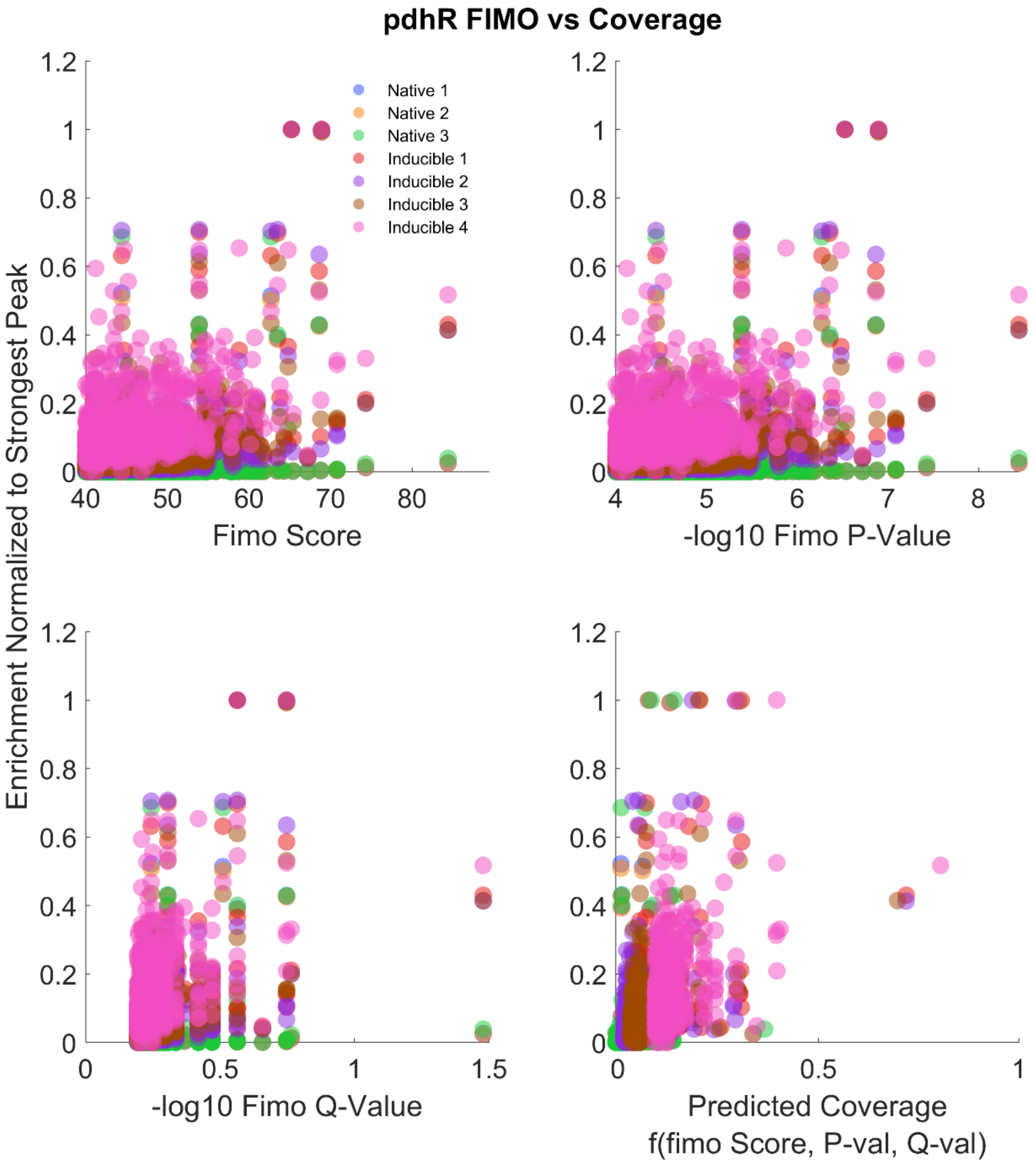

**Figure S18: PFMs are not Predictive of ChIP-Seq Coverage.** We used MEME to generate a PFM for pdhR using our collection of binding sites and scored all genomic sequences against this motif with FIMO (where higher scores indicate a better match to the PFM). The top-left scatterplot shows the FIMO score of each motif match on the x-axis and the ChIP-Seq enrichment of each FIMO hit normalized to the strongest pdhR binding site on the y-axis. The color of each point corresponds to a ChIP-Seq experiment. The top-right scatterplot shows the same enrichment values plotted against the negative log of the FIMO p-value and bottom-left uses the negative log of the FIMO q-value. For the bottom right plot, we perform a regression on the coverage values using the FIMO score, negative log p-value, and negative log q-value as predictors, and the x-axis shows the coverage predicted from this regression. In all cases, there is clearly no relationship between any scoring measure using the PFM and experimental coverage.

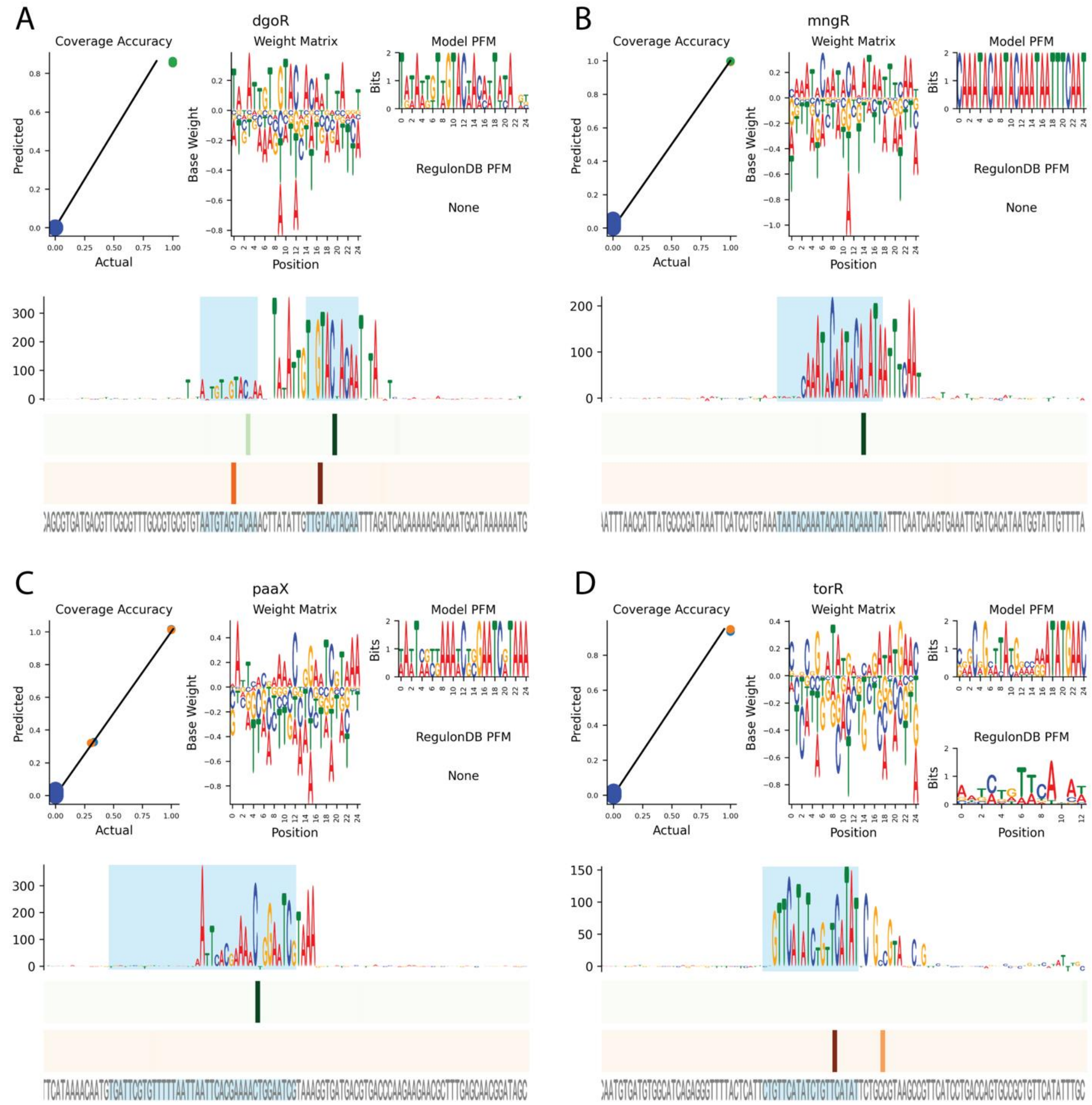

**Figure S19: Examples of Models Built for TFs with Only a Single Called Binding Region.** Top row of each panel displays a scatter plot of peak coverage prediction accuracy (left) and the weight matrix (right). The bottom half of each panel shows nucleotide level resolution predictions for the binding site for each TF. Sequence logo shows base contribution score. Cyan shading shows known binding site regions. Heatmaps shows predicted affinity at each position in positive (green) and negative (red) orientation. Sequence for binding site region show in gray.

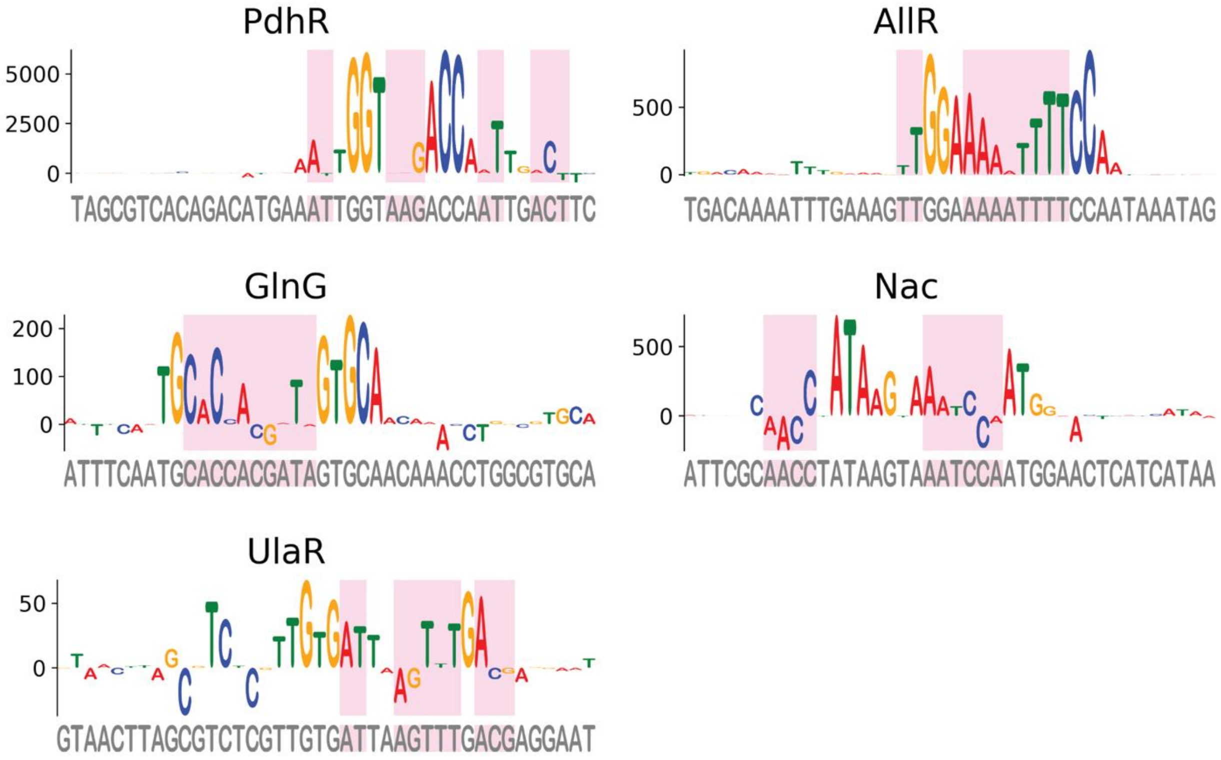

**Figure S20: Novel Site Design.** Novel binding sites were designed for 5 TFs. For each TF, the top singleton binding site was selected as a reference (sequence logo) and all combination of accessory bases (pink shading) we generated and used as input to the corresponding BoltzNet model.

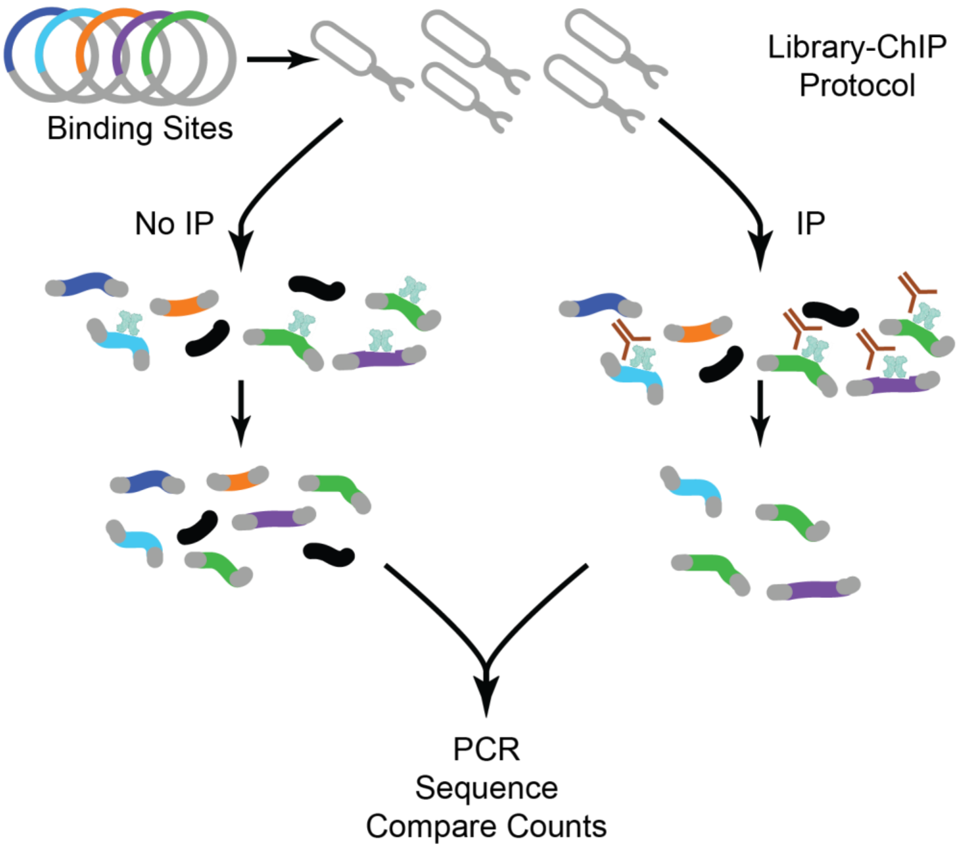

**Figure S21: Library-ChIP schematic.** A library of plasmids each containing 1 designed (or genomic reference) binding site is transformed into *E. coli* (top). After the crosslinking and DNA-shearing steps of ChIP, a small amount of DNA-containing lysate is reserved as a no-IP control (left). The remaining fraction of lysate undergoes IP as in a normal ChIP-Seq experiment (right). Control and IP samples then undergo two rounds of PCR to add Illumina adapters and barcodes, and NGS is performed. We then compare the abundance of each sequence in IP and control samples to determine the Library-ChIP enrichment.

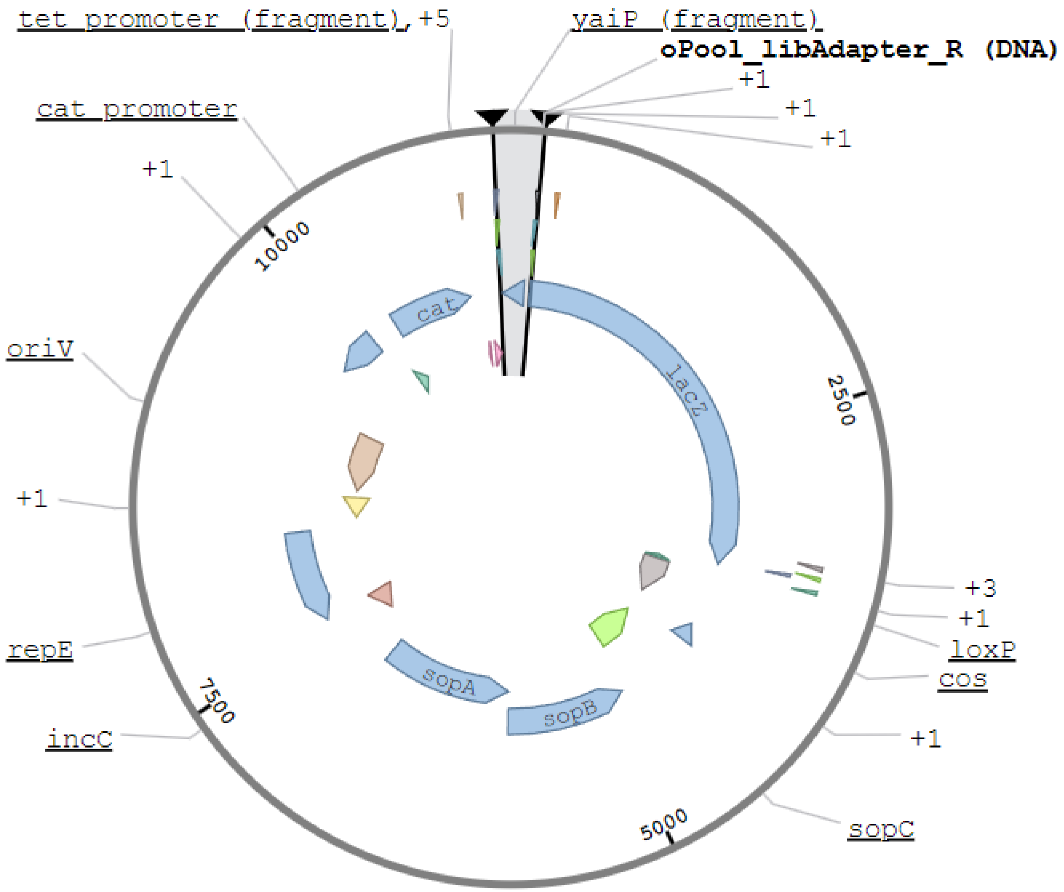

**Figure S22: pAMD Plasmid Map.** Library-ChIP pool contruction begins with inserting designed binding sites into the pAMD-BA plasmid. This vector is selected with chloramphenicol resistance (Cat gene), and contains two origins of replication – the standard oriV (single-copy), and ori2, which can make the plasmid high-copy under induction of a TrfA gene present in certain *E. coli* strains.

**Table S1: Table of TF ChIP-Seq Stats.**

| TF | Bnumber | Native SPA | Native FLAG | Native V5 | Inducible | New Regions | Found Known Regions | Total Known Regions | Total Regions | Has Autobinding |
|---|---|---|---|---|---|---|---|---|---|---|
| **aaeR** | b3243 | ✗ | ✗ | ✗ | ✓ | 69 | 0 | 0 | 69 | 1 |
| **agaR** | b3131 | ✓ | ✗ | ✗ | ✓ | 6 | 4 | 4 | 10 | 1 |
| **allR** | b0506 | ✓ | ✗ | ✓ | ✗ | 90 | 2 | 2 | 92 | 1 |
| **allS** | b0504 | ✗ | ✗ | ✗ | ✓ | 37 | 0 | 0 | 37 | 0 |
| **argP** | b2916 | ✗ | ✗ | ✗ | ✓ | 110 | 3 | 7 | 113 | 0 |
| **argR** | b3237 | ✗ | ✓ | ✗ | ✗ | 18 | 13 | 14 | 31 | 1 |
| **arsR** | b3501 | ✗ | ✗ | ✓ | ✗ | 64 | 1 | 1 | 65 | 1 |
| **ascG** | b2714 | ✗ | ✗ | ✓ | ✗ | 1 | 2 | 5 | 3 | 0 |
| **asnC** | b3743 | ✓ | ✗ | ✗ | ✗ | 7 | 0 | 0 | 7 | 1 |
| **basR** | b4113 | ✗ | ✗ | ✗ | ✓ | 1 | 7 | 9 | 8 | 1 |
| **betI** | b0313 | ✗ | ✗ | ✓ | ✓ | 17 | 1 | 1 | 18 | 1 |
| **cadC** | b4133 | ✗ | ✗ | ✗ | ✓ | 27 | 0 | 0 | 27 | 1 |
| **cbl** | b1987 | ✓ | ✗ | ✗ | ✓ | 17 | 1 | 2 | 18 | 0 |
| **cpxR** | b3912 | ✓ | ✗ | ✗ | ✗ | 8 | 15 | 39 | 23 | 1 |
| **csgD** | b1040 | ✗ | ✗ | ✗ | ✓ | 393 | 7 | 12 | 400 | 1 |
| **csiR** | b2664 | ✗ | ✗ | ✗ | ✓ | 86 | 1 | 1 | 87 | 0 |
| **cusR** | b0571 | ✗ | ✗ | ✗ | ✓ | 2 | 2 | 2 | 4 | 1 |
| **cynR** | b0338 | ✗ | ✗ | ✗ | ✓ | 104 | 1 | 1 | 105 | 1 |
| **cysB** | b1275 | ✗ | ✓ | ✗ | ✗ | 168 | 3 | 5 | 171 | 1 |
| **deoR** | b0840 | ✗ | ✗ | ✗ | ✓ | 35 | 3 | 5 | 38 | 0 |
| **dgoR** | b4479 | ✗ | ✗ | ✓ | ✓ | 0 | 1 | 1 | 1 | 1 |
| **dgsA** | b1594 | ✗ | ✗ | ✓ | ✗ | 3 | 3 | 3 | 6 | 1 |
| **dicA** | b1570 | ✗ | ✓ | ✗ | ✓ | 88 | 1 | 1 | 89 | 1 |
| **dicC** | b1569 | ✗ | ✗ | ✗ | ✓ | 279 | 0 | 0 | 279 | 1 |
| **dinJ** | b0226 | ✗ | ✗ | ✓ | ✓ | 0 | 1 | 1 | 1 | 1 |
| **ebgR** | b3075 | ✗ | ✗ | ✓ | ✓ | 2 | 0 | 0 | 2 | 1 |
| **envY** | b0566 | ✗ | ✗ | ✗ | ✓ | 3 | 0 | 0 | 3 | 0 |
| **exuR** | b3094 | ✗ | ✗ | ✓ | ✗ | 39 | 2 | 3 | 41 | 1 |
| **fabR** | b3963 | ✗ | ✗ | ✓ | ✗ | 1 | 2 | 2 | 3 | 0 |
| **fimZ** | b0535 | ✗ | ✗ | ✗ | ✓ | 49 | 1 | 1 | 50 | 0 |
| **fucR** | b2805 | ✗ | ✗ | ✗ | ✓ | 1 | 0 | 0 | 1 | 1 |
| **gadW** | b3515 | ✗ | ✗ | ✓ | ✓ | 6 | 1 | 4 | 7 | 1 |
| **gadX** | b3516 | ✗ | ✗ | ✗ | ✓ | 174 | 5 | 7 | 179 | 1 |
| **galR** | b2837 | ✗ | ✓ | ✗ | ✓ | 9 | 4 | 4 | 13 | 1 |
| **galS** | b2151 | ✗ | ✗ | ✓ | ✓ | 1 | 3 | 3 | 4 | 0 |
| **gcvA** | b2808 | ✗ | ✗ | ✓ | ✗ | 14 | 3 | 3 | 17 | 1 |
| **gcvR** | b2479 | ✗ | ✗ | ✓ | ✓ | 3 | 0 | 0 | 3 | 0 |
| **glnG** | b3868 | ✗ | ✗ | ✗ | ✓ | 166 | 3 | 4 | 169 | 1 |
| **glpR** | b3423 | ✗ | ✗ | ✓ | ✗ | 0 | 5 | 9 | 5 | 0 |
| **hcaR** | b2537 | ✗ | ✗ | ✗ | ✓ | 7 | 1 | 1 | 8 | 1 |
| **hdfR** | b4480 | ✗ | ✗ | ✗ | ✓ | 3 | 0 | 0 | 3 | 0 |
| **hipA** | b1507 | ✗ | ✗ | ✓ | ✓ | 19 | 1 | 2 | 20 | 1 |
| **hipB** | b1508 | ✗ | ✗ | ✗ | ✓ | 31 | 1 | 5 | 32 | 1 |
| **iclR** | b4018 | ✗ | ✗ | ✓ | ✗ | 2 | 2 | 2 | 4 | 1 |
| **iscR** | b2531 | ✗ | ✗ | ✓ | ✗ | 104 | 5 | 9 | 109 | 1 |

| TF | Bnumber | Native SPA | Native FLAG | Native V5 | Inducible | New Regions | Found Known Regions | Total Known Regions | Total Regions | Has Autobinding |
|---|---|---|---|---|---|---|---|---|---|---|
| **kdgR** | b1827 | ✗ | ✗ | ✓ | ✓ | 95 | 1 | 1 | 96 | 1 |
| **lacI** | b0345 | ✗ | ✗ | ✓ | ✓ | 0 | 3 | 3 | 3 | 0 |
| **lexA** | b4043 | ✗ | ✓ | ✗ | ✓ | 11 | 8 | 17 | 19 | 0 |
| **lldR** | b3604 | ✗ | ✗ | ✗ | ✓ | 21 | 2 | 2 | 23 | 1 |
| **marA** | b1531 | ✗ | ✓ | ✗ | ✓ | 44 | 6 | 19 | 50 | 1 |
| **matA** | b0294 | ✗ | ✗ | ✗ | ✓ | 6 | 0 | 0 | 6 | 1 |
| **mazE** | b2783 | ✓ | ✓ | ✗ | ✗ | 0 | 1 | 1 | 1 | 1 |
| **melR** | b4118 | ✗ | ✗ | ✗ | ✓ | 56 | 2 | 2 | 58 | 1 |
| **metR** | b3828 | ✓ | ✗ | ✗ | ✓ | 8 | 2 | 2 | 10 | 1 |
| **mhpR** | b0346 | ✗ | ✗ | ✓ | ✓ | 1 | 1 | 1 | 2 | 0 |
| **mlrA** | b2127 | ✗ | ✗ | ✓ | ✓ | 2 | 0 | 4 | 2 | 0 |
| **mngR** | b0730 | ✗ | ✗ | ✓ | ✓ | 0 | 1 | 1 | 1 | 1 |
| **modE** | b0761 | ✓ | ✗ | ✗ | ✓ | 4 | 4 | 7 | 8 | 0 |
| **mraZ** | b0081 | ✗ | ✗ | ✓ | ✓ | 96 | 3 | 3 | 99 | 1 |
| **nac** | b1988 | ✗ | ✗ | ✗ | ✓ | 477 | 2 | 2 | 479 | 1 |
| **nagC** | b0676 | ✓ | ✗ | ✗ | ✗ | 8 | 7 | 9 | 15 | 1 |
| **nanR** | b3226 | ✗ | ✗ | ✗ | ✓ | 386 | 2 | 2 | 388 | 1 |
| **nhaR** | b0020 | ✗ | ✗ | ✗ | ✓ | 0 | 2 | 4 | 2 | 1 |
| **nrdR** | b0413 | ✗ | ✗ | ✓ | ✗ | 0 | 3 | 3 | 3 | 0 |
| **nsrR** | b4178 | ✓ | ✗ | ✗ | ✗ | 4 | 2 | 4 | 6 | 0 |
| **ompR** | b3405 | ✗ | ✓ | ✗ | ✓ | 12 | 6 | 10 | 18 | 1 |
| **paaX** | b1399 | ✗ | ✗ | ✓ | ✓ | 0 | 2 | 3 | 2 | 0 |
| **pdhR** | b0113 | ✓ | ✗ | ✗ | ✓ | 193 | 11 | 11 | 204 | 1 |
| **pepA** | b4260 | ✗ | ✗ | ✓ | ✓ | 0 | 2 | 3 | 2 | 0 |
| **pspF** | b1303 | ✓ | ✗ | ✗ | ✓ | 0 | 2 | 2 | 2 | 1 |
| **purR** | b1658 | ✗ | ✗ | ✓ | ✗ | 14 | 10 | 13 | 24 | 1 |
| **racR** | b1356 | ✗ | ✗ | ✗ | ✓ | 1 | 1 | 1 | 2 | 1 |
| **relE** | b1563 | ✗ | ✗ | ✗ | ✓ | 8 | 0 | 0 | 8 | 0 |
| **rstA** | b1608 | ✗ | ✗ | ✗ | ✓ | 0 | 2 | 2 | 2 | 0 |
| **rutR** | b1013 | ✗ | ✗ | ✗ | ✓ | 150 | 4 | 5 | 154 | 1 |
| **sfsB** | b3188 | ✗ | ✗ | ✗ | ✓ | 258 | 0 | 0 | 258 | 1 |
| **sgcR** | b4300 | ✗ | ✗ | ✗ | ✓ | 1 | 0 | 0 | 1 | 1 |
| **sgrR** | b0069 | ✗ | ✗ | ✗ | ✓ | 1 | 2 | 2 | 3 | 1 |
| **srlR** | b2707 | ✗ | ✗ | ✗ | ✓ | 2 | 0 | 0 | 2 | 1 |
| **stpA** | b2669 | ✗ | ✗ | ✓ | ✗ | 201 | 0 | 1 | 201 | 1 |
| **sxy** | b0959 | ✗ | ✗ | ✗ | ✓ | 164 | 0 | 0 | 164 | 1 |
| **torR** | b0995 | ✗ | ✗ | ✓ | ✗ | 0 | 1 | 3 | 1 | 1 |
| **treR** | b4241 | ✗ | ✗ | ✗ | ✓ | 47 | 1 | 1 | 48 | 1 |
| **trpR** | b4393 | ✓ | ✗ | ✗ | ✗ | 1 | 5 | 5 | 6 | 1 |
| **ttk** | b3641 | ✗ | ✓ | ✓ | ✗ | 32 | 0 | 0 | 32 | 0 |
| **tyrR** | b1323 | ✓ | ✗ | ✗ | ✗ | 1 | 6 | 9 | 7 | 0 |
| **uidR** | b1618 | ✗ | ✗ | ✓ | ✗ | 1 | 0 | 0 | 1 | 1 |
| **ulaR** | b4191 | ✓ | ✗ | ✗ | ✓ | 71 | 2 | 2 | 73 | 1 |
| **uvrY** | b1914 | ✓ | ✗ | ✗ | ✗ | 0 | 1 | 1 | 1 | 0 |
| **uxuR** | b4324 | ✗ | ✗ | ✗ | ✓ | 60 | 1 | 3 | 61 | 1 |
| **yafC** | b0208 | ✗ | ✗ | ✗ | ✓ | 37 | 0 | 0 | 37 | 1 |
| **yafN** | b0232 | ✓ | ✗ | ✗ | ✓ | 225 | 0 | 0 | 225 | 1 |
| **yagI** | b0272 | ✗ | ✗ | ✗ | ✓ | 152 | 1 | 1 | 153 | 0 |

| TF | Bnumber | Native SPA | Native FLAG | Native V5 | Inducible | New Regions | Found Known Regions | Total Known Regions | Total Regions | Has Autobinding |
|---|---|---|---|---|---|---|---|---|---|---|
| **ybaQ** | b0483 | ✗ | ✗ | ✓ | ✗ | 43 | 0 | 0 | 43 | 0 |
| **ybcM** | b0546 | ✗ | ✗ | ✓ | ✗ | 16 | 0 | 0 | 16 | 1 |
| **ybhD** | b0768 | ✗ | ✗ | ✗ | ✓ | 2 | 0 | 0 | 2 | 1 |
| **ybiH** | b0796 | ✗ | ✗ | ✗ | ✓ | 4 | 1 | 1 | 5 | 1 |
| **ycgE** | b1162 | ✓ | ✗ | ✗ | ✓ | 1 | 1 | 1 | 2 | 1 |
| **yciT** | b1284 | ✗ | ✗ | ✓ | ✓ | 87 | 0 | 0 | 87 | 0 |
| **ycjW** | b1320 | ✗ | ✗ | ✓ | ✓ | 119 | 1 | 1 | 120 | 0 |
| **ycjZ** | b1328 | ✗ | ✗ | ✓ | ✓ | 13 | 1 | 1 | 14 | 1 |
| **ydaS** | b1357 | ✗ | ✗ | ✗ | ✓ | 1 | 0 | 0 | 1 | 1 |
| **ydcN** | b1434 | ✓ | ✗ | ✗ | ✓ | 0 | 1 | 1 | 1 | 1 |
| **ydcR** | b1439 | ✗ | ✗ | ✓ | ✗ | 20 | 0 | 0 | 20 | 1 |
| **yddM** | b1477 | ✗ | ✗ | ✓ | ✗ | 20 | 0 | 0 | 20 | 1 |
| **ydfH** | b1540 | ✗ | ✗ | ✓ | ✓ | 11 | 1 | 1 | 12 | 0 |
| **ydjF** | b1770 | ✓ | ✗ | ✗ | ✓ | 2 | 0 | 0 | 2 | 1 |
| **yeaM** | b1790 | ✗ | ✗ | ✓ | ✓ | 0 | 2 | 2 | 2 | 1 |
| **yeaT** | b1799 | ✗ | ✗ | ✓ | ✗ | 2 | 0 | 0 | 2 | 1 |
| **yedW** | b1969 | ✗ | ✗ | ✗ | ✓ | 0 | 2 | 2 | 2 | 0 |
| **yegW** | b2101 | ✗ | ✓ | ✗ | ✓ | 167 | 0 | 0 | 167 | 1 |
| **yeiE** | b2157 | ✗ | ✗ | ✓ | ✓ | 155 | 0 | 0 | 155 | 1 |
| **yeiI** | b2160 | ✗ | ✗ | ✗ | ✓ | 34 | 0 | 0 | 34 | 0 |
| **yfeC** | b2398 | ✗ | ✗ | ✓ | ✗ | 4 | 0 | 0 | 4 | 0 |
| **yfeT** | b2427 | ✗ | ✗ | ✓ | ✓ | 27 | 2 | 2 | 29 | 0 |
| **yfhH** | b2561 | ✓ | ✗ | ✗ | ✓ | 2 | 0 | 0 | 2 | 1 |
| **yfiE** | b2577 | ✗ | ✗ | ✓ | ✓ | 1 | 0 | 0 | 1 | 1 |
| **ygbI** | b2735 | ✗ | ✗ | ✓ | ✓ | 2 | 0 | 0 | 2 | 1 |
| **ygeH** | b2852 | ✗ | ✗ | ✗ | ✓ | 1 | 0 | 0 | 1 | 0 |
| **ygiT** | b3021 | ✓ | ✗ | ✗ | ✗ | 1 | 2 | 6 | 3 | 1 |
| **yhaJ** | b3105 | ✓ | ✗ | ✗ | ✗ | 4 | 0 | 0 | 4 | 1 |
| **yheO** | b3346 | ✗ | ✗ | ✓ | ✓ | 5 | 0 | 0 | 5 | 0 |
| **yhfR** | b3375 | ✗ | ✗ | ✓ | ✓ | 91 | 1 | 1 | 92 | 1 |
| **yhfZ** | b3383 | ✗ | ✗ | ✗ | ✓ | 1 | 0 | 0 | 1 | 1 |
| **yhjC** | b3521 | ✗ | ✗ | ✗ | ✓ | 3 | 0 | 0 | 3 | 0 |
| **yidZ** | b3711 | ✗ | ✗ | ✗ | ✓ | 233 | 0 | 0 | 233 | 0 |
| **yihW** | b3884 | ✗ | ✗ | ✓ | ✓ | 2 | 1 | 1 | 3 | 1 |
| **yijO** | b3954 | ✓ | ✗ | ✗ | ✓ | 1 | 0 | 0 | 1 | 0 |
| **yjgJ** | b4251 | ✗ | ✗ | ✓ | ✗ | 1 | 0 | 0 | 1 | 1 |
| **yjiR** | b4340 | ✗ | ✗ | ✓ | ✓ | 142 | 0 | 0 | 142 | 1 |
| **yjjQ** | b4365 | ✗ | ✗ | ✗ | ✓ | 365 | 2 | 2 | 367 | 1 |
| **ykgD** | b0305 | ✗ | ✗ | ✓ | ✗ | 2 | 0 | 0 | 2 | 1 |
| **yncC** | b1450 | ✗ | ✗ | ✓ | ✓ | 189 | 0 | 1 | 189 | 0 |
| **yoeB** | b4539 | ✗ | ✗ | ✓ | ✓ | 12 | 0 | 0 | 12 | 1 |
| **ypdC** | b2382 | ✗ | ✗ | ✗ | ✓ | 160 | 0 | 0 | 160 | 0 |
| **yqjI** | b3071 | ✓ | ✗ | ✗ | ✗ | 0 | 2 | 2 | 2 | 1 |
| **ytfH** | b4212 | ✗ | ✗ | ✓ | ✓ | 43 | 0 | 0 | 43 | 1 |
| **zraR** | b4004 | ✗ | ✗ | ✗ | ✓ | 0 | 2 | 2 | 2 | 1 |
| **zur** | b4046 | ✗ | ✗ | ✓ | ✗ | 4 | 2 | 2 | 6 | 0 |

**Table S2: Table of Model Weight Matrices and Corresponding PFMs.**

| TF | Bnumber | Model PWM | Model PFM |
|---|---|---|---|
| **aaeR** | b3243 | 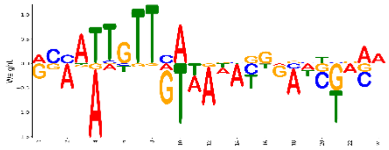 | 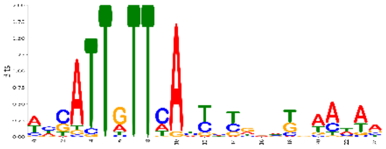 |
| **agaR** | b3131 | 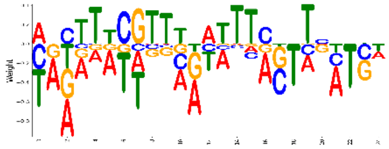 | 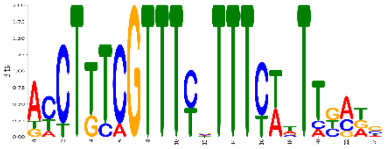 |
| **allR** | b0506 | 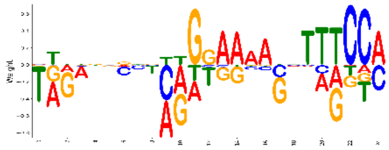 | 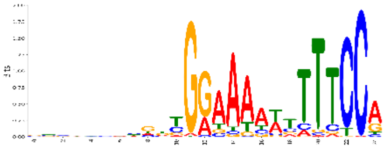 |
| **allS** | b0504 | None | None |
| **argP** | b2916 | 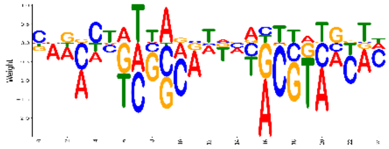 | 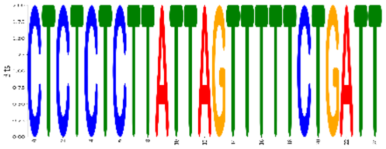 |
| **argR** | b3237 | 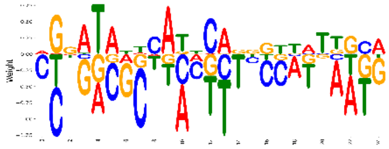 | 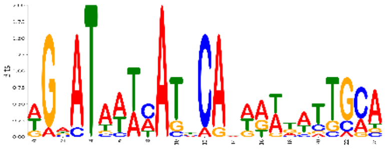 |
| **arsR** | b3501 | 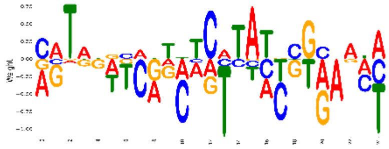 | 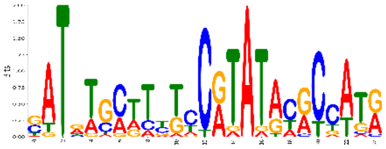 |
| **ascG** | b2714 | 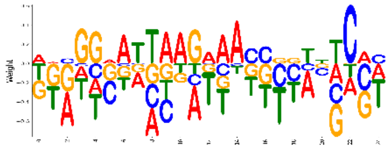 | 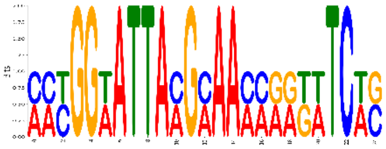 |
| **asnC** | b3743 | 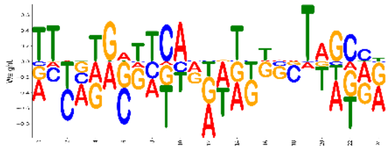 | 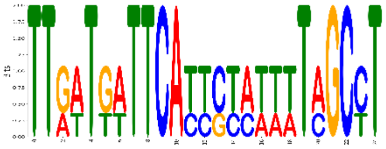 |
| **basR** | b4113 | 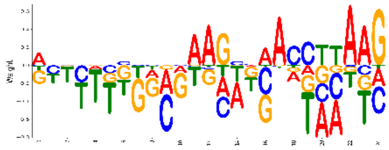 | 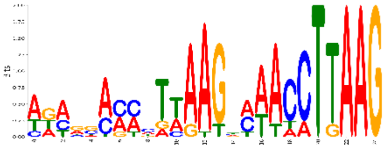 |
| **betI** | b0313 | 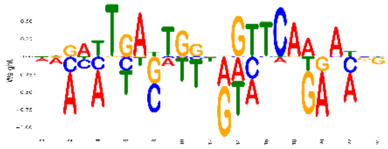 | 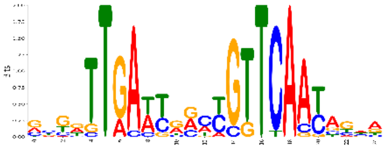 |

| TF | Bnumber | Model PWM | Model PFM |
| --- | --- | --- | --- |
| **cadC** | b4133 | 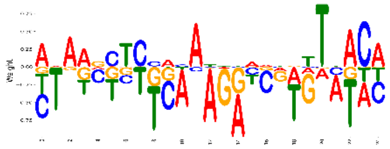 | 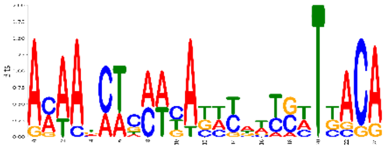 |
| **cbl** | b1987 | 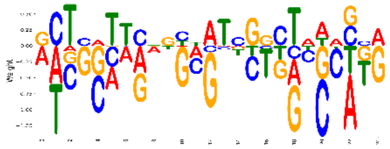 | 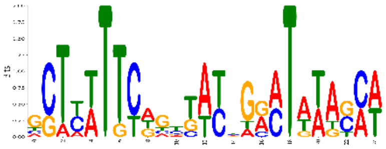 |
| **cpxR** | b3912 | 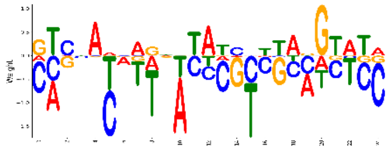 | 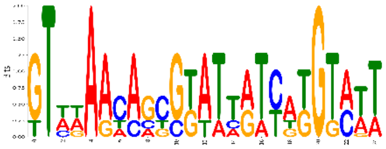 |
| **csgD** | b1040 | None | None |
| **csiR** | b2664 | None | None |
| **cusR** | b0571 | 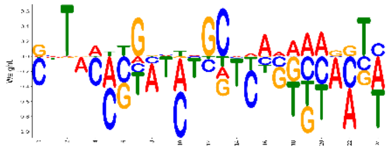 | 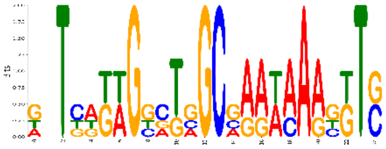 |
| **cynR** | b0338 | 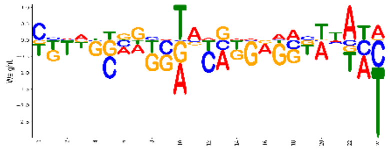 | 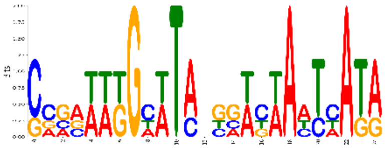 |
| **cysB** | b1275 | 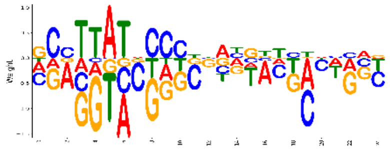 | 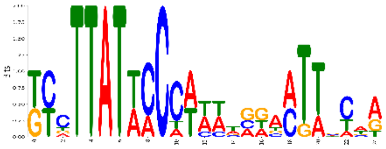 |
| **deoR** | b0840 | 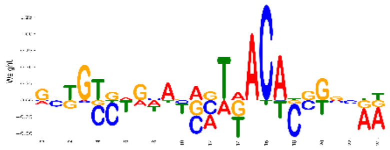 | 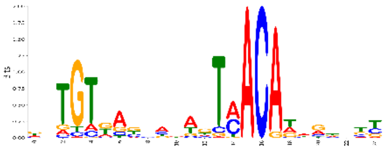 |
| **dgoR** | b4479 | 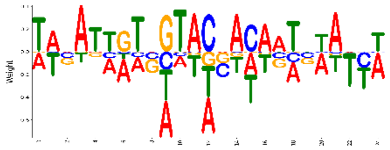 | 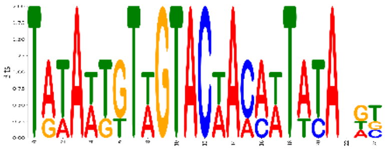 |
| **dgsA** | b1594 | 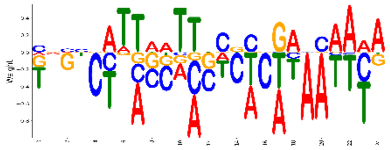 | 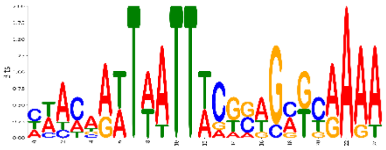 |
| **dicA** | b1570 | 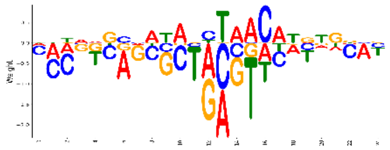 | 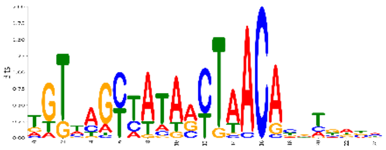 |

| TF | Bnumber | Model PWM | Model PFM |
|---|---|---|---|
| **dicC** | b1569 | 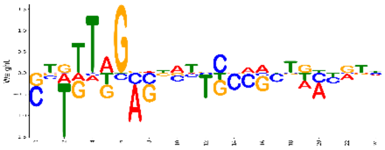 | 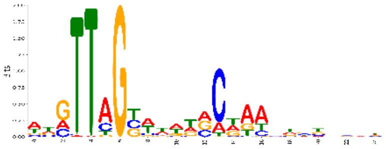 |
| **dinJ** | b0226 | 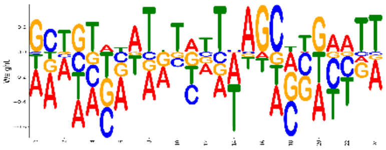 | 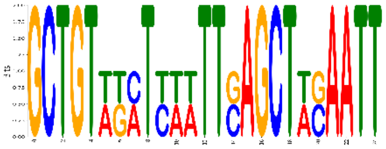 |
| **ebgR** | b3075 | 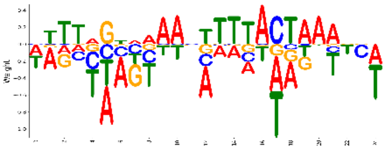 | 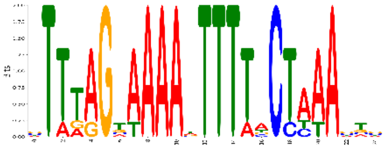 |
| **envY** | b0566 | 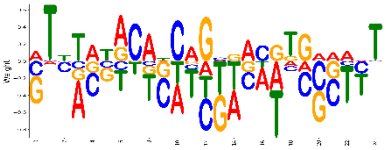 | 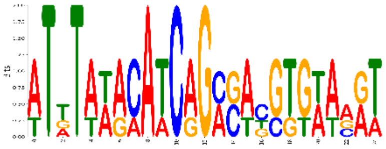 |
| **exuR** | b3094 | 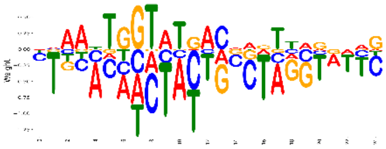 | 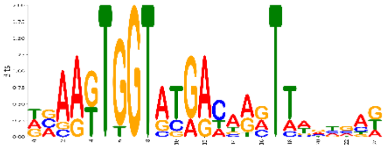 |
| **fabR** | b3963 | 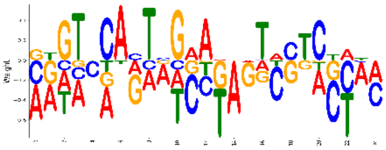 | 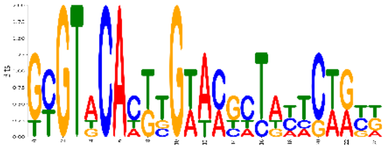 |
| **fimZ** | b0535 | 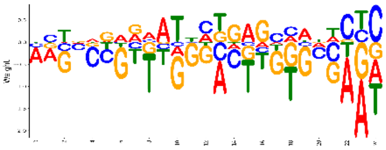 | 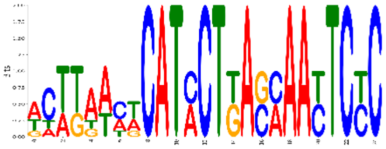 |
| **fucR** | b2805 | 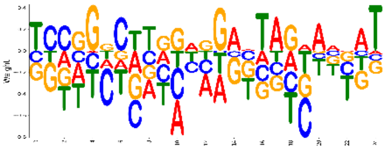 | 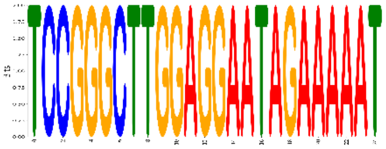 |
| **gadW** | b3515 | 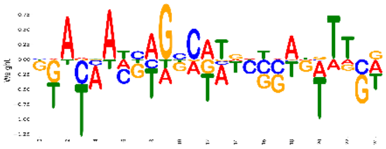 | 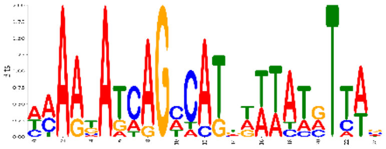 |
| **gadX** | b3516 | None | None |
| **galR** | b2837 | 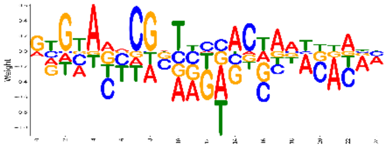 | 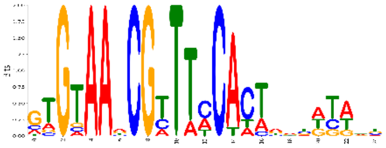 |

| TF | Bnumber | Model PWM | Model PFM |
|---|---|---|---|
| **galS** | b2151 | 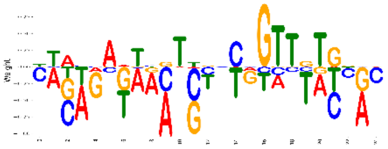 | 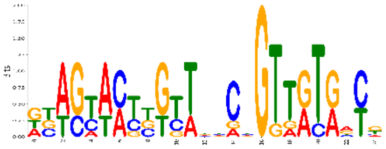 |
| **gcvA** | b2808 | 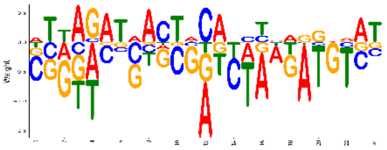 | 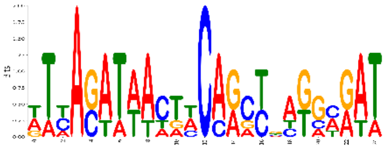 |
| **gcvR** | b2479 | None | None |
| **glnG** | b3868 | 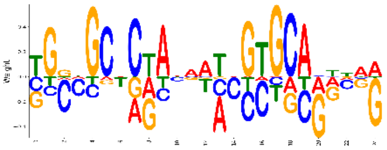 | 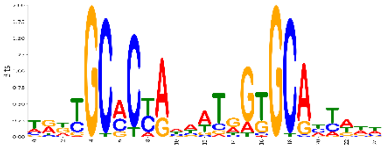 |
| **glpR** | b3423 | 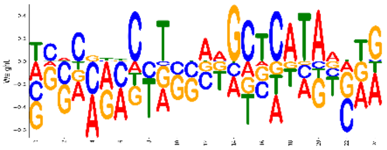 | 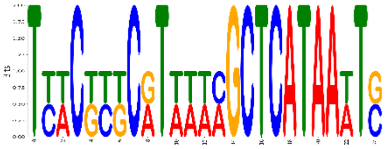 |
| **hcaR** | b2537 | 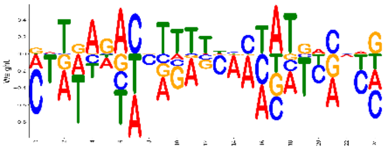 | 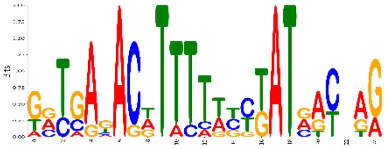 |
| **hdfR** | b4480 | 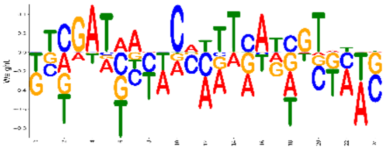 | 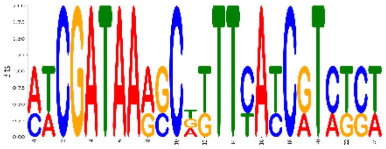 |
| **hipA** | b1507 | None | None |
| **hipB** | b1508 | None | None |
| **iclR** | b4018 | 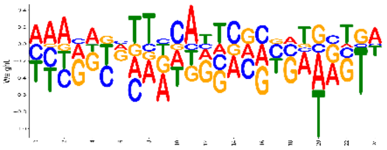 | 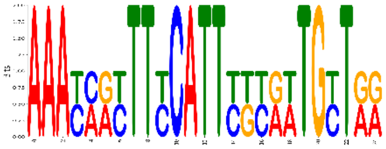 |
| **iscR** | b2531 | 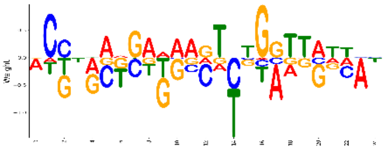 | 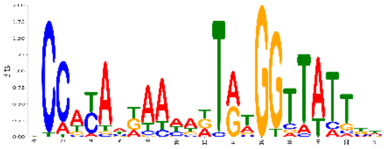 |
| **kdgR** | b1827 | 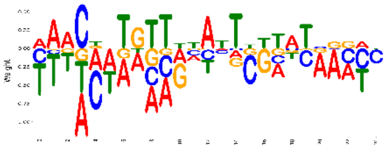 | 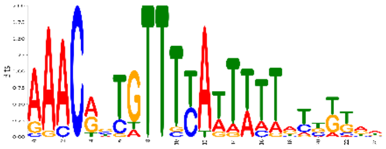 |

| TF | Bnumber | Model PWM | Model PFM |
|---|---|---|---|
| **lacI** | b0345 | 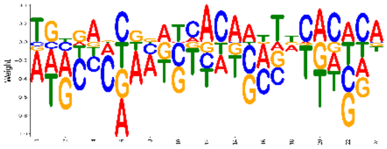 | 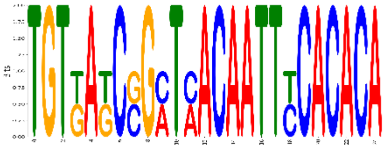 |
| **lexA** | b4043 | 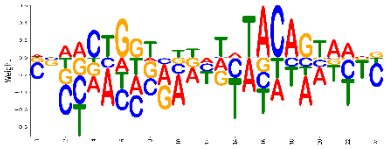 | 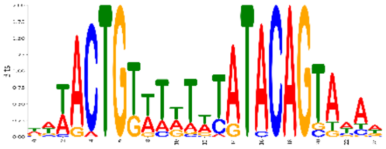 |
| **lldR** | b3604 | 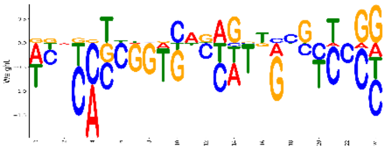 | 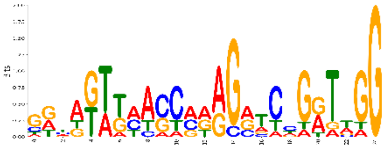 |
| **marA** | b1531 | 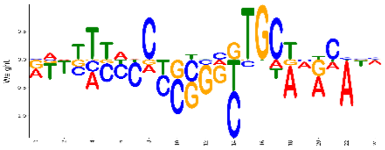 | 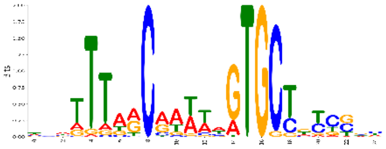 |
| **matA** | b0294 | 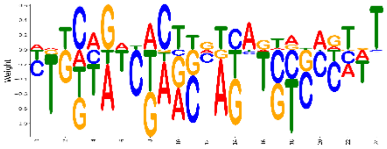 | 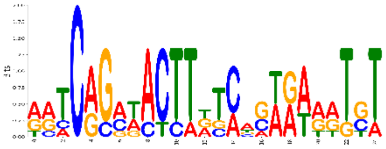 |
| **mazE** | b2783 | None | None |
| **melR** | b4118 | 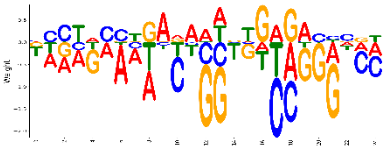 | 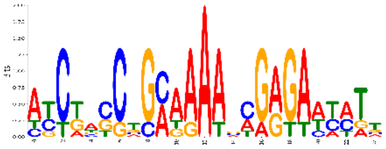 |
| **metR** | b3828 | 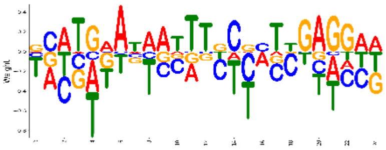 | 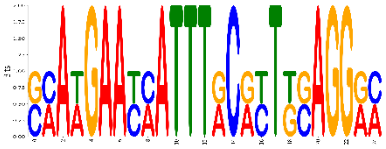 |
| **mhpR** | b0346 | 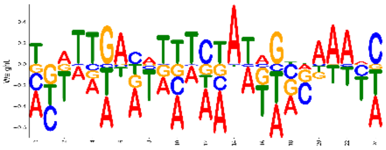 | 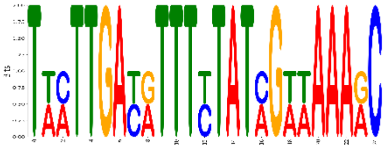 |
| **mlrA** | b2127 | 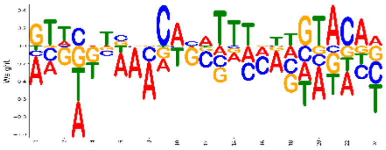 | 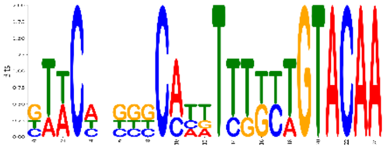 |
| **mngR** | b0730 | 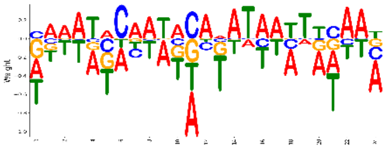 | 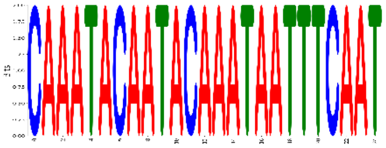 |

| TF | Bnumber | Model PWM | Model PFM |
|---|---|---|---|
| **modE** | b0761 | 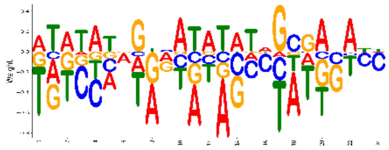 | 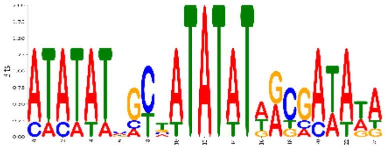 |
| **mraZ** | b0081 | 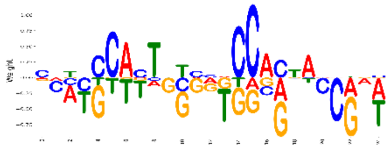 | 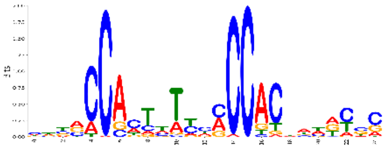 |
| **nac** | b1988 | 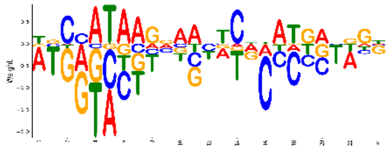 | 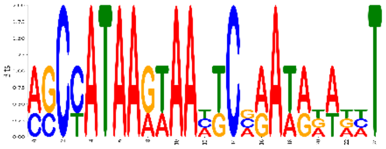 |
| **nagC** | b0676 | 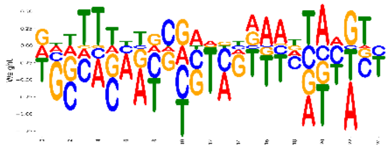 | 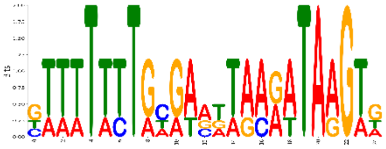 |
| **nanR** | b3226 | None | None |
| **nhaR** | b0020 | 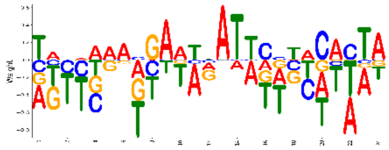 | 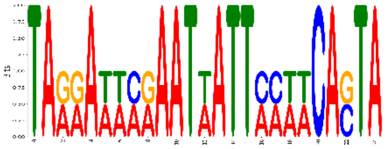 |
| **nrdR** | b0413 | 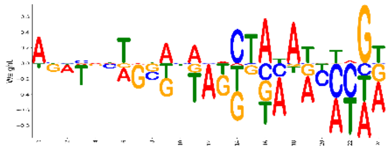 | 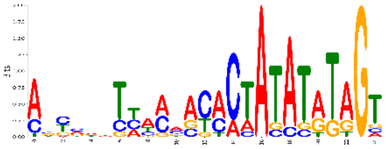 |
| **nsrR** | b4178 | 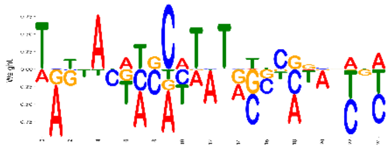 | 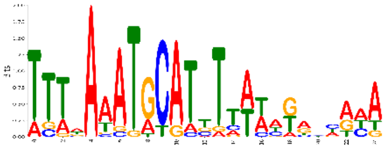 |
| **ompR** | b3405 | 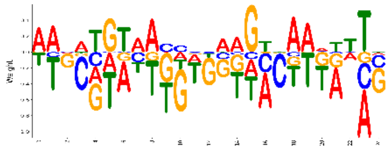 | 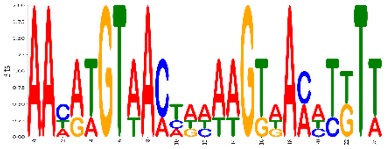 |
| **paaX** | b1399 | 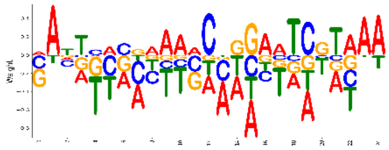 | 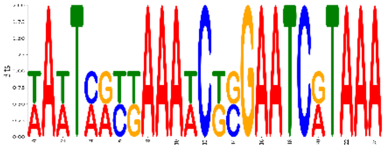 |
| **pdhR** | b0113 | 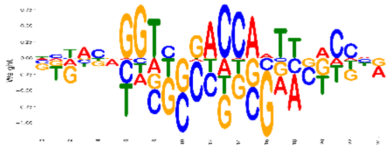 | 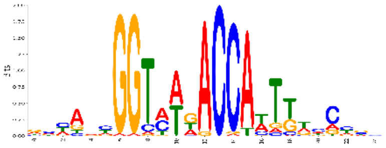 |

| TF | Bnumber | Model PWM | Model PFM |
|---|---|---|---|
| **pepA** | b4260 | 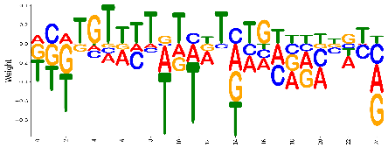 | 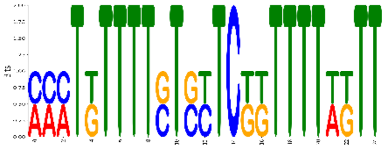 |
| **pspF** | b1303 | 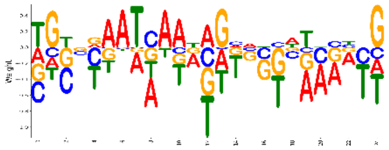 | 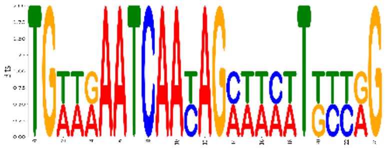 |
| **purR** | b1658 | 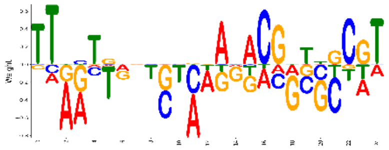 | 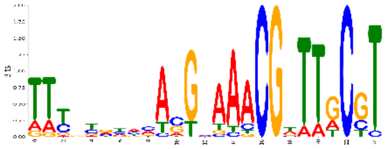 |
| **racR** | b1356 | 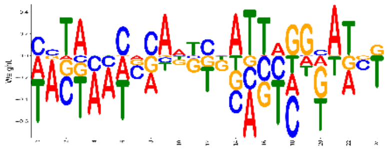 | 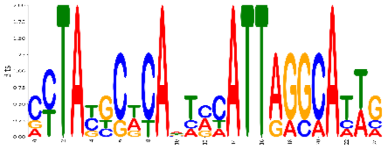 |
| **relE** | b1563 | 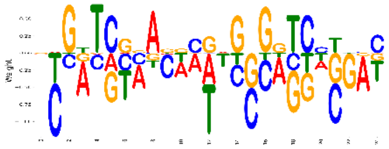 | 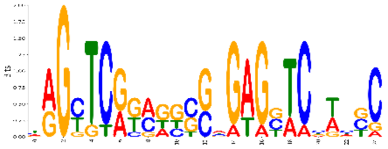 |
| **rstA** | b1608 | 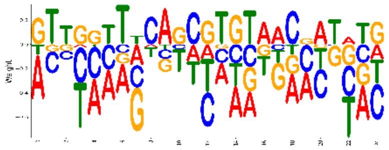 | 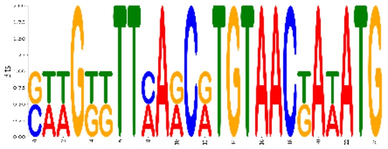 |
| **rutR** | b1013 | 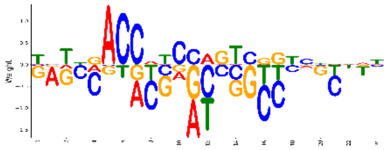 | 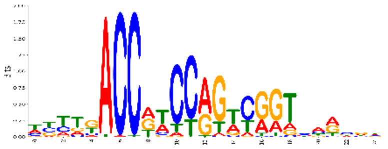 |
| **sfsB** | b3188 | 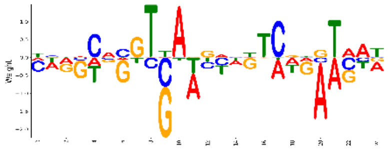 | 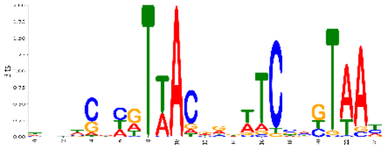 |
| **sgcR** | b4300 | 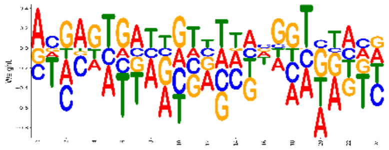 | 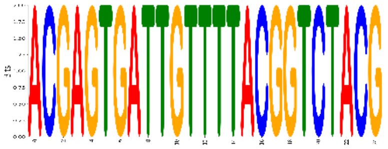 |
| **sgrR** | b0069 | None | None |
| **srlR** | b2707 | 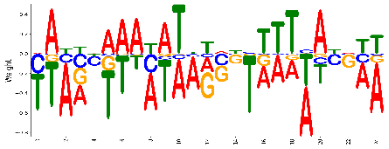 | 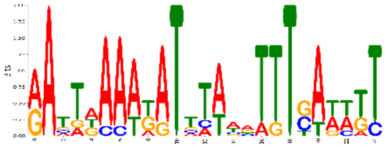 |
| **stpA** | b2669 | None | None |

| TF | Bnumber | Model PWM | Model PFM |
|---|---|---|---|
| **sxy** | b0959 | 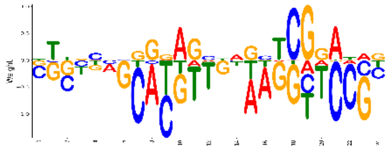 | 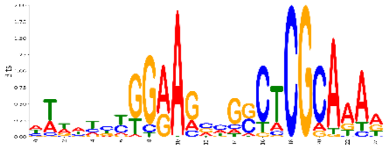 |
| **torR** | b0995 | 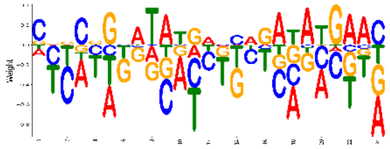 | 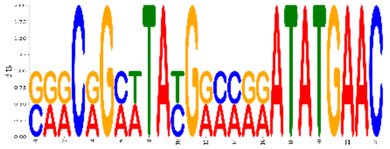 |
| **treR** | b4241 | 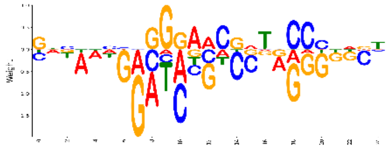 | 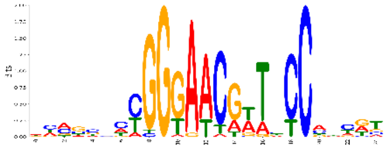 |
| **trpR** | b4393 | 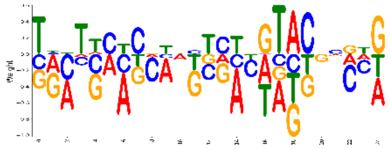 | 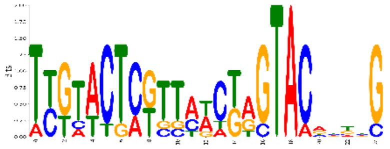 |
| **ttk** | b3641 | 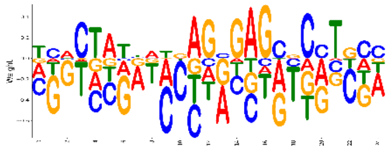 | 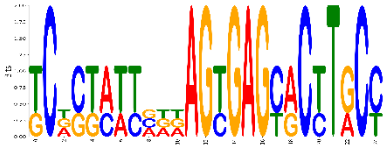 |
| **tyrR** | b1323 | 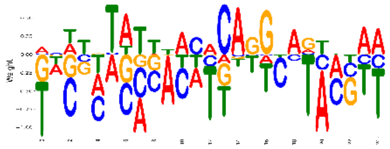 | 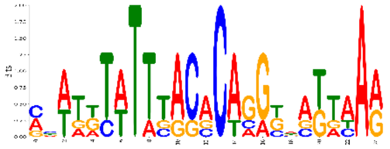 |
| **uidR** | b1618 | 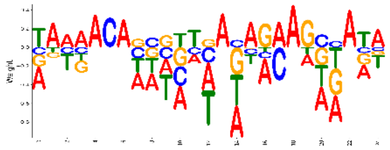 | 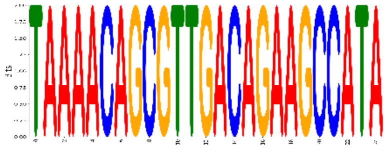 |
| **ulaR** | b4191 | 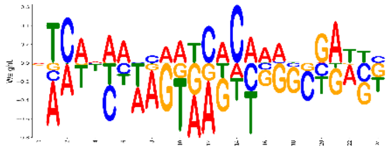 | 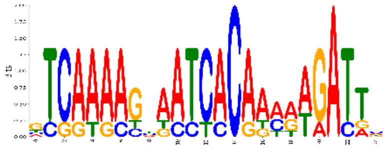 |
| **uvrY** | b1914 | 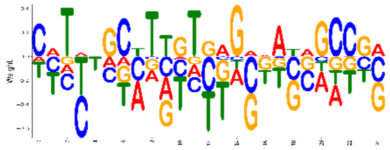 | 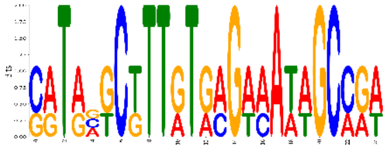 |
| **uxuR** | b4324 | 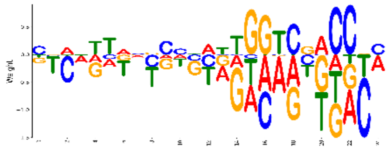 | 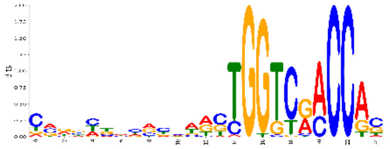 |
| **yafC** | b0208 | None | None |

| TF | Bnumber | Model PWM | Model PFM |
|---|---|---|---|
| **yafN** | b0232 | 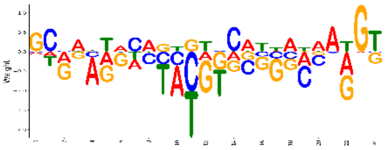 | 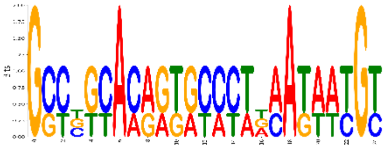 |
| **yagI** | b0272 | 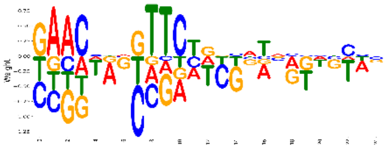 | 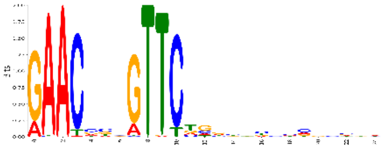 |
| **ybaQ** | b0483 | 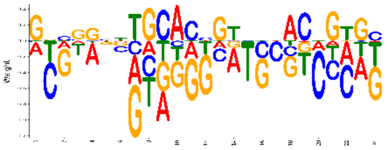 | 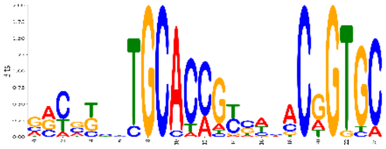 |
| **ybcM** | b0546 | 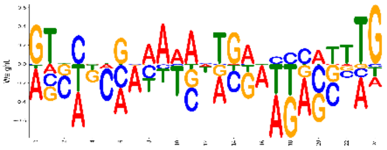 | 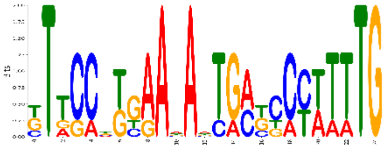 |
| **ybhD** | b0768 | 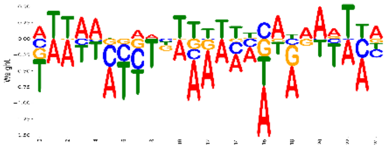 | 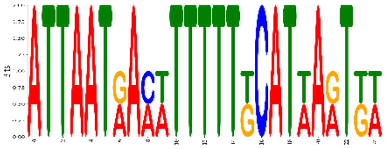 |
| **ybiH** | b0796 | 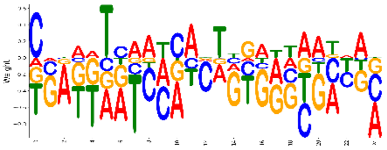 | 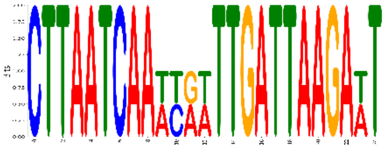 |
| **ycgE** | b1162 | 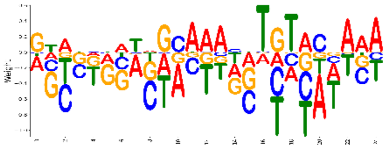 | 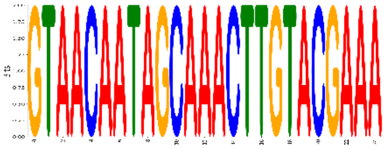 |
| **yciT** | b1284 | 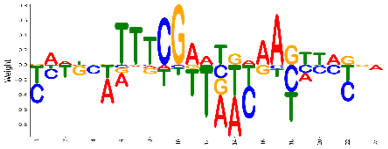 | 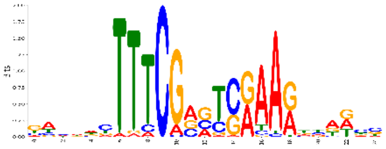 |
| **ycjW** | b1320 | 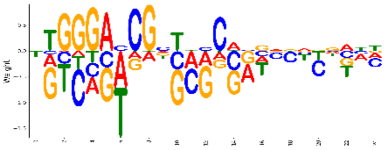 | 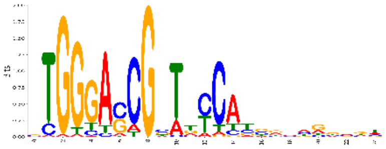 |
| **ycjZ** | b1328 | 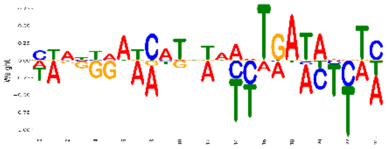 | 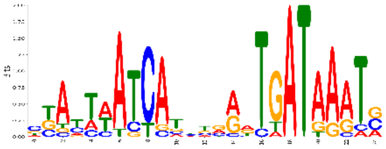 |

| TF | Bnumber | Model PWM | Model PFM |
|---|---|---|---|
| **ydaS** | b1357 | 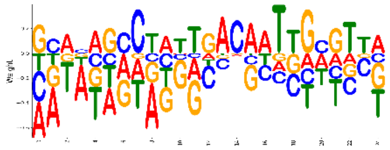 | 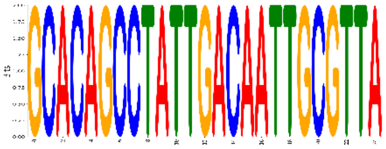 |
| **ydcN** | b1434 | 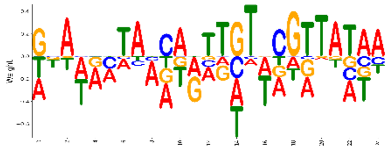 | 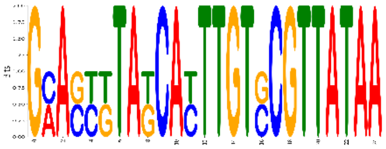 |
| **ydcR** | b1439 | 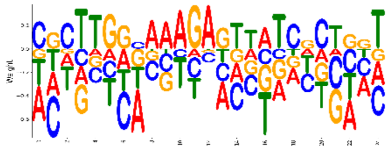 | 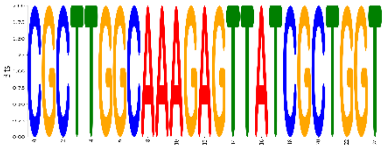 |
| **yddM** | b1477 | 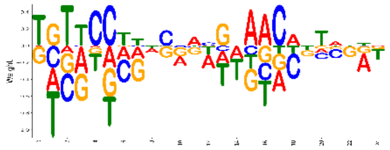 | 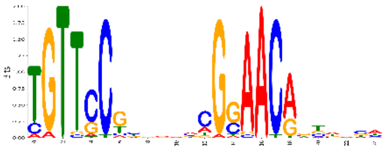 |
| **ydfH** | b1540 | 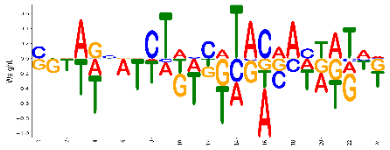 | 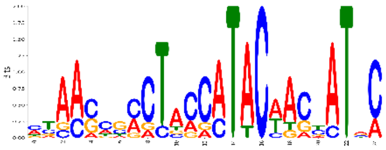 |
| **ydjF** | b1770 | 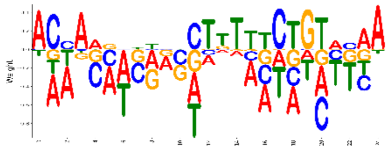 | 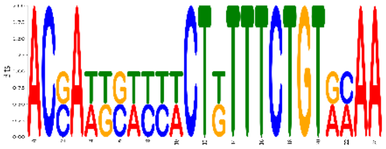 |
| **yeaM** | b1790 | 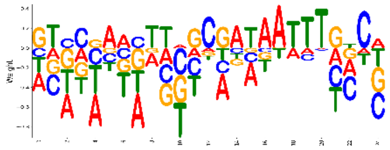 | 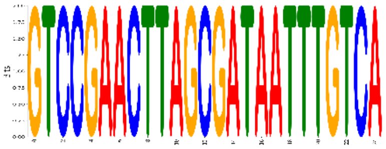 |
| **yeaT** | b1799 | 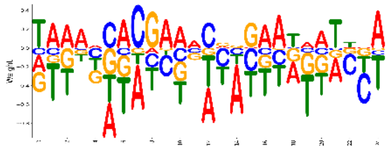 | 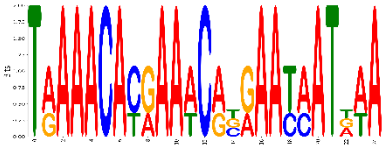 |
| **yedW** | b1969 | None | None |
| **yegW** | b2101 | 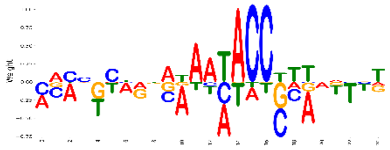 | 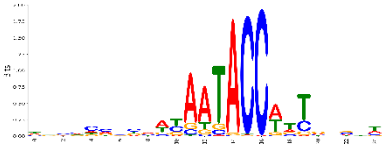 |
| **yeiE** | b2157 | 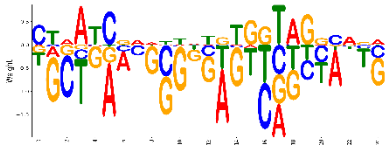 | 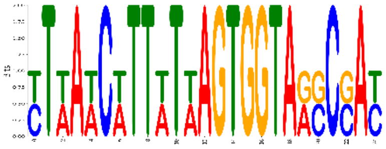 |

| TF | Bnumber | Model PWM | Model PFM |
|---|---|---|---|
| **yeiI** | b2160 | 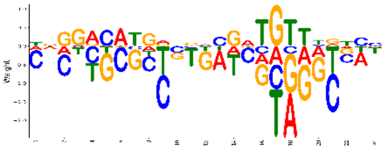 | 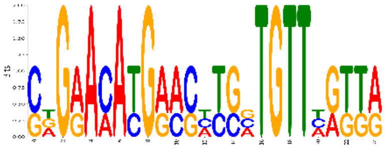 |
| **yfeC** | b2398 | 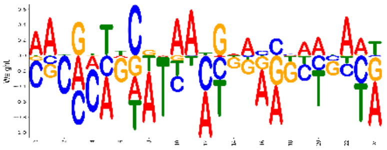 | 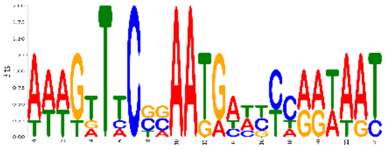 |
| **yfeT** | b2427 | 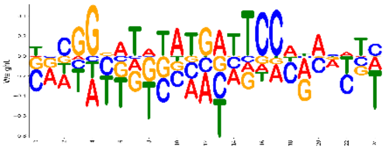 | 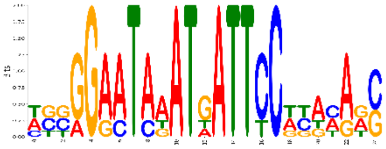 |
| **yfhH** | b2561 | 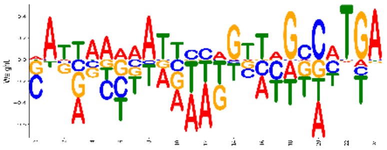 | 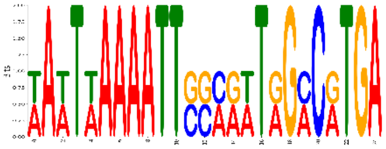 |
| **yfiE** | b2577 | None | None |
| **ygbI** | b2735 | 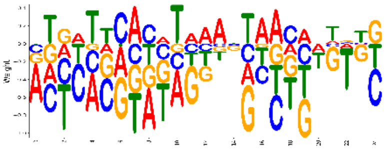 | 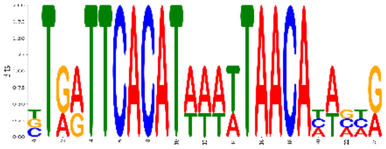 |
| **ygeH** | b2852 | 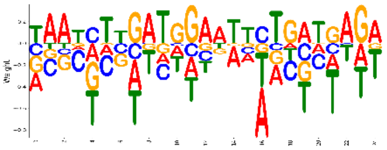 | 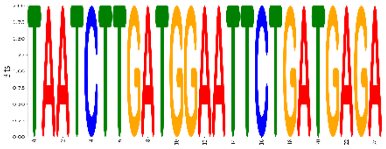 |
| **ygiT** | b3021 | 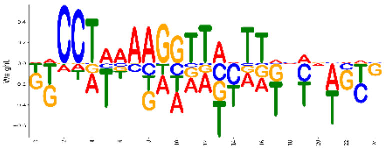 | 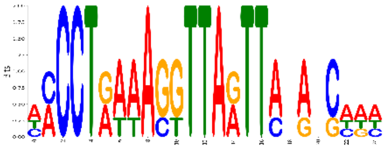 |
| **yhaJ** | b3105 | 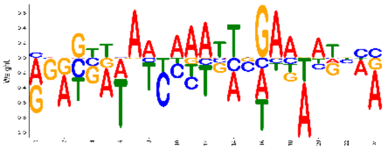 | 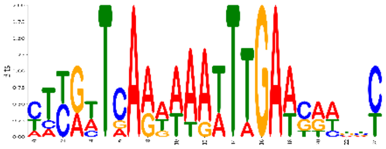 |
| **yheO** | b3346 | 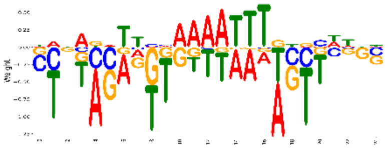 | 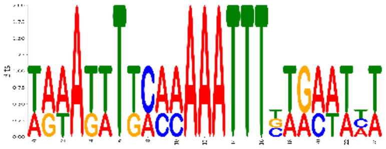 |
| **yhfR** | b3375 | 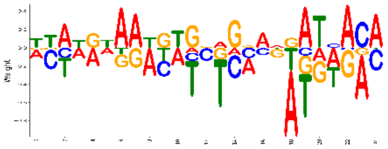 | 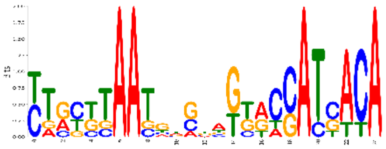 |

| TF | Bnumber | Model PWM | Model PFM |
| --- | --- | --- | --- |
| **yhfZ** | b3383 | 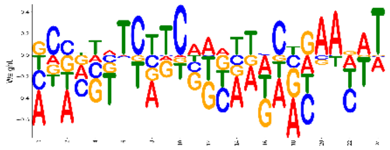 | 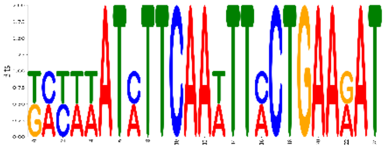 |
| **yhjC** | b3521 | 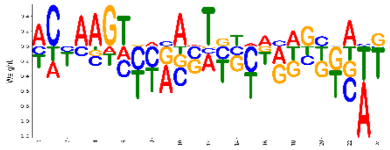 | 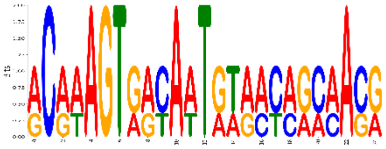 |
| **yidZ** | b3711 | 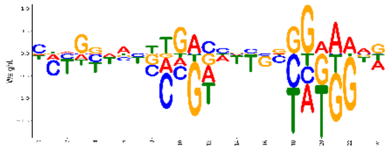 | 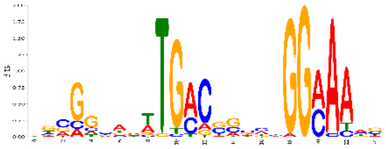 |
| **yihW** | b3884 | 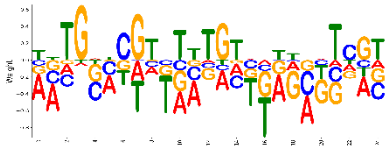 | 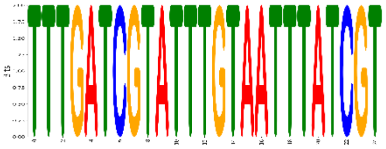 |
| **yijO** | b3954 | 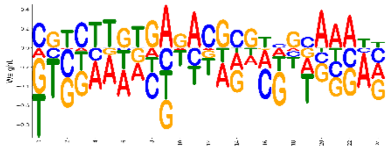 | 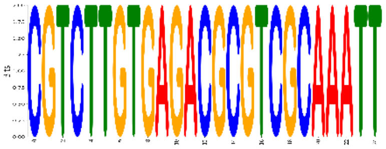 |
| **yjgJ** | b4251 | None | None |
| **yjiR** | b4340 | 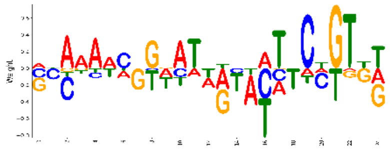 | 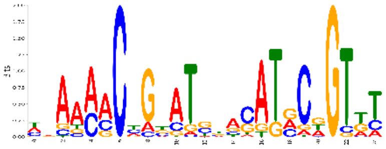 |
| **yjjQ** | b4365 | 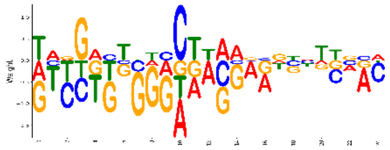 | 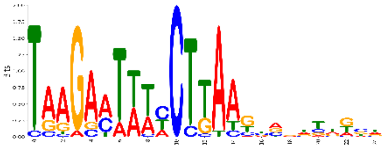 |
| **ykgD** | b0305 | 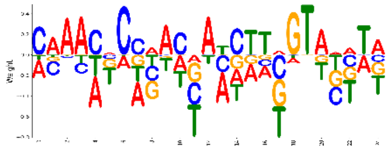 | 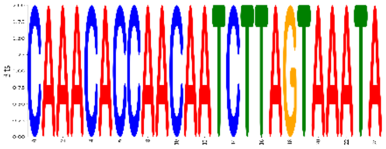 |
| **yncC** | b1450 | 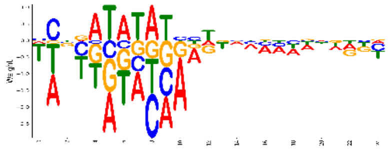 | 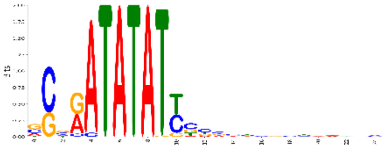 |
| **yoeB** | b4539 | 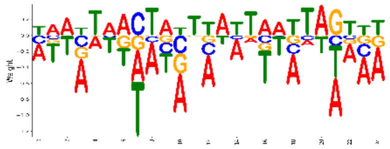 | 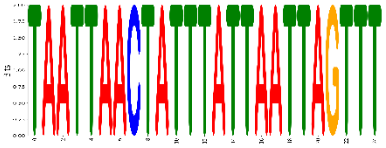 |

| TF | Bnumber | Model PWM | Model PFM |
|---|---|---|---|
| **ypdC** | b2382 | 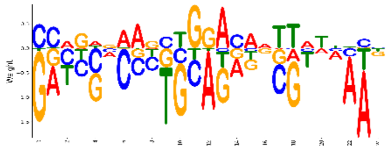 | 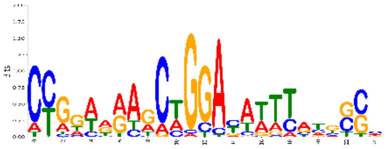 |
| **yqjI** | b3071 | 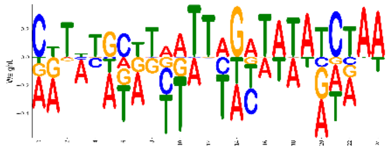 | 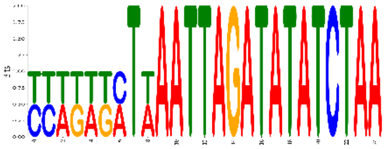 |
| **ytfH** | b4212 | 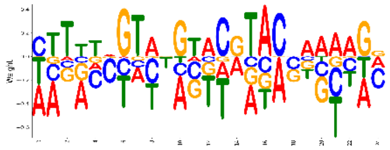 | 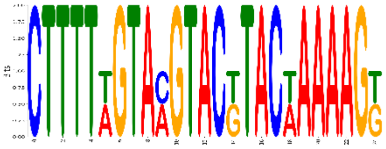 |
| **zraR** | b4004 | 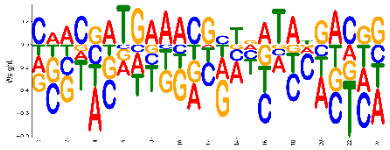 | 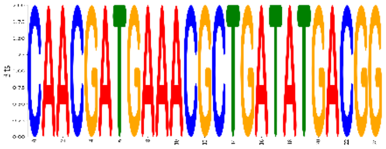 |
| **zur** | b4046 | 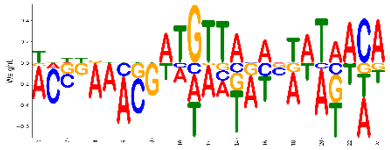 | 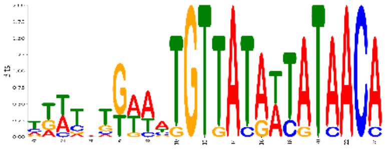 |

**Table S3: Table of ChIP-Seq Artifacts.**

| Start | Stop | Artifact Type |
|---|---|---|
| 1792 | 2084 | **NagC** |
| 83400 | 84400 | **Strong** |
| 228650 | 229000 | **Strong** |
| 262600 | 263600 | **Strong** |
| 412000 | 413000 | **Strong** |
| 451400 | 452400 | **Weak** |
| 656800 | 657800 | **Strong** |
| 689000 | 689400 | **Weak** |
| 702894 | 704338 | **NagC** |
| 703500 | 704000 | **Weak** |
| 707643 | 708608 | **NagC** |
| 707700 | 708700 | **Weak** |
| 765500 | 766500 | **Weak** |
| 847600 | 849000 | **Weak** |
| 852500 | 853300 | **Strong** |
| 986800 | 988000 | **Weak** |
| 997200 | 998200 | **Weak** |
| 1019000 | 1020500 | **Weak** |
| 1031700 | 1032200 | **Weak** |
| 1250600 | 1252985 | **Strong** |
| 1502800 | 1504400 | **Weak** |
| 1552500 | 1553000 | **Weak** |
| 1582233 | 1583181 | **NagC** |
| 1582300 | 1582900 | **Weak** |
| 1592000 | 1593000 | **Weak** |
| 1610200 | 1611400 | **Weak** |
| 1821447 | 1822024 | **NagC** |
| 1853252 | 1853933 | **NagC** |
| 1901200 | 1902300 | **Weak** |
| 1907400 | 1908600 | **Weak** |
| 1986000 | 1987000 | **Weak** |
| 2099400 | 2101200 | **Weak** |
| 2517400 | 2519200 | **Weak** |
| 2559600 | 2561200 | **Strong** |
| 2563000 | 2564500 | **Strong** |
| 2690800 | 2691800 | **Strong** |
| 2726000 | 2731300 | **Strong** |
| 2923900 | 2925000 | **Strong** |
| 2930100 | 2931000 | **Weak** |

| | | |
|---|---|---|
| 2970954 | 2971343 | **NagC** |
| 2978800 | 2979200 | **Weak** |
| 2984000 | 2984500 | **Weak** |
| 3055600 | 3056600 | **Strong** |
| 3087500 | 3089000 | **Weak** |
| 3087544 | 3088523 | **NagC** |
| 3118979 | 3119693 | **NagC** |
| 3119000 | 3119800 | **Weak** |
| 3184000 | 3185500 | **Weak** |
| 3278300 | 3279300 | **Weak** |
| 3281400 | 3282300 | **Weak** |
| 3317600 | 3319200 | **Strong** |
| 3423300 | 3429000 | **Strong** |
| 3580600 | 3583000 | **Strong** |
| 3707400 | 3708900 | **Weak** |
| 3719600 | 3720400 | **Strong** |
| 3742300 | 3743000 | **Weak** |
| 3914500 | 3916000 | **Weak** |
| 3915026 | 3915637 | **NagC** |
| 3949400 | 3951400 | **Strong** |
| 3957600 | 3958600 | **Weak** |
| 3982000 | 3983000 | **Strong** |
| 4035500 | 4040800 | **Strong** |
| 4057800 | 4058800 | **Strong** |
| 4171500 | 4172000 | **Strong** |
| 4539500 | 4540700 | **Strong** |
| 4539724 | 4540633 | **NagC** |
| 4603000 | 4606700 | **Strong** |
| 4640049 | 4641198 | **NagC** |

**Table S4: Table of Primers Used**. Contains primers used in Library-ChIP for cloning and generation of NGS libraries, as well as primers used in generation of tagged-TF strains.

| Name | Purpose | Strand | Sequence |
|---|---|---|---|
| pBAD Overhangs | Overhangs added to TF-specific primers for pBAD cloning. F overhang was added to the first 15-18 nt of TF sequence after the start codon (overhang includes start). R overhang was added to the reverse complement of the last 15-18 nt of TF sequence before the stop codon. | F | AGCAGGAGGAATTCACCATG |
| | | R | TCTTTGTAGTCGCCACCGCC |
| pBAD Test Primer | Primers for testing TF insertion via Sanger sequencing | F | ATGCCATAGCATTTTTATCC |
| | | R | GATTTAATCTGTATCAGG |
| oPool dsDNA Oligos | For PCR of Library-ChIP oligo-pool into dsDNA | F | GAGTAGGACGGATCCTCGAGCATGC |
| | | R | CGACGGCCAGTGCCAAGCTT |
| oPool Sanger Test Primers | Sanger Sequencing of Library-ChIP Inserts | F | CGCCTGGTTGCTACGCCTGAATAAG |
| | | R | GATCGGTGCGGGCCTCTTCG |
| oPool Universal LibPrep Primers | 1st PCR in Library-ChIP Lib. Prep. | F | TCGTCGGCAGCGTCAGATGTGTATAAGAGACAGNNNNNNNGAGTAGGACGGATCCTCGAGCATGC |
| | | R | GTCTCGTGGGCTCGGAGATGTGTATAAGAGACAGNNNNNNNNCGACGGCCAGTGCCAAGCTT |

| |
|---|
| SPA-FLAG Cloning Primers |
| SPA-FLAG Test Primers |
| V5 Cloning Primers |
| V5 Test Primer |

**Table S5: Table of Oligos Used in Library-ChIP.** Sequences were ordered as ssDNA and converted to dsDNA via primer extension for cloning into pAMD and use in Library-ChIP.

| Name | Sequence |
|---|---|
| PdhR_S1 | GGACGGATCCTCGAGCATGCAGCGTCACAGACATGAAATTGGTCAGACCAATTGACCTCGGCAAGTGG<br>CTTAAGACAGGAAAAGCTTGGCACTGGCCGTCG |
| PdhR_S2 | GGACGGATCCTCGAGCATGCAGCGTCACAGACATGAAATTGGTCAGACCATTTGACCTCGGCAAGTGG<br>CTTAAGACAGGAAAAGCTTGGCACTGGCCGTCG |
| PdhR_S3 | GGACGGATCCTCGAGCATGCAGCGTCACAGACATGAAATTGGTCTGACCAATTGACCTCGGCAAGTGG<br>CTTAAGACAGGAAAAGCTTGGCACTGGCCGTCG |
| PdhR_S4 | GGACGGATCCTCGAGCATGCAGCGTCACAGACATGAAATTGGTCTGACCATTTGACCTCGGCAAGTGG<br>CTTAAGACAGGAAAAGCTTGGCACTGGCCGTCG |
| PdhR_S5 | GGACGGATCCTCGAGCATGCAGCGTCACAGACATGAAAATGGTCAGACCAATTGACCTCGGCAAGTGG<br>CTTAAGACAGGAAAAGCTTGGCACTGGCCGTCG |
| PdhR_S6 | GGACGGATCCTCGAGCATGCAGCGTCACAGACATGAAAATGGTCAGACCATTTGACCTCGGCAAGTGG<br>CTTAAGACAGGAAAAGCTTGGCACTGGCCGTCG |
| PdhR_S7 | GGACGGATCCTCGAGCATGCAGCGTCACAGACATGAAAATGGTCTGACCAATTGACCTCGGCAAGTGG<br>CTTAAGACAGGAAAAGCTTGGCACTGGCCGTCG |
| PdhR_S8 | GGACGGATCCTCGAGCATGCAGCGTCACAGACATGAAAATGGTCTGACCATTTGACCTCGGCAAGTGG<br>CTTAAGACAGGAAAAGCTTGGCACTGGCCGTCG |
| PdhR_S9 | GGACGGATCCTCGAGCATGCAGCGTCACAGACATGAAAGTGGTCAGACCATTTGACCTCGGCAAGTGG<br>CTTAAGACAGGAAAAGCTTGGCACTGGCCGTCG |
| PdhR_S10 | GGACGGATCCTCGAGCATGCAGCGTCACAGACATGAAAGTGGTCAGACCAATTGACCTCGGCAAGTG<br>GCTTAAGACAGGAAAAGCTTGGCACTGGCCGTCG |
| PdhR_S11 | GGACGGATCCTCGAGCATGCAGCGTCACAGACATGAAAATGGTCAGACCACTTGACCTCGGCAAGTGG<br>CTTAAGACAGGAAAAGCTTGGCACTGGCCGTCG |
| PdhR_S12 | GGACGGATCCTCGAGCATGCAGCGTCACAGACATGAAAGTGGTCAGACCAACTGACCTCGGCAAGTG<br>GCTTAAGACAGGAAAAGCTTGGCACTGGCCGTCG |
| PdhR_S13 | GGACGGATCCTCGAGCATGCAGCGTCACAGACATGAAAGTGGTCTGACCAATTGGCCTCGGCAAGTG<br>GCTTAAGACAGGAAAAGCTTGGCACTGGCCGTCG |
| PdhR_S14 | GGACGGATCCTCGAGCATGCAGCGTCACAGACATGAAAGTGGTCCGACCATTTGACCTCGGCAAGTGG<br>CTTAAGACAGGAAAAGCTTGGCACTGGCCGTCG |
| PdhR_S15 | GGACGGATCCTCGAGCATGCAGCGTCACAGACATGAAAATGGTCTGACCAATTGTCCTCGGCAAGTGG<br>CTTAAGACAGGAAAAGCTTGGCACTGGCCGTCG |
| PdhR_S16 | GGACGGATCCTCGAGCATGCAGCGTCACAGACATGAATTTGGTCTGACCATTTGACCTCGGCAAGTGG<br>CTTAAGACAGGAAAAGCTTGGCACTGGCCGTCG |
| PdhR_S17 | GGACGGATCCTCGAGCATGCAGCGTCACAGACATGAAGTTGGTCAGACCATTTGCCCTCGGCAAGTGG<br>CTTAAGACAGGAAAAGCTTGGCACTGGCCGTCG |
| PdhR_S18 | GGACGGATCCTCGAGCATGCAGCGTCACAGACATGAAGTTGGTAAGACCAATTGACCTCGGCAAGTGG<br>CTTAAGACAGGAAAAGCTTGGCACTGGCCGTCG |
| PdhR_S19 | GGACGGATCCTCGAGCATGCAGCGTCACAGACATGAAAGTGGTCCGACCAATTGACCTCGGCAAGTG<br>GCTTAAGACAGGAAAAGCTTGGCACTGGCCGTCG |
| PdhR_S20 | GGACGGATCCTCGAGCATGCAGCGTCACAGACATGAAAGTGGTCTGACCATCTGACCTCGGCAAGTGG<br>CTTAAGACAGGAAAAGCTTGGCACTGGCCGTCG |
| PdhR_EC_Ref_1 | GGACGGATCCTCGAGCATGCAGCGTCACAGACATGAAATTGGTAAGACCAATTGACTTCGGCAAGTGG<br>CTTAAGACAGGAAAAGCTTGGCACTGGCCGTCG |

| | |
|---|---|
| PdhR_S22 | GGACGGATCCTCGAGCATGCAGCGTCACAGACATGAAAATGGTATGACCATTTGACATCGGCAAGTGG CTTAAGACAGGAAAAGCTTGGCACTGGCCGTCG |
| PdhR_S23 | GGACGGATCCTCGAGCATGCAGCGTCACAGACATGAAATTGGTCTGACCAGTTGGCCTCGGCAAGTGG CTTAAGACAGGAAAAGCTTGGCACTGGCCGTCG |
| PdhR_S24 | GGACGGATCCTCGAGCATGCAGCGTCACAGACATGAAATTGGTAAGACCATTTGATCTCGGCAAGTGG CTTAAGACAGGAAAAGCTTGGCACTGGCCGTCG |
| PdhR_S25 | GGACGGATCCTCGAGCATGCAGCGTCACAGACATGAAATTGGTCTGACCAGTTGAGCTCGGCAAGTGG CTTAAGACAGGAAAAGCTTGGCACTGGCCGTCG |
| PdhR_S26 | GGACGGATCCTCGAGCATGCAGCGTCACAGACATGAAAATGGTCTGACCAATTGGCATCGGCAAGTGG CTTAAGACAGGAAAAGCTTGGCACTGGCCGTCG |
| PdhR_S27 | GGACGGATCCTCGAGCATGCAGCGTCACAGACATGAACATGGTCTGACCATTTGTCCTCGGCAAGTGG CTTAAGACAGGAAAAGCTTGGCACTGGCCGTCG |
| PdhR_S28 | GGACGGATCCTCGAGCATGCAGCGTCACAGACATGAAAGTGGTAAGACCATTTGTCCTCGGCAAGTGG CTTAAGACAGGAAAAGCTTGGCACTGGCCGTCG |
| PdhR_S29 | GGACGGATCCTCGAGCATGCAGCGTCACAGACATGAAAATGGTCAGACCAATTGCCGTCGGCAAGTG GCTTAAGACAGGAAAAGCTTGGCACTGGCCGTCG |
| PdhR_S30 | GGACGGATCCTCGAGCATGCAGCGTCACAGACATGAAATTGGTATGACCAATTGACTTCGGCAAGTGG CTTAAGACAGGAAAAGCTTGGCACTGGCCGTCG |
| PdhR_S31 | GGACGGATCCTCGAGCATGCAGCGTCACAGACATGAAGTTGGTCAGACCAGATGACCTCGGCAAGTG GCTTAAGACAGGAAAAGCTTGGCACTGGCCGTCG |
| PdhR_S32 | GGACGGATCCTCGAGCATGCAGCGTCACAGACATGAAATTGGTCATACCAGTTGCTCTCGGCAAGTGG CTTAAGACAGGAAAAGCTTGGCACTGGCCGTCG |
| PdhR_S33 | GGACGGATCCTCGAGCATGCAGCGTCACAGACATGAAAGTGGTAAGACCAAATGAGCTCGGCAAGTG GCTTAAGACAGGAAAAGCTTGGCACTGGCCGTCG |
| PdhR_S34 | GGACGGATCCTCGAGCATGCAGCGTCACAGACATGAAATTGGTATTACCACTTGCCCTCGGCAAGTGG CTTAAGACAGGAAAAGCTTGGCACTGGCCGTCG |
| PdhR_S35 | GGACGGATCCTCGAGCATGCAGCGTCACAGACATGAAGTTGGTCTGACCAGTTGACATCGGCAAGTGG CTTAAGACAGGAAAAGCTTGGCACTGGCCGTCG |
| PdhR_S36 | GGACGGATCCTCGAGCATGCAGCGTCACAGACATGAAATTGGTATGACCAAGTGTCCTCGGCAAGTGG CTTAAGACAGGAAAAGCTTGGCACTGGCCGTCG |
| PdhR_S37 | GGACGGATCCTCGAGCATGCAGCGTCACAGACATGAATATGGTCAGACCATTTGCCTTCGGCAAGTGG CTTAAGACAGGAAAAGCTTGGCACTGGCCGTCG |
| PdhR_S38 | GGACGGATCCTCGAGCATGCAGCGTCACAGACATGAAGTTGGTAAGACCATTTGGGCTCGGCAAGTG GCTTAAGACAGGAAAAGCTTGGCACTGGCCGTCG |
| PdhR_S39 | GGACGGATCCTCGAGCATGCAGCGTCACAGACATGAAGTTGGTCTTACCATCTGCCCTCGGCAAGTGG CTTAAGACAGGAAAAGCTTGGCACTGGCCGTCG |
| PdhR_S40 | GGACGGATCCTCGAGCATGCAGCGTCACAGACATGAAAATGGTATTACCATTTGCGCTCGGCAAGTGG CTTAAGACAGGAAAAGCTTGGCACTGGCCGTCG |
| PdhR_S41 | GGACGGATCCTCGAGCATGCAGCGTCACAGACATGAAGCTGGTCTGACCACTTGGCTTCGGCAAGTGG CTTAAGACAGGAAAAGCTTGGCACTGGCCGTCG |
| PdhR_S42 | GGACGGATCCTCGAGCATGCAGCGTCACAGACATGAAGATGGTATGACCATTTGGTTTCGGCAAGTGG CTTAAGACAGGAAAAGCTTGGCACTGGCCGTCG |
| PdhR_S43 | GGACGGATCCTCGAGCATGCAGCGTCACAGACATGAACTTGGTCCGACCAATTGAATTCGGCAAGTGG CTTAAGACAGGAAAAGCTTGGCACTGGCCGTCG |
| PdhR_S44 | GGACGGATCCTCGAGCATGCAGCGTCACAGACATGAACGTGGTCATACCATTTGGCTTCGGCAAGTGG CTTAAGACAGGAAAAGCTTGGCACTGGCCGTCG |

| | |
|---|---|
| PdhR_S45 | GGACGGATCCTCGAGCATGCAGCGTCACAGACATGAACATGGTTATACCATATGACCTCGGCAAGTGGCTTAAGACAGGAAAAGCTTGGCACTGGCCGTCG |
| PdhR_S46 | GGACGGATCCTCGAGCATGCAGCGTCACAGACATGAATTTGGTCAAACCACTTGCCCTCGGCAAGTGGCTTAAGACAGGAAAAGCTTGGCACTGGCCGTCG |
| PdhR_S47 | GGACGGATCCTCGAGCATGCAGCGTCACAGACATGAAAATGGTCACACCATCTGAACTCGGCAAGTGGCTTAAGACAGGAAAAGCTTGGCACTGGCCGTCG |
| PdhR_S48 | GGACGGATCCTCGAGCATGCAGCGTCACAGACATGAAGTTGGTCTGACCACCTGAAATCGGCAAGTGGCTTAAGACAGGAAAAGCTTGGCACTGGCCGTCG |
| PdhR_S49 | GGACGGATCCTCGAGCATGCAGCGTCACAGACATGAAAGTGGTTTTACCAAGTGACCTCGGCAAGTGGCTTAAGACAGGAAAAGCTTGGCACTGGCCGTCG |
| PdhR_S50 | GGACGGATCCTCGAGCATGCAGCGTCACAGACATGAAGTTGGTAAGACCATTTGTGATCGGCAAGTGGCTTAAGACAGGAAAAGCTTGGCACTGGCCGTCG |
| AllR_S1 | GGACGGATCCTCGAGCATGCTTGACAAAATTTGAAAGTTGGAAAATTTTTCCAATAAATAGAGGTAGGAATAAAATGGCAAAAGCTTGGCACTGGCCGTCG |
| AllR_EC_Ref_1 | GGACGGATCCTCGAGCATGCTTGACAAAATTTGAAAGTTGGAAAAATTTTCCAATAAATAGAGGTAGGAATAAAATGGCAAAAGCTTGGCACTGGCCGTCG |
| AllR_S3 | GGACGGATCCTCGAGCATGCTTGACAAAATTTGAAAGGTGGAAAAATTTTCCAATAAATAGAGGTAGGAATAAAATGGCAAAAGCTTGGCACTGGCCGTCG |
| AllR_S4 | GGACGGATCCTCGAGCATGCTTGACAAAATTTGAAAGGTGGAAAATTTTTCCAATAAATAGAGGTAGGAATAAAATGGCAAAAGCTTGGCACTGGCCGTCG |
| AllR_S5 | GGACGGATCCTCGAGCATGCTTGACAAAATTTGAAAGTTGGAAATTTTTTCCAATAAATAGAGGTAGGAATAAAATGGCAAAAGCTTGGCACTGGCCGTCG |
| AllR_S6 | GGACGGATCCTCGAGCATGCTTGACAAAATTTGAAAGTTGGAAAATATTTCCAATAAATAGAGGTAGGAATAAAATGGCAAAAGCTTGGCACTGGCCGTCG |
| AllR_S7 | GGACGGATCCTCGAGCATGCTTGACAAAATTTGAAAGTTGGAAATATTTTCCAATAAATAGAGGTAGGAATAAAATGGCAAAAGCTTGGCACTGGCCGTCG |
| AllR_S8 | GGACGGATCCTCGAGCATGCTTGACAAAATTTGAAAGTTGGAAACTTTTTCCAATAAATAGAGGTAGGAATAAAATGGCAAAAGCTTGGCACTGGCCGTCG |
| AllR_S9 | GGACGGATCCTCGAGCATGCTTGACAAAATTTGAAAGTTGGAAAAAATTTCCAATAAATAGAGGTAGGAATAAAATGGCAAAAGCTTGGCACTGGCCGTCG |
| AllR_S10 | GGACGGATCCTCGAGCATGCTTGACAAAATTTGAAAGATGGAAAAATTTTCCAATAAATAGAGGTAGGAATAAAATGGCAAAAGCTTGGCACTGGCCGTCG |
| AllR_S11 | GGACGGATCCTCGAGCATGCTTGACAAAATTTGAAAGTTGGAAATATATTCCAATAAATAGAGGTAGGAATAAAATGGCAAAAGCTTGGCACTGGCCGTCG |
| AllR_S12 | GGACGGATCCTCGAGCATGCTTGACAAAATTTGAAAGTTGGAAATAATTTCCAATAAATAGAGGTAGGAATAAAATGGCAAAAGCTTGGCACTGGCCGTCG |
| AllR_S13 | GGACGGATCCTCGAGCATGCTTGACAAAATTTGAAAGTTGGAAATGTTTTCCAATAAATAGAGGTAGGAATAAAATGGCAAAAGCTTGGCACTGGCCGTCG |
| AllR_S14 | GGACGGATCCTCGAGCATGCTTGACAAAATTTGAAAGATGGAAATATTTTCCAATAAATAGAGGTAGGAATAAAATGGCAAAAGCTTGGCACTGGCCGTCG |
| AllR_S15 | GGACGGATCCTCGAGCATGCTTGACAAAATTTGAAAGTTGGAATATATTTCCAATAAATAGAGGTAGGAATAAAATGGCAAAAGCTTGGCACTGGCCGTCG |
| AllR_S16 | GGACGGATCCTCGAGCATGCTTGACAAAATTTGAAAGTGGGAAAATTTTCCCAATAAATAGAGGTAGGAATAAAATGGCAAAAGCTTGGCACTGGCCGTCG |
| AllR_S17 | GGACGGATCCTCGAGCATGCTTGACAAAATTTGAAAGTCGGAAAATATTTCCAATAAATAGAGGTAGGAATAAAATGGCAAAAGCTTGGCACTGGCCGTCG |

| | |
|---|---|
| AllR_S18 | GGACGGATCCTCGAGCATGCTTGACAAAATTTGAAAGATGGAAACTTTTTCCAATAAATAGAGGTAGG AATAAAATGGCAAAAGCTTGGCACTGGCCGTCG |
| AllR_S19 | GGACGGATCCTCGAGCATGCTTGACAAAATTTGAAAGTTGGAAAAGATTTCCAATAAATAGAGGTAGG AATAAAATGGCAAAAGCTTGGCACTGGCCGTCG |
| AllR_S20 | GGACGGATCCTCGAGCATGCTTGACAAAATTTGAAAGTTGGAAACTGTTTCCAATAAATAGAGGTAGG AATAAAATGGCAAAAGCTTGGCACTGGCCGTCG |
| AllR_S21 | GGACGGATCCTCGAGCATGCTTGACAAAATTTGAAAGTTGGAAGATATGCCCAATAAATAGAGGTAGG AATAAAATGGCAAAAGCTTGGCACTGGCCGTCG |
| AllR_S22 | GGACGGATCCTCGAGCATGCTTGACAAAATTTGAAAGGTGGACACAGATTCCAATAAATAGAGGTAGG AATAAAATGGCAAAAGCTTGGCACTGGCCGTCG |
| AllR_S23 | GGACGGATCCTCGAGCATGCTTGACAAAATTTGAAAGGTGGAACAGTTATCCAATAAATAGAGGTAGG AATAAAATGGCAAAAGCTTGGCACTGGCCGTCG |
| AllR_S24 | GGACGGATCCTCGAGCATGCTTGACAAAATTTGAAAGTAGGAAATGGATTCCAATAAATAGAGGTAGG AATAAAATGGCAAAAGCTTGGCACTGGCCGTCG |
| AllR_S25 | GGACGGATCCTCGAGCATGCTTGACAAAATTTGAAAGTAGGAATAAGTTGCCAATAAATAGAGGTAGG AATAAAATGGCAAAAGCTTGGCACTGGCCGTCG |
| AllR_S26 | GGACGGATCCTCGAGCATGCTTGACAAAATTTGAAAGGTGGACGTAATTTCCAATAAATAGAGGTAGG AATAAAATGGCAAAAGCTTGGCACTGGCCGTCG |
| AllR_S27 | GGACGGATCCTCGAGCATGCTTGACAAAATTTGAAAGTTGGAAAAGGAATCCAATAAATAGAGGTAGG AATAAAATGGCAAAAGCTTGGCACTGGCCGTCG |
| AllR_S28 | GGACGGATCCTCGAGCATGCTTGACAAAATTTGAAAGGCGGAAGTGGTTTCCAATAAATAGAGGTAGG AATAAAATGGCAAAAGCTTGGCACTGGCCGTCG |
| AllR_S29 | GGACGGATCCTCGAGCATGCTTGACAAAATTTGAAAGTCGGAATCTGCTTCCAATAAATAGAGGTAGG AATAAAATGGCAAAAGCTTGGCACTGGCCGTCG |
| AllR_S30 | GGACGGATCCTCGAGCATGCTTGACAAAATTTGAAAGAGGGAATATGCTTCCAATAAATAGAGGTAGG AATAAAATGGCAAAAGCTTGGCACTGGCCGTCG |
| GlnG_S1 | GGACGGATCCTCGAGCATGCTTCAATGCACTATATTAGTGCAACAAACCTGGCGTGCACCATGATGATCA TTCCATCAGGTAAGCTTGGCACTGGCCGTCG |
| GlnG_S2 | GGACGGATCCTCGAGCATGCTTCAATGCACTATTTAGTGCAACAAACCTGGCGTGCACCATGATGATC ATTCCATCAGGTAAGCTTGGCACTGGCCGTCG |
| GlnG_S3 | GGACGGATCCTCGAGCATGCTTCAATGCACTAATATAGTGCAACAAACCTGGCGTGCACCATGATGATC ATTCCATCAGGTAAGCTTGGCACTGGCCGTCG |
| GlnG_S4 | GGACGGATCCTCGAGCATGCTTCAATGCACTAAAATAGTGCAACAAACCTGGCGTGCACCATGATGATC ATTCCATCAGGTAAGCTTGGCACTGGCCGTCG |
| GlnG_S5 | GGACGGATCCTCGAGCATGCTTCAATGCACTAATTTAGTGCAACAAACCTGGCGTGCACCATGATGATC ATTCCATCAGGTAAGCTTGGCACTGGCCGTCG |
| GlnG_S6 | GGACGGATCCTCGAGCATGCTTCAATGCACTAAATTAGTGCAACAAACCTGGCGTGCACCATGATGATC ATTCCATCAGGTAAGCTTGGCACTGGCCGTCG |
| GlnG_S7 | GGACGGATCCTCGAGCATGCTTCAATGCACTATTATAGTGCAACAAACCTGGCGTGCACCATGATGATCA TTCCATCAGGTAAGCTTGGCACTGGCCGTCG |
| GlnG_S8 | GGACGGATCCTCGAGCATGCTTCAATGCACTATAATAGTGCAACAAACCTGGCGTGCACCATGATGATC ATTCCATCAGGTAAGCTTGGCACTGGCCGTCG |
| GlnG_S9 | GGACGGATCCTCGAGCATGCTTCAATGCACCAATATAGTGCAACAAACCTGGCGTGCACCATGATGATC ATTCCATCAGGTAAGCTTGGCACTGGCCGTCG |
| GlnG_S10 | GGACGGATCCTCGAGCATGCTTCAATGCACCAAAATAGTGCAACAAACCTGGCGTGCACCATGATGATC ATTCCATCAGGTAAGCTTGGCACTGGCCGTCG |

| | |
|---|---|
| GlnG_S11 | GGACGGATCCTCGAGCATGCTTCAATGCTCAATAATTGTGCAACAAACCTGGCGTGCACCATGATGATC ATTCCATCAGGTAAGCTTGGCACTGGCCGTCG |
| GlnG_S12 | GGACGGATCCTCGAGCATGCTTCAATGCACTACTGTCGTGCAACAAACCTGGCGTGCACCATGATGATC ATTCCATCAGGTAAGCTTGGCACTGGCCGTCG |
| GlnG_S13 | GGACGGATCCTCGAGCATGCTTCAATGCACTATCTCAGTGCAACAAACCTGGCGTGCACCATGATGATC ATTCCATCAGGTAAGCTTGGCACTGGCCGTCG |
| GlnG_S14 | GGACGGATCCTCGAGCATGCTTCAATGGTGCATGTTCGTGCAACAAACCTGGCGTGCACCATGATGATC ATTCCATCAGGTAAGCTTGGCACTGGCCGTCG |
| GlnG_S15 | GGACGGATCCTCGAGCATGCTTCAATGCACAAATTCGGTGCAACAAACCTGGCGTGCACCATGATGATC ATTCCATCAGGTAAGCTTGGCACTGGCCGTCG |
| GlnG_S16 | GGACGGATCCTCGAGCATGCTTCAATGCGCTATTATAGTGCAACAAACCTGGCGTGCACCATGATGATC ATTCCATCAGGTAAGCTTGGCACTGGCCGTCG |
| GlnG_S17 | GGACGGATCCTCGAGCATGCTTCAATGCACCAAGACAGTGCAACAAACCTGGCGTGCACCATGATGAT CATTCCATCAGGTAAGCTTGGCACTGGCCGTCG |
| GlnG_S18 | GGACGGATCCTCGAGCATGCTTCAATGCGCTATAATAGTGCAACAAACCTGGCGTGCACCATGATGATC ATTCCATCAGGTAAGCTTGGCACTGGCCGTCG |
| GlnG_S19 | GGACGGATCCTCGAGCATGCTTCAATGCTCTATTACAGTGCAACAAACCTGGCGTGCACCATGATGATC ATTCCATCAGGTAAGCTTGGCACTGGCCGTCG |
| GlnG_S20 | GGACGGATCCTCGAGCATGCTTCAATGGTGCATGTATGTGCAACAAACCTGGCGTGCACCATGATGATC ATTCCATCAGGTAAGCTTGGCACTGGCCGTCG |
| GlnG_S21 | GGACGGATCCTCGAGCATGCTTCAATGCTCTGTAGGTGTGCAACAAACCTGGCGTGCACCATGATGATC ATTCCATCAGGTAAGCTTGGCACTGGCCGTCG |
| GlnG_S22 | GGACGGATCCTCGAGCATGCTTCAATGCTTCAAATACGTGCAACAAACCTGGCGTGCACCATGATGATC ATTCCATCAGGTAAGCTTGGCACTGGCCGTCG |
| GlnG_S23 | GGACGGATCCTCGAGCATGCTTCAATGGCCCTATTTAGTGCAACAAACCTGGCGTGCACCATGATGATC ATTCCATCAGGTAAGCTTGGCACTGGCCGTCG |
| GlnG_S24 | GGACGGATCCTCGAGCATGCTTCAATGCGGCATTTCTGTGCAACAAACCTGGCGTGCACCATGATGATC ATTCCATCAGGTAAGCTTGGCACTGGCCGTCG |
| GlnG_S25 | GGACGGATCCTCGAGCATGCTTCAATGCCCGGATACGGTGCAACAAACCTGGCGTGCACCATGATGATC ATTCCATCAGGTAAGCTTGGCACTGGCCGTCG |
| GlnG_S26 | GGACGGATCCTCGAGCATGCTTCAATGCCACATTGTAGTGCAACAAACCTGGCGTGCACCATGATGATC ATTCCATCAGGTAAGCTTGGCACTGGCCGTCG |
| GlnG_S27 | GGACGGATCCTCGAGCATGCTTCAATGTGCCACGTTAGTGCAACAAACCTGGCGTGCACCATGATGATC ATTCCATCAGGTAAGCTTGGCACTGGCCGTCG |
| GlnG_S28 | GGACGGATCCTCGAGCATGCTTCAATGGTACAAATTGGTGCAACAAACCTGGCGTGCACCATGATGATC ATTCCATCAGGTAAGCTTGGCACTGGCCGTCG |
| GlnG_S29 | GGACGGATCCTCGAGCATGCTTCAATGCCGTAAAAGAGTGCAACAAACCTGGCGTGCACCATGATGAT CATTCCATCAGGTAAGCTTGGCACTGGCCGTCG |
| GlnG_S30 | GGACGGATCCTCGAGCATGCTTCAATGCTCGCGAACAGTGCAACAAACCTGGCGTGCACCATGATGAT CATTCCATCAGGTAAGCTTGGCACTGGCCGTCG |
| GlnG_EC_Ref_1 | GGACGGATCCTCGAGCATGCGCAATACCCTGAAGATGCACCATTCTGGGGCACCAATCTGGTGCGCTAA AATTGTGCACTCAAGCTTGGCACTGGCCGTC |
| GlnG_EC_Ref_3 | GGACGGATCCTCGAGCATGCGCCCTGACTTTGGGCAGCAATGCTTCAAGTTGGGGCGCTAAAATGGG GCATTGTTTGACGTAAGCTTGGCACTGGCCGTC |
| AllR_EC_Ref_2 | GGACGGATCCTCGAGCATGCGTGGACTAAATCTAGTTTTGGAAAAATATTCCAACTTTTGTATTGATGTT GTTCTCTTAAGAAGCTTGGCACTGGCCGTC |

| | |
|---|---|
| AllR_EC_Ref_3 | GGACGGATCCTCGAGCATGCGAAGATGCCGCGAAAATTCTTTCCTTTGGCGCGGATAAAATTTCCATCAACTCTCCTGCGCAAGCTTGGCACTGGCCGTC |
| PdhR_EC_Ref_2 | GGACGGATCCTCGAGCATGCACAAAAGGGGAGTGCTGAAGGAGTCTGGGCGGGCAATTGGTATAACCAATGTGAAATAAAAAAGCTTGGCACTGGCCGTC |
| PdhR_EC_Ref_3 | GGACGGATCCTCGAGCATGCCAACAAAACTTGATTAACATCAATTTTGGTATGACCAATGCACCATTCATGTTATTCTCAAAAGCTTGGCACTGGCCGTC |

**Table S6: Table of Oligos Used in BLI.** A biotinylated-forward strand oligo and corresponding reverse oligo were annealed and used in BLI.

| Name | Sequence |
|---|---|
| **AllR_EC_Ref_1_F** | /5BiosG/TTGACAAAATTTGAAAGTTGGAAAAATTTTCCAATAAATAGAGGTAGGAATAAAATGGCAA |
| **AllR_EC_Ref_2_F** | /5BiosG/GTGGACTAAATCTAGTTTTGGAAAAATATTCCAACTTTTGTATTGATGTTGTTCTCTTAAG |
| **AllR_EC_Ref_3_F** | /5BiosG/GAAGATGCCGCGAAAATTCTTTCCTTTGGCGCGGATAAAATTTCCATCAACTCTCCTGCGC |
| **AllR_EC_Ref_4_F** | /5BiosG/CAAAATATGTCATCTCTGGCGGAAAAGTTTTCCACTGACAACGCGGGCATTGCCTACTTAA |
| **AllR_EC_Ref_5_F** | /5BiosG/CGCGGCTTGCGCCATTCTGACCGCGCTCTGCACGGAAAGATTTTTCAAAACGATAATCGAT |
| **AllR_EC_Ref_6_F** | /5BiosG/CTCGTTCAGGGGGCTGGTAATATTGGAAAAATTGCCCGTTCTTTAGCTGAAATCAAACTGA |
| **AllR_S11_F** | /5BiosG/TTGACAAAATTTGAAAGTTGGAAATATATTCCAATAAATAGAGGTAGGAATAAAATGGCAA |
| **AllR_S21_F** | /5BiosG/TTGACAAAATTTGAAAGTTGGAAGATATGCCCAATAAATAGAGGTAGGAATAAAATGGCAA |
| **GlnG_EC_Ref_2_F** | /5BiosG/ATGCATTTCAATGCACCACGATAGTGCAACAAACCTGGCGTGCACCATGATGATCATTCCA |
| **GlnG_EC_Ref_3_F** | /5BiosG/GCCCTGACTTTGGGCAGCAATGCTTCAAGTTGGGGCGCTAAAATGGGGCATTGTTTGACGT |
| **GlnG_S1_F** | /5BiosG/TTCAATGCACTATATTAGTGCAACAAACCTGGCGTGCACCATGATGATCATTCCATCAGGT |
| **GlnG_S11_F** | /5BiosG/TTCAATGCTCAATAATTGTGCAACAAACCTGGCGTGCACCATGATGATCATTCCATCAGGT |
| **GlnG_S21_F** | /5BiosG/TTCAATGCTCTGTAGGTGTGCAACAAACCTGGCGTGCACCATGATGATCATTCCATCAGGT |
| **Nac_EC_Ref_1_F** | /5BiosG/CATTCGCAACCTATAAGTAAATCCAATGGAACTCATCATAAATGAGACTTTTACCTTATGA |
| **Nac_EC_Ref_2_F** | /5BiosG/CCACACTGTTACATAAGTTAATCTTAGGTGAAATACCGACTTCATAACTTTTACGCATTAT |
| **Nac_EC_Ref_3_F** | /5BiosG/CTTAAATCTTGCATTGGTTTTCCTTATATTCAGACTTTGTTATATAAGGCTTTCGTCTTTG |
| **Nac_S1_430304_F** | /5BiosG/CATTCGCCGGCTATAAGTAAATCTTATGGAACTCATCATAAATGAGACTTTTACCTTATGA |
| **Nac_S11_364632_F** | /5BiosG/CATTCGCCCGCTATAAGTAAACCCTATGGAACTCATCATAAATGAGACTTTTACCTTATGA |
| **Nac_S21_1022037_F** | /5BiosG/CATTCGCTTGCTATAAGTAGACCCAATGGAACTCATCATAAATGAGACTTTTACCTTATGA |
| **Nac_S31_603054_F** | /5BiosG/CATTCGCGCATTATAAGTAATGGTCATGGAACTCATCATAAATGAGACTTTTACCTTATGA |
| **PdhR_EC_Ref_1_F** | /5BiosG/AGCGTCACAGACATGAAATTGGTAAGACCAATTGACTTCGGCAAGTGGCTTAAGACAGGAA |
| **PdhR_EC_Ref_2_F** | /5BiosG/ACAAAAGGGGAGTGCTGAAGGAGTCTGGGCGGGCAATTGGTATAACCAATGTGAAATAAAA |
| **PdhR_EC_Ref_3_F** | /5BiosG/CAACAAAACTTGATTAACATCAATTTTGGTATGACCAATGCACCATTCATGTTATTCTCAA |
| **PdhR_S1_F** | /5BiosG/AGCGTCACAGACATGAAATTGGTCAGACCAATTGACCTCGGCAAGTGGCTTAAGACAGGAA |
| **PdhR_S11_F** | /5BiosG/AGCGTCACAGACATGAAAATGGTCAGACCACTTGACCTCGGCAAGTGGCTTAAGACAGGAA |
| **PdhR_S27_F** | /5BiosG/AGCGTCACAGACATGAACATGGTCTGACCATTTGTCCTCGGCAAGTGGCTTAAGACAGGAA |
| **PdhR_S31_F** | /5BiosG/AGCGTCACAGACATGAAGTTGGTCAGACCAGATGACCTCGGCAAGTGGCTTAAGACAGGAA |
| **PdhR_S41_F** | /5BiosG/AGCGTCACAGACATGAAGCTGGTCTGACCACTTGGCTTCGGCAAGTGGCTTAAGACAGGAA |
| **PdhR_S8_F** | /5BiosG/AGCGTCACAGACATGAAAATGGTCTGACCATTTGACCTCGGCAAGTGGCTTAAGACAGGAA |
| **UlaR_EC_Ref_1_F** | /5BiosG/GGTAACTTAGCGTCTCGTTGTGATTAAGTTTGACGAGGAATGGAATGCGATGCGCATAACG |
| **UlaR_EC_Ref_2_F** | /5BiosG/GTTCAATTCGAATCAATATGTGATTGGTTTTGATTAATCCTGACACTATTTTTTCAGGAAG |
| **UlaR_EC_Ref_3_F** | /5BiosG/AAGAGGAGTTAAATTTTATACAGTATAATCATAATATTGCAGCAAGGTGGTTATAATTGAA |
| **UlaR_S1_147397_F** | /5BiosG/GGTAACTTAGCGTCTCGTTGTGAGTAATTTTGACAAGGAATGGAATGCGATGCGCATAACG |
| **UlaR_S11_196037_F** | /5BiosG/GGTAACTTAGCGTCTCGTTGTGAGTATTTCTGACAAGGAATGGAATGCGATGCGCATAACG |
| **UlaR_S21_502983_F** | /5BiosG/GGTAACTTAGCGTCTCGTTGTGCTTAGGTATGACGAGGAATGGAATGCGATGCGCATAACG |
| **AllR_neg_F** | /5BiosG/GTAAAGTAGGAACGGTATTAATAGAAATGATACTAAAGTACGAAATAATAAACTGGAATTT |
| **GlnG_neg_F** | /5BiosG/TTCTGATCACCGACATGAAAGCTAACAGGAACGTTAAGTGCCTTAGCTCCGGCTACCGTGC |
| **PdhR_neg_F** | /5BiosG/AGATCTCCATCCTTGGGATATGGCTACCAGAAAGTAAAGGGACGATACAAGTCAAGTGAGA |
| **AllR_EC_Ref_1_R** | TTGCCATTTTATTCCTACCTCTATTTATTGGAAAATTTTTCCAACTTTCAAATTTTGTCAA |

| | |
|---|---|
| **AllR_EC_Ref_2_R** | CTTAAGAGAACAACATCAATACAAAAGTTGGAATATTTTTCCAAAACTAGATTTAGTCCAC |
| **AllR_EC_Ref_3_R** | GCGCAGGAGAGTTGATGGAAATTTTATCCGCGCCAAAGGAAAGAATTTTCGCGGCATCTTC |
| **AllR_EC_Ref_4_R** | TTAAGTAGGCAATGCCCGCGTTGTCAGTGGAAAACTTTTCCGCCAGAGATGACATATTTTG |
| **AllR_EC_Ref_5_R** | ATCGATTATCGTTTTGAAAAATCTTTCCGTGCAGAGCGCGGTCAGAATGGCGCAAGCCGCG |
| **AllR_EC_Ref_6_R** | TCAGTTTGATTTCAGCTAAAGAACGGGCAATTTTTCCAATATTACCAGCCCCCTGAACGAG |
| **AllR_S11_R** | TTGCCATTTTATTCCTACCTCTATTTATTGGAATATATTTCCAACTTTCAAATTTTGTCAA |
| **AllR_S21_R** | TTGCCATTTTATTCCTACCTCTATTTATTGGGCATATCTTCCAACTTTCAAATTTTGTCAA |
| **GlnG_EC_Ref_2_R** | TGGAATGATCATCATGGTGCACGCCAGGTTTGTTGCACTATCGTGGTGCATTGAAATGCAT |
| **GlnG_EC_Ref_3_R** | ACGTCAAACAATGCCCCATTTTAGCGCCCCAACTTGAAGCATTGCTGCCCAAAGTCAGGGC |
| **GlnG_S1_R** | ACCTGATGGAATGATCATCATGGTGCACGCCAGGTTTGTTGCACTAATATAGTGCATTGAA |
| **GlnG_S11_R** | ACCTGATGGAATGATCATCATGGTGCACGCCAGGTTTGTTGCACAATTATTGAGCATTGAA |
| **GlnG_S21_R** | ACCTGATGGAATGATCATCATGGTGCACGCCAGGTTTGTTGCACACCTACAGAGCATTGAA |
| **Nac_EC_Ref_1_R** | TCATAAGGTAAAAGTCTCATTTATGATGAGTTCCATTGGATTTACTTATAGGTTGCGAATG |
| **Nac_EC_Ref_2_R** | ATAATGCGTAAAAGTTATGAAGTCGGTATTTCACCTAAGATTAACTTATGTAACAGTGTGG |
| **Nac_EC_Ref_3_R** | CAAAGACGAAAGCCTTATATAACAAAGTCTGAATATAAGGAAAACCAATGCAAGATTTAAG |
| **Nac_S1_430304_R** | TCATAAGGTAAAAGTCTCATTTATGATGAGTTCCATAAGATTTACTTATAGCCGGCGAATG |
| **Nac_S11_364632_R** | TCATAAGGTAAAAGTCTCATTTATGATGAGTTCCATAGGGTTTACTTATAGCGGGCGAATG |
| **Nac_S21_1022037_R** | TCATAAGGTAAAAGTCTCATTTATGATGAGTTCCATTGGGTCTACTTATAGCAAGCGAATG |
| **Nac_S31_603054_R** | TCATAAGGTAAAAGTCTCATTTATGATGAGTTCCATGACCATTACTTATAATGCGCGAATG |
| **PdhR_EC_Ref_1_R** | TTCCTGTCTTAAGCCACTTGCCGAAGTCAATTGGTCTTACCAATTTCATGTCTGTGACGCT |
| **PdhR_EC_Ref_2_R** | TTTTATTTCACATTGGTTATACCAATTGCCCGCCCAGACTCCTTCAGCACTCCCCTTTTGT |
| **PdhR_EC_Ref_3_R** | TTGAGAATAACATGAATGGTGCATTGGTCATACCAAAATTGATGTTAATCAAGTTTTGTTG |
| **PdhR_S1_R** | TTCCTGTCTTAAGCCACTTGCCGAGGTCAATTGGTCTGACCAATTTCATGTCTGTGACGCT |
| **PdhR_S11_R** | TTCCTGTCTTAAGCCACTTGCCGAGGTCAAGTGGTCTGACCATTTTCATGTCTGTGACGCT |
| **PdhR_S27_R** | TTCCTGTCTTAAGCCACTTGCCGAGGACAAATGGTCAGACCATGTTCATGTCTGTGACGCT |
| **PdhR_S31_R** | TTCCTGTCTTAAGCCACTTGCCGAGGTCATCTGGTCTGACCAACTTCATGTCTGTGACGCT |
| **PdhR_S41_R** | TTCCTGTCTTAAGCCACTTGCCGAAGCCAAGTGGTCAGACCAGCTTCATGTCTGTGACGCT |
| **PdhR_S8_R** | TTCCTGTCTTAAGCCACTTGCCGAGGTCAAATGGTCAGACCATTTTCATGTCTGTGACGCT |
| **UlaR_EC_Ref_1_R** | CGTTATGCGCATCGCATTCCATTCCTCGTCAAACTTAATCACAACGAGACGCTAAGTTACC |
| **UlaR_EC_Ref_2_R** | CTTCCTGAAAAAATAGTGTCAGGATTAATCAAAACCAATCACATATTGATTCGAATTGAAC |
| **UlaR_EC_Ref_3_R** | TTCAATTATAACCACCTTGCTGCAATATTATGATTATACTGTATAAAATTTAACTCCTCTT |
| **UlaR_S1_147397_R** | CGTTATGCGCATCGCATTCCATTCCTTGTCAAAATTACTCACAACGAGACGCTAAGTTACC |
| **UlaR_S11_196037_R** | CGTTATGCGCATCGCATTCCATTCCTTGTCAGAAATACTCACAACGAGACGCTAAGTTACC |
| **UlaR_S21_502983_R** | CGTTATGCGCATCGCATTCCATTCCTCGTCATACCTAAGCACAACGAGACGCTAAGTTACC |
| **AllR_neg_R** | AAATTCCAGTTTATTATTTCGTACTTTAGTATCATTTCTATTAATACCGTTCCTACTTTAC |
| **GlnG_neg_R** | GCACGGTAGCCGGAGCTAAGGCACTTAACGTTCCTGTTAGCTTTCATGTCGGTGATCAGAA |
| **PdhR_neg_R** | TCTCACTTGACTTGTATCGTCCCTTTACTTTCTGGTAGCCATATCCCAAGGATGGAGATCT |

**Table S7: Table of TF-DNA Pairs Tested in BLI**. Table lists the DNA sequences TFs were tested against and the fit $K_D$ and binding energy of each pair.

| TF | BLI Oligo | $K_D$ (nM) | Energy ($k_bT$) |
|---|---|---|---|
| **AllR** | **AllR neg** | 138.8 | 15.79 |
| **AllR** | **AllR EC Ref 1** | 9.61 | 18.46 |
| **AllR** | **AllR EC Ref 2** | 8.69 | 18.56 |
| **AllR** | **AllR EC Ref 3** | 45.85 | 16.9 |
| **AllR** | **AllR EC Ref 4** | 53.13 | 16.75 |
| **AllR** | **AllR EC Ref 5** | 98.9 | 16.13 |
| **AllR** | **AllR EC Ref 6** | 231.6 | 15.28 |
| **AllR** | **AllR S11** | 41.87 | 16.99 |
| **AllR** | **AllR S21** | 423.4 | 14.68 |
| **AllR** | **argP EC Ref 1** | 852.3 | 13.98 |
| **AllR** | **GlnG neg** | 1960 | 13.14 |
| **AllR** | **GlnG EC Ref 1** | 1418 | 13.47 |
| **AllR** | **Nac EC Ref 1** | 1374 | 13.5 |
| **AllR** | **PdhR neg** | 1401 | 13.48 |
| **AllR** | **PdhR EC Ref 1** | 1566 | 13.37 |
| **AllR** | **UlaR EC Ref 1** | 1539 | 13.38 |
| **GlnG** | **AllR EC Ref 1** | 12950 | 11.25 |
| **GlnG** | **GlnG EC Ref 2** | 5.52 | 19.01 |
| **GlnG** | **GlnG EC Ref 3** | 41.38 | 17 |
| **GlnG** | **GlnG S1** | 2.79 | 19.7 |
| **GlnG** | **GlnG S11** | 29.39 | 17.34 |

| | | | |
|---|---|---|---|
| **GlnG** | **GlnG S21** | 104.3 | 16.08 |
| **GlnG** | **Nac EC Ref 1** | 11670 | 11.36 |
| **GlnG** | **PdhR EC Ref 1** | 10990 | 11.42 |
| **GlnG** | **UlaR EC Ref 1** | 6413 | 11.96 |
| **Nac** | **AllR neg** | 6.68 | 18.82 |
| **Nac** | **AllR EC Ref 1** | 4280 | 12.36 |
| **Nac** | **argP EC Ref 1** | 21.79 | 17.64 |
| **Nac** | **GlnG neg** | 3217 | 12.65 |
| **Nac** | **GlnG EC Ref 1** | 2761 | 12.8 |
| **Nac** | **Nac EC Ref 1** | 3.31 | 19.53 |
| **Nac** | **Nac EC Ref 2** | 5.78 | 18.97 |
| **Nac** | **Nac EC Ref 3** | 8.98 | 18.53 |
| **Nac** | **Nac S1 430304** | 2.49 | 19.81 |
| **Nac** | **Nac S11 364632** | 3.02 | 19.62 |
| **Nac** | **Nac S21 1022037** | 5.21 | 19.07 |
| **Nac** | **Nac S31 603054** | 20.24 | 17.72 |
| **Nac** | **PdhR neg** | 1663 | 13.31 |
| **Nac** | **PdhR EC Ref 1** | 4825 | 12.24 |
| **Nac** | **UlaR EC Ref 1** | 7432 | 11.81 |
| **PdhR** | **AllR neg** | 3721 | 12.5 |
| **PdhR** | **AllR EC Ref 1** | 190.3 | 15.47 |
| **PdhR** | **argP EC Ref 1** | 818 | 14.02 |
| **PdhR** | **GlnG neg** | 14870 | 11.12 |

| | | | |
|---|---|---|---|
| **PdhR** | **GlnG EC Ref 1** | 4484 | 12.31 |
| **PdhR** | **Nac EC Ref 1** | 207.9 | 15.39 |
| **PdhR** | **PdhR neg** | 6112 | 12.01 |
| **PdhR** | **PdhR EC Ref 1** | 3.73 | 19.41 |
| **PdhR** | **PdhR EC Ref 2** | 3.95 | 19.35 |
| **PdhR** | **PdhR EC Ref 3** | 3.5 | 19.47 |
| **PdhR** | **PdhR S1** | 1.61 | 20.24 |
| **PdhR** | **PdhR S11** | 2.41 | 19.84 |
| **PdhR** | **PdhR S27** | 3.5 | 19.47 |
| **PdhR** | **PdhR S31** | 19.34 | 17.76 |
| **PdhR** | **PdhR S41** | 17.91 | 17.84 |
| **PdhR** | **PdhR S8** | 1.35 | 20.42 |
| **PdhR** | **UlaR EC Ref 1** | 2056 | 13.09 |
| **UlaR** | **AllR neg** | 3051 | 12.7 |
| **UlaR** | **AllR EC Ref 1** | 188 | 15.49 |
| **UlaR** | **argP EC Ref 1** | 1626 | 13.33 |
| **UlaR** | **GlnG neg** | 12070 | 11.32 |
| **UlaR** | **GlnG EC Ref 1** | 83.99 | 16.29 |
| **UlaR** | **Nac EC Ref 1** | 355.9 | 14.85 |
| **UlaR** | **PdhR neg** | 12050 | 11.33 |
| **UlaR** | **PdhR EC Ref 1** | 3525 | 12.56 |
| **UlaR** | **UlaR EC Ref 1** | 18.45 | 17.81 |
| **UlaR** | **UlaR EC Ref 2** | 10.72 | 18.35 |

| | | | |
|---|---|---|---|
| **UlaR** | **UlaR EC Ref 3** | 57.54 | 16.67 |
| **UlaR** | **UlaR S1 147397** | 40.81 | 17.01 |
| **UlaR** | **UlaR S11 196037** | 48.75 | 16.84 |
| **UlaR** | **UlaR S21 502983** | 164.7 | 15.62 |
